# Unconstrained generation of synthetic antibody-antigen structures to guide machine learning methodology for real-world antibody specificity prediction

**DOI:** 10.1101/2021.07.06.451258

**Authors:** Philippe A. Robert, Rahmad Akbar, Robert Frank, Milena Pavlović, Michael Widrich, Igor Snapkov, Andrei Slabodkin, Maria Chernigovskaya, Lonneke Scheffer, Eva Smorodina, Puneet Rawat, Brij Bhushan Mehta, Mai Ha Vu, Ingvild Frøberg Mathisen, Aurél Prósz, Krzysztof Abram, Alex Olar, Enkelejda Miho, Dag Trygve Tryslew Haug, Fridtjof Lund-Johansen, Sepp Hochreiter, Ingrid Hobæk Haff, Günter Klambauer, Geir Kjetil Sandve, Victor Greiff

## Abstract

Machine learning (ML) is a key technology for accurate prediction of antibody-antigen binding. Two orthogonal problems hinder the application of ML to antibody-specificity prediction and the benchmarking thereof: The lack of a unified ML formalization of immunological antibody specificity prediction problems and the unavailability of large-scale synthetic benchmarking datasets of real-world relevance. Here, we developed the Absolut! software suite that enables parameter-based unconstrained generation of synthetic lattice-based 3D-antibody-antigen binding structures with ground-truth access to conformational paratope, epitope, and affinity. We formalized common immunological antibody specificity prediction problems as ML tasks and confirmed that for both sequence and structure-based tasks, accuracy-based rankings of ML methods trained on experimental data hold for ML methods trained on Absolut!-generated data. The Absolut! framework thus enables real-world relevant development and benchmarking of ML strategies for biotherapeutics design.

**Graphical abstract:** 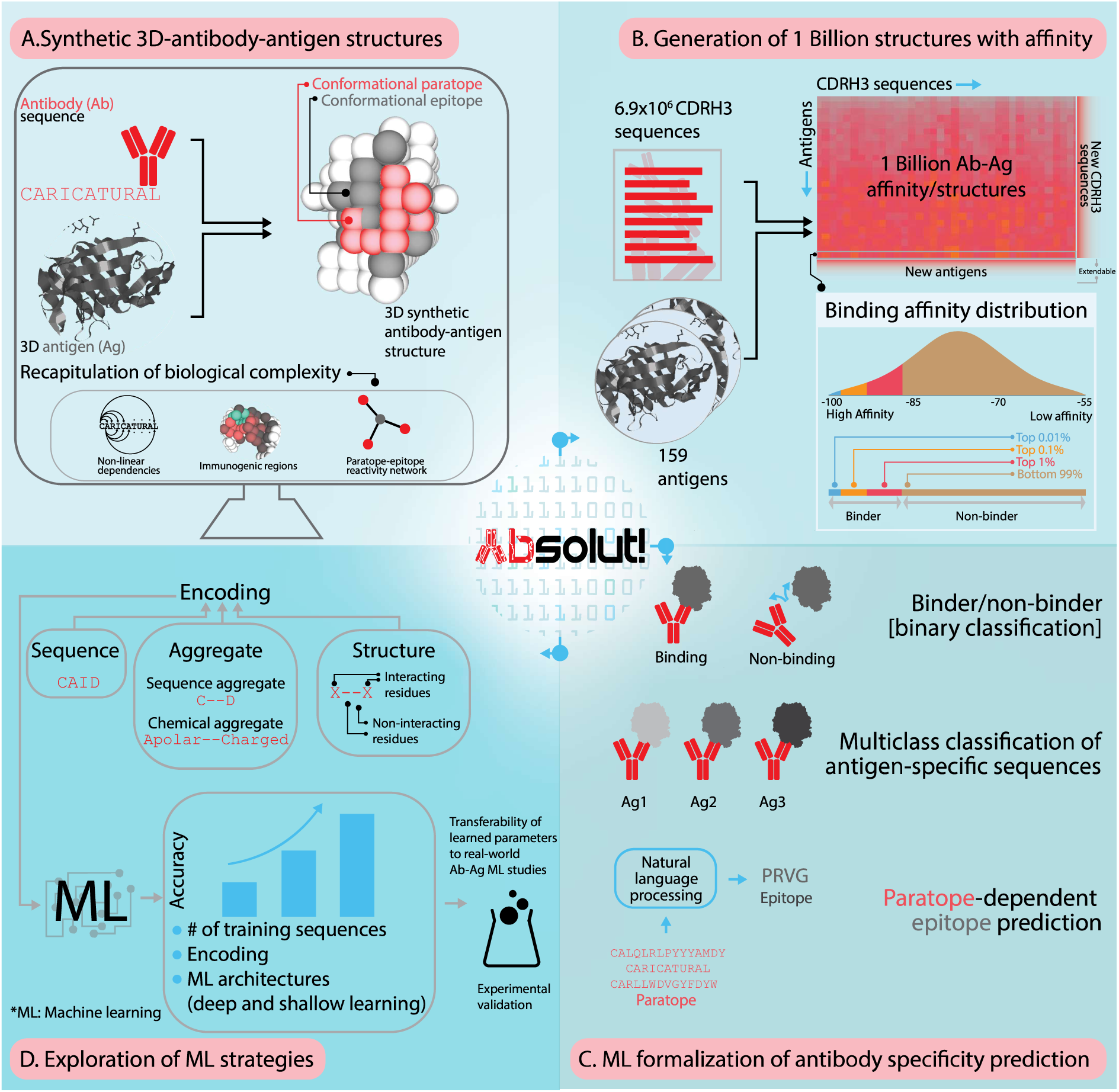
The software framework Absolut! enables (A,B) the generation of virtually arbitrarily large numbers of synthetic 3D-antibody-antigen structures, (C,D) the formalization of antibody specificity as machine learning (ML) tasks as well as the exploration of ML strategies for real-world antibody-antigen binding or paratope-epitope prediction.

**Highlights:**

- Software framework Absolut! to generate an arbitrarily large number of synthetic 3D-antibody-antigen structures that contain biological layers of antibody-antigen binding complexity that render ML predictions challenging
- Immunological antibody specificity prediction problems formalized as machine learning tasks for which the in silico complexes are immediately usable as benchmark datasets
- Exploration of machine learning prediction accuracy as a function of architecture, dataset size, choice of negatives, and sequence-structure encoding
- Relative ML performance learnt on Absolut! datasets transfers to experimental datasets

## Introduction

Antibodies bind foreign molecules (antigens) with high specificity. Antibody therapeutics have led to impressive medical breakthroughs in the treatment of infection, cancer, and autoimmunity ^1,2^. The 3D antibody-antigen binding interface is formed by the paratope on the antibody side, and the epitope on the antigen side ^3,4^. The antibody CDRH3 (complementarity determining region of the heavy chain) region contributes predominantly to the paratope ^5–7^.

The prediction of the 3D (or conformational) paratope and/or epitope for an antibody-antigen pair is crucial for addressing long-standing problems in computational antibody ^8–13^ and vaccine design ^14,15^. 3D-antibody-antigen complexes resolved at the atomic level represent the gold standard for describing antibody-antigen binding but they are time and cost-intensive to generate. Currently, there exist only ≈10^3^ non-redundant antibody-antigen structures ^16–18^, which is many orders of magnitudes smaller than the diversity of antibody sequences (>10^13^) ^19^. Furthermore, affinity values remain unavailable for the majority of both 3D-structural datasets and antigen-specific antibody sequences obtained from repertoire ^20^ or library-based screening approaches ^21^. The lack of structural antibody-antigen binding data combined with the complexity of antibody-antigen binding ^22,23^ and protein-protein docking ^24–27^ are one of the main reasons why the prediction of antibody-antigen binding remains an unresolved problem.

Machine learning (ML) is increasingly used for antibody-antigen binding prediction ^5,10,21,28–41^ given its capacity to infer the hidden nonlinear rules underlying high-complexity protein-protein interaction ^37,42–47^; with rules including long-distance dependencies between amino acids at the binding interface. Such ML methods encompass sequence-based paratope prediction ^5,7,29,48–50^ and paratope-epitope-linked prediction ^5,37,49,51^ while varying in the extent of inclusion of structural information. Structure- or sequence-based binding prediction may be feasible provided that sufficiently large antigen(epitope)-specific antibody datasets become available ^16,17,52–56^. Currently, ML applications are developed on very small or incomplete-knowledge datasets (joint information on paratope, epitope, affinity is unavailable, size of datasets usually <10 000 antibodies) ^57^. Restricted experimental datasets allow neither the benchmarking and stress testing of ML methods nor the verification whether ML conclusions generalize to other datasets.

Simulation allows the generation of synthetic complete-knowledge ground-truth datasets (i.e., datasets whose generation rules are known and therefore contain validated properties to be learned) containing desired levels of signal and noise that reflect experimental settings and biological mechanisms ^58–60^. Simulated datasets have been used in methodological development and calibration before large-scale datasets become available, to disentangle machine learning hypotheses and to prioritize the design of future experiments ^61,62^. For antibody-antigen binding prediction, simulations may help precisely and meaningfully define different real-world antibody-antigen binding problems, which requires levels of annotation that are not yet available in experimental data. Furthermore, the simulation of ground-truth complete-knowledge datasets is critical for benchmarking or ranking ML prediction strategies. Since ML encodings span sequence- to structure- to hybrid formalizations, simulated antibody-antigen data need to (i) recapitulate structural levels of complexity of experimental antibody-antigen binding (especially for defining paratopes and epitopes); (ii) enable the generation of large-scale datasets; and (iii) allow the integration of sequence and structural information into hybrid encodings.

Here, we provide a deterministic 3D-antibody-antigen binding simulation framework to enable ML method development and formalization on parametrized, large-scale datasets. Synthetic antibody-antigen structures are generated as the energetically optimal binding structure in a 3D lattice and recapitulate many levels of complexity inherent to antibody-antigen binding physiology, and allow for the exploration of various types of dataset designs that are largely unfeasible to generate experimentally. Specifically, we generated *synthetic* binding structures of 6.9 million murine CDRH3 sequences to 159 antigens (≈1B antibody-antigen binding pairs). Our work is based on the premise that a successful ML antibody binding strategy for experimental datasets should also perform well on synthetic datasets (and vice-versa). Therefore, the synthetic datasets should be complex enough such that accuracy-based ML method rankings can be transferred from synthetic to experimental datasets. For three use cases, we investigated the extent to which 1D-sequence and 3D-structural information is required for achieving high prediction accuracy of antibody-antigen binding: (i,ii) binary and multi-class classification of binding, and (iii) paratope-epitope prediction ^8,9^. We found that *in silico* investigated conditions predicted to increase antibody specificity prediction reflect ML performance on experimental antibody-antigen sequential and structural data.

## Results

### Formalizing and benchmarking antibody specificity prediction problems as ML tasks requires simulated 3D-antibody-antigen data

Predicting antibody specificity refers to identifying which antibody sequence(s) or structure(s) bind to which antigen(s), and vice versa (Figure 1A), and remains an unresolved challenge ^10^. There exists a large variety of possible biological and computational problem formulations and assumptions (Supplementary Figure 1), and a formal framework that enables a unified formulation of antibody-antigen binding tasks is lacking.

**Figure 1.**
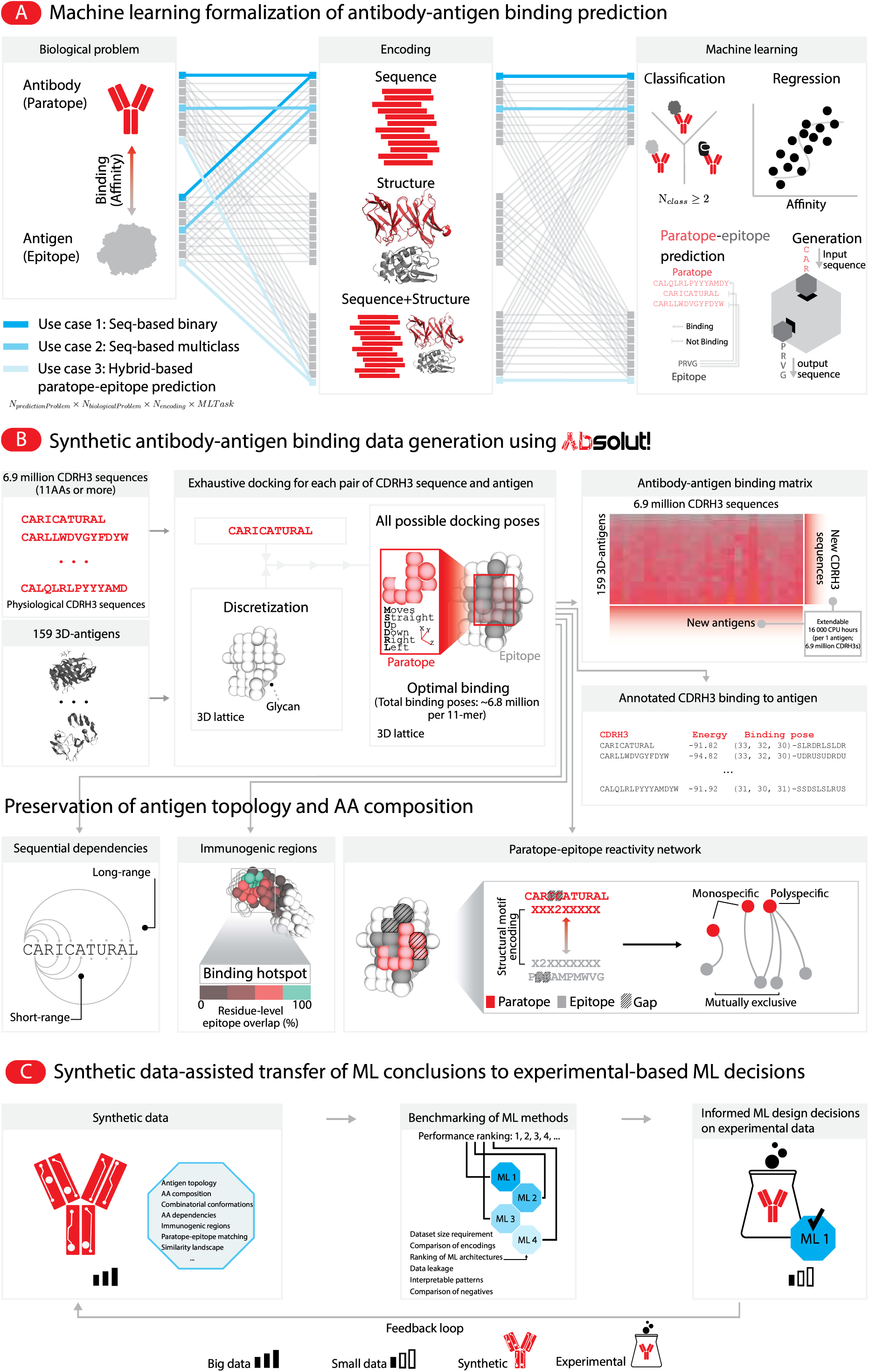
Machine-learning formalization of antibody-antigen binding prediction tasks and pipeline for the high-throughput generation of 3D antibody-antigen structure datasets. (A) Machine-learning formalization of the biological problem of antibody-antigen binding prediction. The biological formulation of the problem may involve either antibody or antigen, subsets thereof (paratope, epitope), affinity, sequence or structure or any combination thereof. The choice of data encoding can be divided into sequence-based and structure-based ones, while hybrid formalizations leverage both types of datasets (Supplementary Figure 1). On the ML side, predicting whether an antibody sequence binds to an antigen may be broadly grouped into binary, multiclass or multilabel classification and regression. Regression may be used to predict the affinity of an antibody sequence to a target antigen. Furthermore, specific problems of antibody-antigen binding may be, for instance, predicting which residues of an antibody or an antigen are involved in the binding (paratope and epitope prediction, and generation of new antibody or antigen sequences). (B) Synthetic antibody-antigen binding data generation pipeline using the Absolut! framework: the PDB structure of an antigen is transformed into a 3D lattice representation largely preserving antigen topology and surface amino acid composition (Supplementary Figure 7). Datasets can be generated with unconstrained size. CDRH3 sequences are then tested for binding using exhaustive docking (6.8 million binding poses per each 11-mer of each CDRH3 sequence and antigen, Supplementary Figure 2) to identify the energetically optimal binding structure. The presence of glycans on the protein is parsed from the original PDB and may be included in the lattice representation. For each optimal antigen-binding structure, the 3D paratope, 3D-epitope and affinity values are obtained and recorded. By screening 6.9 million experimental CDRH3 sequences of length 11 or more (therefore containing physiological amino acid composition and dependencies) ^63^, we thus produced a database (antibody-antigen binding matrix) of 1 billion antibody-antigen binding pairs with 3D paratope+epitope+affinity resolution. The generated dataset encompasses the following levels of biological complexity of antibody-antigen binding: 1) Antigen topology; 2) antigen AA composition; 3) physiological CDRH3 sequences; 4) a combinatorially large amount of millions of possible binding conformations sampled during exhaustive docking; 5) positional amino acid dependencies in high-affinity sequences; 6) immunogenic regions (binding hotspots, showing clusters of epitopes that share residues. Turquoise residues are contained in all epitopes binding this region, while levels of red show how often a residue is contained in an epitope); 7) a complex paratope-epitope matching landscape (that can be visualized on reactivity network showing the organization of binding pairs according to a specific encoding and can be compared to atomistic experimental 3D-antibody-antigen binding data); and 8) a “broken similarity” binding landscape, where similar antibodies do not necessarily bind the same antigens. (C) Synthetic antibody-antigen binding data with native-like levels of complexity is necessary for understanding the relative performance of dataset design and ML benchmarking of antibody specificity prediction methods. In this manuscript, we show that ML insights (performance ranking of ML methods) informed by ML application to *synthetic* antibody-antigen data, transfer to *experimental* antibody-antigen binding data.

Most ML formalized tasks fall into three main categories: (i) Classification: A binary classification predicts whether an antibody binds to a predefined antigen or not (Figure 1A). An antibody may further be separated in more classes describing the binding to one antigen (for example low-affinity, medium-affinity, high-affinity classes); the binding to different epitopes on an antigen; or even the binding to different antigens (Figure 1A), defining a multiclass (or multilabel) classification problem if an antibody can only belong to one class (or to multiple classes). (ii) Regression: Prediction of an antibody sequence affinity to a target antigen, or sequence developability parameter values. (iii) Paratope-epitope prediction: Prediction of which residues of an antibody-antigen complex are involved in their binding interface (paratope or epitope prediction, Figure 1A), or prediction of the matching between a paratope and an epitope, possibly in an encoded form. Of note, beyond only encoding antibody and antigen sequences, all these ML tasks may involve the reconstruction of complex data structures describing features of the binding interface, that can be leveraged e.g., by Natural Language Processing (NLP) architectures. All aforementioned tasks may also be formulated starting from the antigen perspective and predict their binding to predefined antibodies (bidirectionality of antibody-antigen binding prediction).

Each ML problem formulation requires different types of dataset structures and encodings, rendering the comparison of their effectiveness challenging. Large and reproducible synthetic datasets can be parametrized into specific immunology problems. ML-task-adapted training datasets can enable the relative ranking of ML strategies for antibody-antigen prediction problems, following the hypothesis that architectures or ML strategies that perform better on synthetic datasets are expected to also perform better on experimental datasets (Figure 1C).

### Unconstrained generation of *in silico* antibody-antigen structures

We present the Absolut! framework that enables deterministic generation of large synthetic datasets of 3D-antibody-antigen binding at moderate computing costs. Absolut! simulates the binding of antibody sequences (CDRH3) to antigens (from PDB) *in silico* (Figure 1B) using a lattice representation of protein interactions (Methods, Supplementary Figure 2A).

First, Absolut! creates a discretized lattice representation of the protein antigen (Figure 1B), by minimizing the dRMSD between the original PDB structure and its many possible lattice counterparts ^64^ (Supplementary Figure 3A, Methods). Protein glycosylation is modeled as “inaccessible positions” on the surface of the discretized antigen (Supplementary Figure 3A,B), which impact the binding affinity of the CDRH3 sequences (Supplementary Figure 4G), as observed experimentally ^65,66^. We optimized the lattice resolution (distance between neighboring amino acids) to reach an average RMSD of 3.5Å to the original PDB (Supplementary Figure 3C–F). In brief, the protein antigen discretization step preserves realistic 3D antigen sizes (Supplementary Figure 4B), shapes, and surface amino acid composition (Supplementary Figure 7A).

Second, Absolut! enables the calculation of the energetically optimal binding of a CDRH3 sequence to a lattice-discretized antigen. Briefly, Absolut! enumerates all possible binding poses of the CDRH3 to the antigen ^67^, computes their binding energy (scoring function) using the Miyazawa-Jernigan energy potential ^68^, and returns the best pose as the “binding structure” of the CDRH3, a step we termed “exhaustive docking” (Figure 1B, Supplementary Figure 2B, Methods). Each binding structure provides 3D-information on paratope, epitope, and affinity. The advantage of exhaustive docking is twofold: it screens the entire lattice epitope space and ensures that the energetically global optimal binding is always found, in contrast to docking of experimental or modeled structures ^69^.

### One billion Absolut!-generated *in silico* antibody-antigen structures as a basis for ML benchmarking and ML antibody-antigen binding task formulation

We generated a library of synthetic 3D-antigens (including pathogenic and self-antigens, see Table S1) of 159 antigens from crystallized antibody-antigen complexes ^16^, and further calculated the binding structures of a database of 6.9 million unique CDRH3 antibody sequences obtained from murine naïve B-cells ^63^ to the 159 discretized antigens (Figure 1B, Supplementary Figure 4A-C). Using experimental CDRH3 sequences ensures that antibody sequences possess, analogously to the antigen, physiological amino acid composition and positional dependencies. The CDRH3 sequences were assessed for binding by sub-peptides of size 11 amino acids for the purpose of computational tractability and consistency (Figure 1B, “exhaustive docking” in Methods). The generated database contains 1.1 billion antibody-antigen binding structures with conformational paratope, conformational epitope and affinity resolution (Figure 1B). Human CDRH3 ^70^ may be used instead of murine ones, and showed similar binding behavior in terms of affinity distribution and epitope convergence (Supplementary Figure 4H,I). Each 11-mer of a CDRH3 sequence required on average the enumeration of 6.8 millions binding poses per antigen by exhaustive docking (Supplementary Figure 4D), depending on the antigen size (Supplementary Figure 4E), and required 7.9 seconds per CDRH3 on average (Supplementary Figure 4F). This is a fairly moderate computational requirement in view of the 6.9 million CDRH3 times 6.8 million docking poses per antigen. Therefore, the Absolut! antibody-antigen binding dataset is not only ultra-large but also easily extendable.

Importantly, for ML analyses, Absolut!-generated data enables the extraction of various features describing the paratope-epitope interface (Supplementary Figure 5), and mirrors the diversity of used encodings in ML studies, from binary vectors ^29^ to 3D distance representations ^71,72^. Altogether, the Absolut! framework and the generated database of *in silico* antibody-antigen binding complexes enable the unconstrained generation of antibody-antigen datasets designed specifically for different ML problems.

### Absolut!-generated datasets reflect multiple levels of biological antibody-antigen binding complexity

A prerequisite for the real-world-relevant comparison of ML strategies (architecture, dataset design, encodings) for antibody-antigen binding prediction is that Absolut! datasets reflect as many levels of biological antibody-antigen complexity as possible.

In the Absolut! framework, antibody sequences bind to an antigen with a large diversity of binding energies (Figure 2A) and structures, as shown as an example, for the antigen with pdb-id 1ADQ_A (Figure 2B). We arbitrarily define “binders” to an antigen as those (CDR3–derived) 11-mer sequences within the top 1% affinity threshold (1% lowest binding energies, Figure 2B, Supplementary Figure 9A, see Methods). Grouping CDRH3 sequences (or 11-mers) by affinity classes of arbitrarily adjustable thresholds allows exploring different definitions of negative (non-binder) samples in ML tasks. Among the 159 antigens, the top 1% binders used on average 62 different binding structures per antigen, with on average 17 distinct epitopes (Supplementary Figure 4J–L), reflecting experimentally observed antibody binding diversity ^73^.

**Figure 2.**
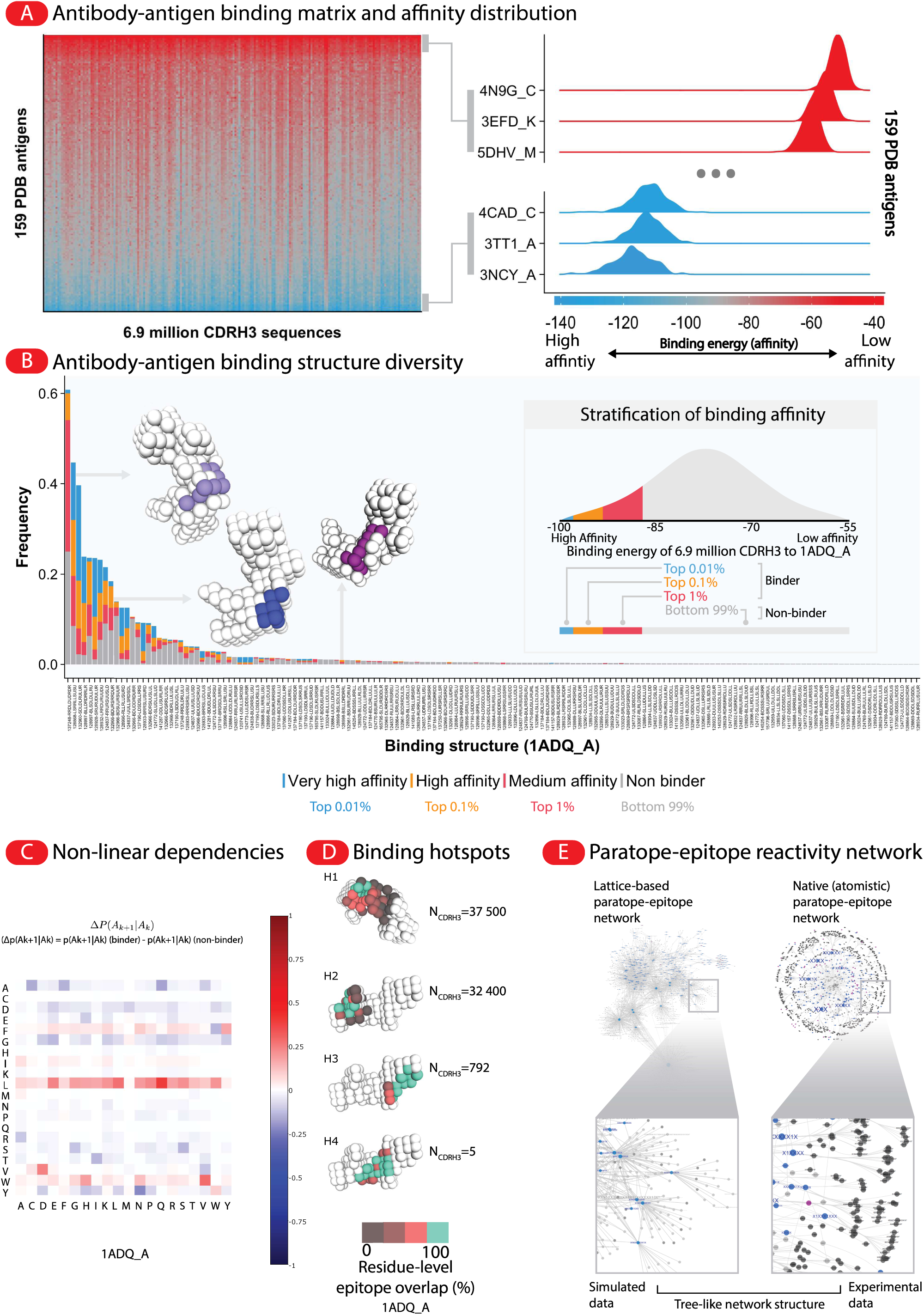
The Absolut! dataset reflects granular levels of the biological complexity of antibody-antigen binding. (A) Affinity of the 1 113 000 000 antibody-antigen pairs in the Absolut! database, shown as an affinity matrix of 159 antigens and 6.9 million CDRH3 sequences (left panel). Each heatmap tile represents the binding energy (affinity) of a CDRH3 sequence against an antigen. The right panel illustrates the affinity distributions of the top and bottom three antigens in terms of median affinity. (B) Diversity of binding structures in the dataset for antigen 1ADQ_A (see Methods). [Inset] Affinity annotation in (A) allows custom-stratification of CDRH3 sequences into binders and substratification thereof (the top 1% of affinity sorted CDRH3 sequences for each antigen separately) and non-binders (the bottom 99% of affinity sorted CDRH3 sequences). [Main panel] Distribution of 3D-binding structures with respect to each binding class as defined in the inset. The x-axis represents the antibody binding structures to this antigen, written as a 6-digit number representing a starting 3D position in the lattice, followed by a list of moves in 3D-space (see Supplementary Figure 3A or *Methods* for details). Different CDRH3 sequences may converge to the same binding structure albeit with different affinities (i.e., one bar with 4 colors reflects a single binding structure containing all four binding classes). An affinity class could be distributed across multiple binding structures (i.e., blue bars: very high-affinity sequences, are found across multiple structures in the x-axis). Only 159 of the total 842 identified binding structures are shown for legibility, only 65 of them are used by the top 1% binding sequences. (C) Positional dependencies are illustrated by comparing the observed conditional probabilities of an amino acid A_k+1_ at position k+1, knowing the amino acid A_k_ at the previous position (p(A_k+1_|A_k_)), between binders and non-binders (Δp(A_k+1_|A_k_) = p(A_k+1_|A_k_) (binder) - p(A_k+1_|A_k_) (non-binder). Positional dependencies at longer range, are shown in Supplementary Figure 6 for amino acids within a distance up to 5 and 11 amino acids, instead of only 2 amino acids here). Red squares denote a case where two consecutive amino acids are more highly dependent in binders than in non-binders, while blue squares show more negatively dependent consecutive amino acids in binders than in non-binders. (D) Binding hotspots identified for antigen 1ADQ_A, named H1 to H4 (see Methods and Supplementary Figure 22). (E) The paratope-epitope binding network (reactivity network) constructed with the Absolut! dataset (left) shows a complex topology with oligo- and polyspecific characteristics also found in experimentally determined reactivity networks ^5,16^. Different paratope-epitope encodings determine different reactivity networks. The network shows gapped structural interaction motifs, for instance, epitope X1X2X3XX82X3XXXXXXX and paratope XXXX1XX1XXX where X denotes an interacting residue and the numbers refer to the number of non-interacting residues in-between (see Methods and ^5^). The Absolut!-generated network contains 6092 unique paratope-epitope pairs, with 2572 unique epitopes and 324 unique paratopes with this encoding. Epitopes are shown in blue and purple, and paratopes appear in light or dark gray. Specific epitopes (purple) are defined as being bound by only one paratope in the dataset (all binder sequences use this paratope-epitope pair), while specific paratopes (light gray) are only bound by one epitope. The reactivity network degree distributions of Absolut! and experimental immune (antibody-antigen) and non-immune (protein-protein) datasets using the most predictive encoding from Figure 5A are compared in Supplementary Figure 8.

Analysis of binder sequences to an antigen (1ADQ_A) showed non-linear positional dependencies ^74^ of amino acids as compared to non-binder sequences. (Figure 2C, Supplementary Figure 6) ^5,39^. Clustering the binding sequences to 20 different antigens showed that very similar antibody sequences recognized different antigens (Supplementary Figure 14D), which is a major current challenge for ML prediction tasks applied to experimental antibody-antigen binding ^34,39,75^ (Supplementary Figure 14E-H).

To quantitatively describe preferential CDRH3 binding to specific antigen regions, we developed an algorithm to cluster the modes of antibody binding (epitopes of binder sequences) into, hereafter called, “binding hotspots” (see Methods, Supplementary Figure 22, Figure 2D for 1ADQ_A), mirroring the concept of immunogenic regions of an antigen ^76^. Each cluster is defined by a core set of at least four shared epitope positions between different epitopes (turquoise), while other residues are colored with levels of red representing the fraction of epitopes (of the cluster) containing the residue, and white areas denote antigenic regions that are more challenging to bind with high affinity, as no predicted binding from the top 1% binders had its epitope in these regions. The number of binding sequences to each hotspot ranged from 5 to 37 500 CDRH3 sequences for the antigen 1ADQ_A (Figure 2D), suggesting the Absolut! database mirrors the experimentally observed hierarchy of more or less immunogenic domains within the same antigen ^77^.

Although Absolut! is neither suited nor designed to directly predict where an antibody sequence would bind in the real world, we quantified whether antigen regions predicted to be binding hotspots overlap with known binding sites in experimental crystal structures. In 75% of examined experimental antibody-antigen structures from which the antigens were discretized, the experimental paratope was overlapping with a binding hotspot core residue on the discretized antigen, while this number reached 85% when including binding hotspot side residues (Supplementary Figure 7B-E). Since for this comparison full chain experimental antibody structures were compared with CDRH3-based Absolut! structures, the overlap calculated represents an upper bound. This suggests that topological factors that make antigen regions immunogenic are somewhat preserved in Absolut! datasets.

ML tasks are often set up to predict which antibody sequence binds to which antigen, according to specific encodings. The reactivity network of Absolut! links 53 000 binder sequences on average to each antigen. Here, we compare the reactivity networks of Absolut! (Figure 2E, left) and experimentally determined binders (Figure 2E, right, generated from 825 experimentally-determined antibody-antigen pairs ^5^) according to such encoding. In both synthetic and experimental data, we observed a complex organization (tree-like structure) and mutually exclusive polyreactivity of certain binding modes (motifs). Properties of the synthetic reactivity network are statistically more similar to the experimental antibody reactivity network than to the protein-protein interaction network (Supplementary Figure 8).

Taken all together, although simplifying antibody-antigen binding, Absolut! encompasses eight substantial levels of structure- and sequence-based complexity levels (Figure 1, Figure 2), a subset of which are encountered in experimental antibody-antigen binding highlighting the relevance of Absolut!-generated data for developing and benchmarking antibody-antigen binding prediction methods.

### Absolut! allows the prospective evaluation of ML strategies with custom-designed datasets

To demonstrate Absolut!’s usefulness in assessing the efficacy of different binary and multi-class classification ML strategies (Figure 1A), we compiled three datasets (Figure 3A). Binary classification datasets D1 and D2 were generated for each antigen separately. D1 was generated with 50% binders of top 1% affinity (blue) and 50% non-binders containing both top 1–5% affinity (red) and bottom 95% affinity (gray), while in D2 the non-binders are instead only defined by top 1–5% affinity (Figure 3A, Methods, Supplementary Figure 9A). Our selected multi-class classification example aims at identifying the target (class) of an antibody sequence among N antigens (class) excluding cross-reactive sequences (one label per sequence only). The multiclass dataset 3 (D3) was generated for a set of 5 to 140 randomly chosen antigens, by pooling the top 1% binder sequences to each antigen (Figure 3A), while discarding those sequences that bind more than one antigen. The size of D3 depends on the number of selected antigens and is imbalanced (Supplementary Figure 9B,D,E). As a comparison, including cross-reactive sequences in D3 would reach up to 1 250 000 sequences to all antigens (Supplementary Figure 9C) and also allows evaluating ML performance on the multilabel problem (Figure 1, Supplementary Figure 12).

**Figure 3.**
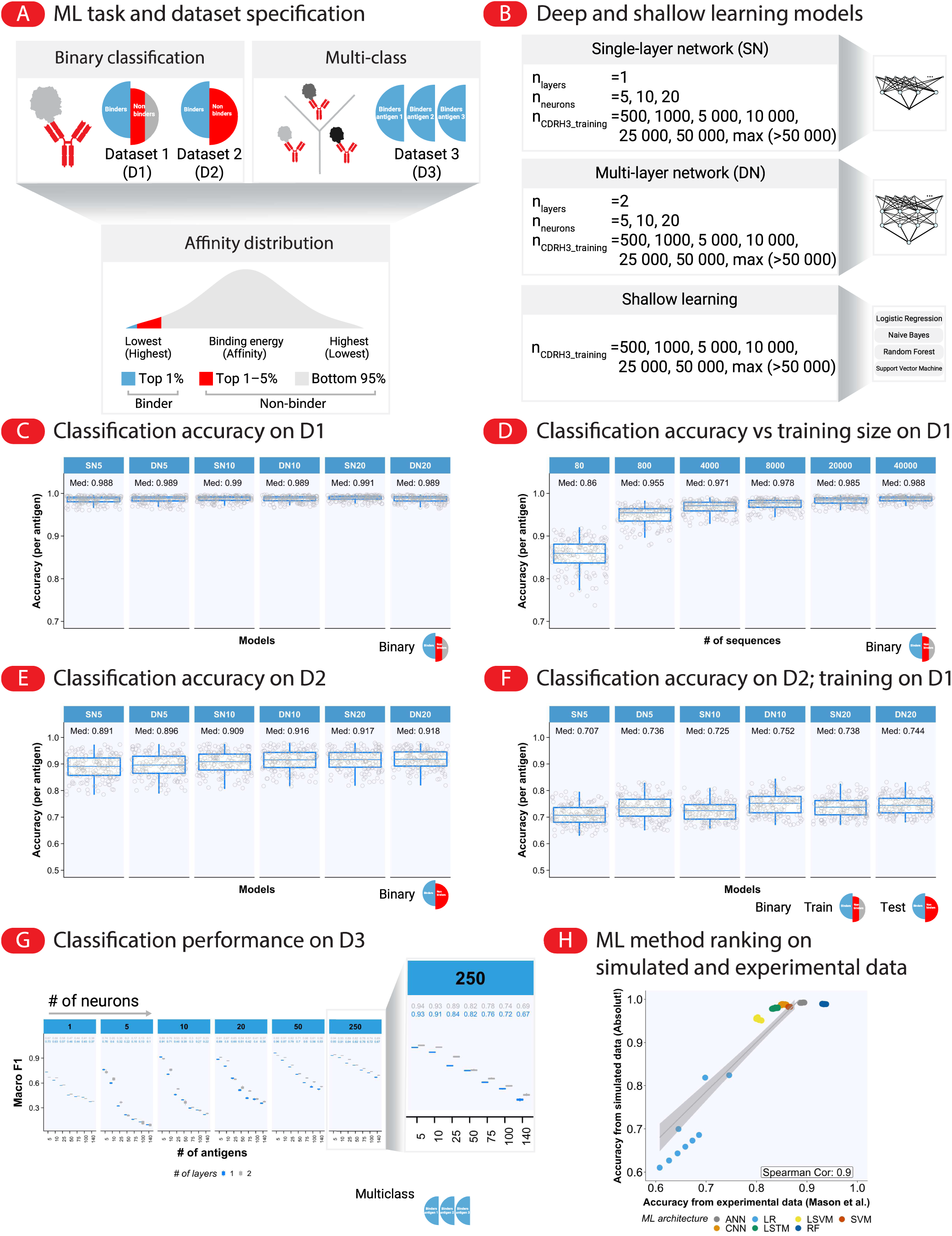
Classification of binding and non-binding antibody sequences with machine learning. (A) Design of datasets for the ML task of classifying a sequence as binding in a binary (binding, non-binding) or multi-class (binding to one antigen among a set of N antigens) setting. Binding is assessed for sub-sequences of size 11 amino acids derived from CDRH3 sequences (see Methods Supplementary Figure 9). For each antigen separately, we defined binders as 11-mer sequences within the top 1% affinity threshold (blue), non-binders as 11-mer from the top 1–5% (red), and the bottom 95% (gray). Datasets 1 (D1) and 2 (D2) have been created for binary classification for each antigen separately, and D3 for multi-class classification for a set of n_antigens_ antigens (Supplementary Figure 9). D1 was generated with 50% binders of top 1% affinity (blue) and 50% non-binders containing both top 1–5% affinity (red) and bottom 95% affinity (gray), while in D2 the non-binders are defined only by top 1–5% affinity (excluding the bottom 95%), rendering this dataset a priori more challenging to classify since binders and non-binders are closer affinity-wise. In both datasets, binders and non-binders were sampled as 11-mers taken from CDRH3 sequences of the same length distribution, to limit potential amino acid composition bias (see Methods). D3 was generated by pooling sequences that were among the top 1% binders across multiple antigens, labeled with the name of their antigen, discarding 11-mers that bound multiple antigens to fit a multi-class classification problem. We also discarded antigens representing variants of the same protein, leaving 142 non-redundant antigens, i.e., different labels (see Table S1 for the list of discarded antigens). Due to the removal of cross-reactive sequences, D3 is imbalanced, because the amount of binder sequences to an antigen that was cross-reactive (with another selected antigen) were highly variable, as seen by the number of available sequences for D3 (Supplementary Figure 9C). (B) Model architectures considered: Feed-forward neural network with a single layer (SN), feed-forward neural network with deep (two-layers) (DN) of equal number of neurons (5–20 neurons; in this Figure, SN5 indicates a single-layer network with 5 neurons and DN5 indicates a DN with 5 neurons), and four shallow learning methods namely logistic regression (LR), Naive Bayes (NB), random forest (RF), and support vector machine (SVM) (see Supplementary Figure 10 and Supplementary Figure 11). The ML models were trained on the two tasks: binary classification (binder or non-binder) on D1 and D2 for each antigen separately, and multi-class classification (prediction of an antibody’s target out of n_antigens_ targets) on D3. n_training_=80 to 40 000 sequences were used for D1 and D2 datasets as training dataset while n_testing_=40 000 sequences were always used for testing, to avoid a bias of test size. For D3, n_training_=200 000 sequences were used for training and n_testing_=100 000 sequences for testing (or scaled-down when less than 300 000 sequences were available, see Methods for details). (C–F) Accuracy of binary classification prediction of the considered model architectures per antigen. One datapoint represents the accuracy to the dataset (D1 or D2) of one antigen (each point is the average of 10 independent replicates, 159 points in total). Median accuracies among the antigens are shown in each panel. Prediction accuracies of shallow architectures and conditions are shown in Supplementary Figure 10, including a control with shuffled labels (i.e., where the causal link between paratopes and epitopes was removed) to quantify prediction accuracy on randomized data. (C) All models, trained on 40 000 sequences from D1 irrespective of their network architectures, yielded accurate predictions for the binary classification task with median accuracy values ranging between 0.988–0.991 (see Supplementary Figure 10 for shallow learning). (D) Prediction accuracy improved as a function of the number of training sequences (here shown for the DN10 architecture). The model trained on the least number of sequences (n_training_=80) yielded the lowest accuracy (median 0.86) whereas the model trained on the largest dataset (n_training_=40 000) yielded the highest accuracy (median 0.988). (E) Prediction accuracy (trained on 40 000 sequences) was slightly lower on D2 (compared to D1 in (C)) with median accuracy values ranging between 0.891–0.918. (F) Models trained on D1 and tested on D2 yielded notably lower performance with median accuracy values ranging between 0.704–0.744. (G) Multi-class prediction performance improved as a function of model complexity (number of neurons and number of layers) and decreased as a function of the number of antigens (classes). We quantified the accuracy as macro F1 score ^78^ due to the class imbalance of D3 10 independent replicates are shown as boxplot for each condition and show very little variation. Macro-averaging weights each class equally (see Methods). Specifically, the median macro F1 values for the model with the least number of classes (n_antigens_=5) ranged between 0.76–0.962 whereas the model with the most number of classes (n_antigens_=140) yielded macro F1 values between 0.097–0.67, depending on the number of neurons and layers. Baseline shuffled (expected) median macro F1 values ranged between 0–0.02 and 0–0.03 (Supplementary Figure 11). (H) Comparison of the relative performance of ML architectures on a sequence binary classification problem when they are trained and tested either on an experimental dataset ^39^ containing 7 894 binding sequences and 17 237 non-binders to HER2, or on Absolut! D1 where the same amounts of binder and non-binder sequences were taken for 10 randomly selected antigens independently. CDRH3 sequences were one-hot encoded as 1D or 2D vectors depending on the architecture (see Table 3). For each ML method (color), eight “balancing conditions” (data points) were investigated where the relative amounts of binder and non-binder in the training was varied, as in Mason et al. (see Methods and Supplementary Figure 24). One datapoint represents one ML method and one condition, and is the average of 10 independent simulations for this condition (each time with a newly sampled set of binders and non-binders from Mason dataset, and each time using the D1 from a different antigen for Absolut! dataset). The x-axis shows the performance of an ML method and balancing condition on a Mason processed dataset, while the y axis shows the performance of the same ML method and condition on the Absolut! processed dataset. As a general trend, the ML methods that perform better on the Absolut! dataset also perform better on the Mason (experimental) dataset (Spearman correlation of 0.9 between ML-based accuracies on Absolut! and the Mason dataset using identical ML architectures in both datasets).

We compared a set of ML methods with respect to binary and multi-class classification, namely Logistic Regression (LR), Random Forest (RF), Naive Bayes (NB), Support Vector Machine (SVM) a single layer neural network (SN*n*), as well as a deep (two-layer) network (DN*n*) (Figure 3B) where *n* stands for the number of neurons in each hidden layer. For brevity, we showed the results for SN and DN in Figure 3C–G and the remainder (LR, RF, NB, and SVM together with amino acid composition and shuffling controls for SN) in Supplementary Figure 10. Nota bene, in this manuscript, we do not aim to optimize the architectures and hyperparameters of the machine learning models (Figure 3, Figure 4, Figure 5). The ML investigations shown merely showcase the utility of Absolut! in creating custom-designed datasets with varying specifications that can be used to explore and train a wide range of ML methods for antibody specificity research.

**Figure 4.**
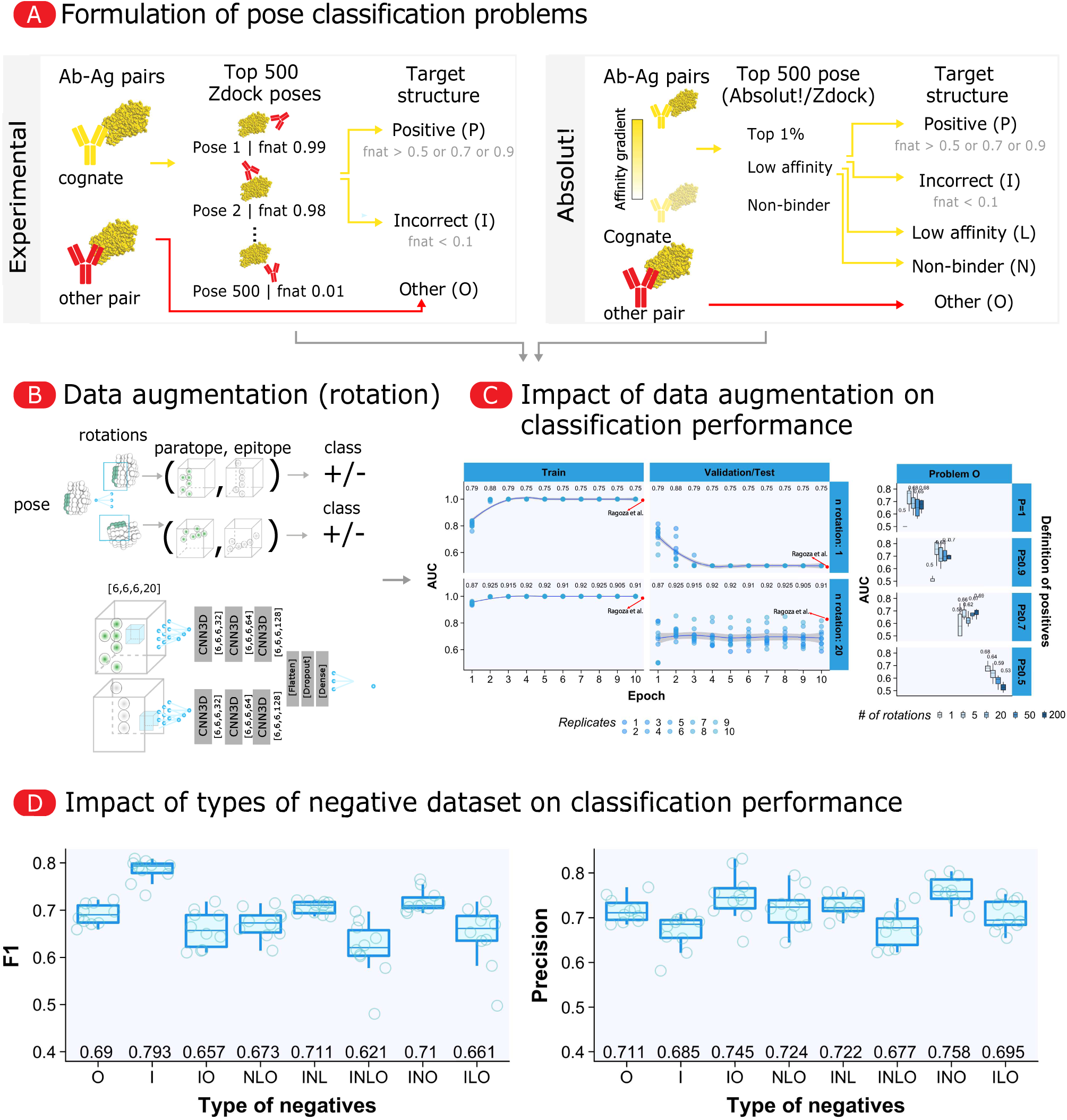
Transferability of ML method rankings in the context of pose classification and impact of type of non-binding pose (negative example) in the training datasets. (A) Pose classification problem formulation and data generation pipeline. Starting from an antibody-antigen pair, an antibody-antigen pose proposed by docking was classified as positive if it is similar to an experimental known binding structure for this antibody-antigen pair (the “target binding structure”). If the pair was not binding, there is no target binding structure and all poses should be classified as negative. If the pair was binding with high affinity, only poses similar to the target structures should be classified positive. The similarity between a pose and the target pose was quantified using the “fnat” score (fraction of conserved native contacts), which calculates the relative amount of shared interacting residue pairs between two poses. The fnat ranges between 0 (no shared interaction) and 1 (all interactions of the target pose are recapitulated). We created an *in silico* pipeline to generate docking poses for the pose classification problem, reflecting the DLAB-VS pipeline used on experimental data by Schneider et al. ^81^. In DLAB-VS (left panel), the antibody structure is modeled and docked to the given antigen structure, and a positive pose (P) is a pose from a high affinity binding antibody-antigen pair that has a fnat score over a certain threshold (for instance, 0.7). Two types of poses were defined as negative examples: those from a cognate pair with fnat lower than 0.1 (i.e., very dissimilar to the target binding structure), called “incorrect” (I), and any pose from a non-cognate antibody-antigen pair created by taking a binding antibody sequence to an antigen and pairing it with another antigen, called “other pair” (O). DLAB-VS datasets can therefore contain P, O and I poses. In the *in silico* pipeline (right panel), the experimental binding pose of a binding pair is replaced by the energetically optimal pose calculated from exhaustive docking, as the target pose. For any antibody-antigen pair, we used Absolut! to output the 500 energetically best poses among the millions of poses enumerated during exhaustive docking (6.8 millions on average, Supplementary Figure 2B). Additionally to high-affinity pairs (top 1% binders, defining “P” and “I”) and other pairs created from top 1% binders to other antigens, we defined two new types of negative examples: poses from low affinity antibody-antigen pairs (L), from the top 1%–5% high-affinity sequences to an antigen, and poses from non-binding pairs (N), from the bottom 95% affinity sequences. Indeed, the “O” negatives have the bias to already bind another antigen, and might thereby have a non-random AA composition or pattern, while the “N” are only selected based on their non-binding affinity to the considered antigen. For 10 different antigens, 10 000 antibody sequences were randomly selected per antigen (among which, 500 high-affinity sequences (to generate “P” and “I”), 2500 low-affinity sequences (L), 2500 non-binders (N) and 5500 sequences with known high affinity to another antigen among the library (O), see Methods). 500 poses were generated using Absolut! for each of the 100 000 pairs, creating a dataset of 50 million poses. (B) ML formalization from DLAB-VS ^81^: a pose is encoded as two cubic lattices^81^ of size 6×6×6 amino acids, containing the spatial organization of its paratope and epitope residues, respectively. To compensate for the lack of rotational invariance of CNNs, each pose is repeated multiple times according to a different, randomly sampled, rotation. A two-track ML architecture takes the two cubic lattices and returns “1” if the pose is of type “P”, “0” otherwise. Two 3D CNNs composed of three layers process in parallel the lattice representation of the paratope and epitope residues of a pose. Their output is flattened and converted into a binary prediction using a dense layer. (C,D) Comparison of ML strategies related to data augmentation and the definition of positives and negatives. (C) Data augmentation by rotation can reduce overfitting. Comparison of prediction accuracy when each pose is randomly rotated 1, 5, 20, 50 or 200 times during training, and depending on the fnat threshold defining positive poses. The AUC resulting from 1 and 20 rotations with threshold 1 are compared to AUC observed on experimental protein-protein binding structures in Ragoza et al. ^82^. Data augmentation by rotation is not beneficial using a large definition for positive poses (fnat > 0.5). (D) The choice of negatives in the training dataset influences the pose classification prediction efficacy. The x-axis shows conditions with different negative examples included in the training dataset. The training dataset was balanced by repeating the binding poses (positive examples) to have 50% “P” and 50% of all combined negatives during training. All poses from an antibody-antigen pair were segregated either all in the training or test dataset but not both to prevent data leakage. 8 000 poses were taken for training before data augmentation by rotations (n_Rotations_ = 20). Since each condition defines a different ML problem, the trained models were evaluated on test datasets designed with the same type of negative poses, consisting of 1000 “P” and 1000 “O” pairs before rotations. This represents a form of out-of-distribution testing as the distribution of negative poses was different in the training dataset of each condition, while the same test dataset was used to compare these conditions.

**Figure 5.**
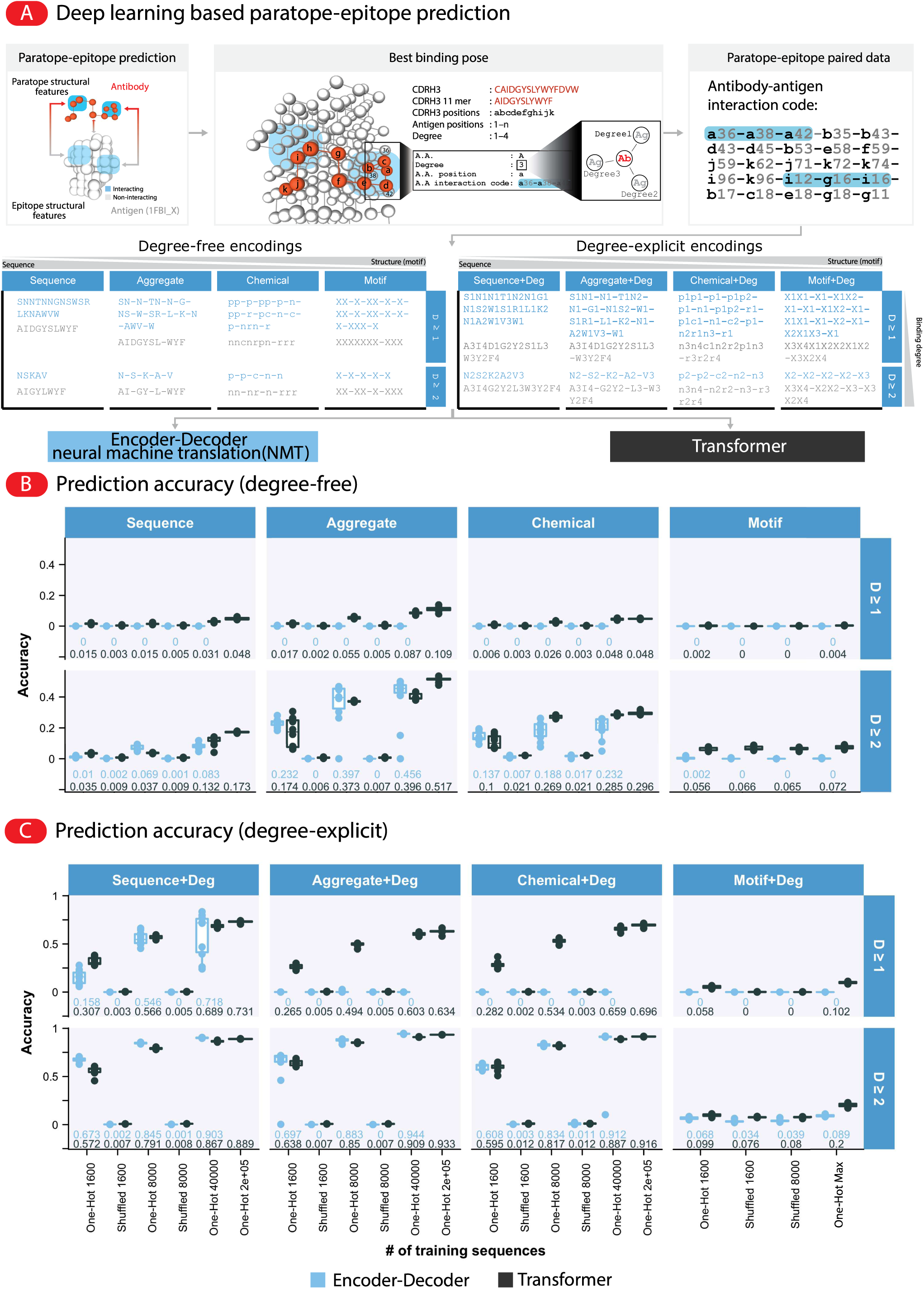
ML prediction of paratope-epitope pairs involved in antibody-antigen binding. (A) Problem formulation of paratope-epitope prediction: using a range of encodings of paratope and epitope pairs, we aim to predict the epitope encoding from the paratope encoding. From an Absolut! antibody-antigen binding complex (the best binding pose), possible elements that can be incorporated into a paratope-epitope encoding are shown on one example binding (of CDRH3 sequence CAIDGYSLYWYFDVW to antigen 1FBI_X). The interaction code stores which residues on the CDRH3 (denoted by their position as a letter from a (position 0) to k (position 11)) bind to which residues on the antigen (numbers). From the interaction code, a list of encodings is proposed to describe both sequence and structural information on the paratope and epitope of a binding. Encodings contain the list of binding residues (i.e., paratope or epitope residues), possibly separated by gaps (stretches of non-interacting residues, noted as “-”). Residues can be encoded as their name (“Sequence” without gaps or “Aggregate”, when using gaps), their chemical property (“Chemical”, with p: polar, r: aromatic, n: non-polar, c: charged) or structural interaction motif where interacting residues are encoded as the character “X”. Compared encodings can contain (degree-explicit) or not (degree-free) information on the degree of binding of each residue. Of note, the sequence encoding implicitly contains structural information, as non-binding residues are not included. We leveraged common natural language processing (NLP) architectures for the paratope-epitope prediction: an encoder-decoder with attention ^87^ and the transformer architecture ^86^ (as described in Supplementary Figure 23). (B,C) Prediction accuracy (exact match of the predicted epitope to the test-set epitope) for both architectures as a function of a set of encodings, for degree-free (B) or degree-explicit (C) encodings, with (upper facet) or without (lower facet) residues binding with degree 1. Each point is an independent training instance and 10 different replicates per condition were performed. Learning on a paratope-epitope shuffled dataset was compared with learning on 1 600–40 000 (and 200 000 only for the Transformer architecture due to computational reasons) unique paratope-epitope pairs, while all conditions were tested on 100 000 unique pairs when available, or downscaled to reach 80% training and 20% testing (see Methods). The “motifs” encodings only allowed for 2 262 to 9 065 unique paratope-epitopes pairs (see Supplementary Figure 15A), which are referred to as One-Hot (Max) conditions).

SN and DN architectures achieved high median accuracy (0.982–0.99) for the binary classification use case on D1 (Figure 3C). >4 000 training sequences were sufficient to reach accuracy values >0.97 (Figure 3D). Prediction accuracy was slightly lower for SN and DN architectures on D2 (median 0.891–0.918) underlining the impact of the choice of the type of negative samples on the degree of difficulty of the ML task (Figure 3E). Interestingly, models trained only on D1 using the amino acid composition of the sequences, reached a substantially higher accuracy than random (Supplementary Figure 10C): shallow models reached 0.63 to 0.73 median accuracy trained on 40 000 sequences from D1 (Supplementary Figure 10, “AA-comp”) while all neural network architectures (SN5–20 and DN5–20) consistently reached 0.974 accuracy on D1 with 40 000 sequences on average across all antigens (Supplementary Figure 10B), which was very close to accuracies using the one-hot encoding (0.989, Figure 3). In comparison, models trained on the amino acid composition of 40 000 sequences from D2 reached 0.768 to 0.775 median accuracies (Supplementary Figure 10C), which is lower than accuracies using the one-hot encoding (0.823–0.918, Figure 3E). We further used an out-of distribution approach to evaluate models that were trained on D1 on their capacity to predict binding in the D2 dataset of the same antigen (Figure 3F) (i.e., assessing accuracy on the distribution-shifted dataset ^79^), and found lower median accuracy values (0.707–0.744), showing that training on D1 is not very informative to learn the delineation between binders and non-binders of D2.

As with sequence-based experimental datasets, D1 or D2 contain a mixture of sequences that bind to different epitopes on the target antigen, and it is not clear whether ML models trained on sequence datasets implicitly learn information on epitope specificity. One can naïvely assume that similar sequences bind the same epitope, and that the knowledge of the binding epitope of a few sequences could be sufficient to tell that other similar sequences would also bind with high affinity and with the same epitope. To explore this assumption, we clustered 25 000 binder sequences to 1ADQ_A (positive class in D1) according to their Levenshtein Distance (LD) and could observe 10 clusters (Supplementary Figure 14A), which did neither correlate with the epitope nor paratope encoding of these sequences, showing that similar sequences bound to different epitopes. We used an integrated gradients (IG) analysis ^80^ as an attribution method to identify which residues were important in the classification of binder sequences in Dataset 1 (Supplementary Figure 14B-C) ^39^. To this end, the SN10 architecture was trained on D1 of antigen 1ADQ_A and each sequence was annotated with its IG weights of the trained model; i.e., a measure on how each amino acid of this sequence quantitatively decided its output binding probability. The integrated gradient approach allowed to cluster sequences according to their paratope and epitope encoding (Supplementary Figure 14B) as opposed to clustering by sequence similarity, showing that the information learned by a neural network uncovers properties of the functional similarity between sequences and therefore may be used not only to find new sequences with high affinity ^32^ but also with desirable paratope or epitopes. The clustering was not due to a confirmation bias, as the clusters disappeared after shuffling the labels of each sequence (Supplementary Figure 14C). Therefore, Absolut! can be used to benchmark attribution methods for their usefulness in identifying the binding properties of antigen-specific antibodies.

We assessed whether ML architectures for binary classification followed the same performance ranking in Absolut! or in experimental datasets. We used an experimental dataset containing high-affinity (class 1) and low-affinity (class 2) antibody sequences to the tumor antigen HER2 ^39^ (Figure 3H). Using Absolut! sequences from D1, with the same amount of positives and negatives as in ^39^, the ranking of ML architecture performances were shared on both Absolut! and experimental datasets.

In the context of multiclass antigen binding prediction (on dataset D3), we used SN and DN architectures on multi-class classification varying ML model complexity (number of neurons 5–250 and number of layers). The classification performance (in terms of macro-averaged F1 score) decreased with increasing number of antigens (classes) included in the task (Figure 3G). Conversely, the accuracy values improved as a function of model complexity (number of neurons) with the largest models (n_neurons_=250) reaching between 0.67–0.932 median macro F1 depending on the number of antigens. In comparison, shallow learning models yielded markedly lower median macro F1 accuracies (LR: 0.36–0.66; NB: 0.01–0.34; RF: 0.–0.15; and SVM: 0.35–0.68 for SVM, Supplementary Figure 11A). The F1 scores for the largest model (n_neurons_=250) were higher than those of models trained on D3 with shuffled labels (median macro F1 < 0.02) or using only amino acid composition (median macro F1: 0.05–0.16, Supplementary Figure 11B).

To summarize, Absolut! is useful for comparing ML architectures and attribution strategies on different dataset specifications including the influence of different negative samples design

### Impact of the choice of negatives on binary classification of antibody-antigen binding poses

We investigated the relative performance of different ML strategies on a structural antibody-antigen prediction problem that classifies whether an antibody-antigen binding pose is of high affinity (the pose classification problem ^81,82^, Figure 4A) and assessed if the ML method rankings transfer to experimental datasets.

Using antibody-antigen pairs whose experimental antibody-antigen structures are known (“binding pairs”) as the largest currently available source of high-affinity binding poses, the DLAB-VS (Virtual Screening) pipeline by Schneider et al. ^81^ models the 3D structure of the antibody sequence ^83^, and generates 500 docking poses for a given antibody-antigen pair. The poses are compared to the experimental (target) structure using the “fnat” score (Figure 4A, top panel and Methods). Any pose that is similar to the target structure according to an fnat threshold was considered as positive (P). In contrast, remaining poses that are not “high affinity” can be separated into different types of negative examples, that can be included or not in the binary pose classification problem definition: Schneider et al. defined as negative examples poses of a binding pair that are highly dissimilar to the target structure (fnat < 0.1, incorrect poses (I)) as well as any pose generated from a an antibody-antigen pair that was formed by randomly matching the antibody sequence of a binding pair with another antigen (O), as commonly performed in PPI problems ^81,82^. Here, we reproduced the DLAB-VS pipeline on Absolut! datasets (Figure 4A, bottom panel) by selecting 10 000 antibody sequences of different affinity levels to 10 different antigens; extracting the energetically best 500 poses from the ∼6.8 million poses screened during exhaustive docking for each of the 100 000 antibody-antigen pairs. If the pair was of high affinity (top 1% affinity), we calculated the fnat between the poses and the optimal binding structure as target structure. Additionally to “P”, “I” (from the top 1% antibody binding sequences), and “O” poses, we defined new types of negative poses: any pose from a low affinity antibody sequence (“L”, top 1%-5%), or any pose from a non-binding affinity sequence (“N”, bottom 95%). The choice of the fnat threshold for positive poses, as well as the type of negative poses included in datasets, define a pose classification problem.

We reproduced the ML architecture of DLAB-VS ^81,82^, which takes as input the 3D encoding of an antibody-antigen pose as two cubic lattices, one for the paratope residues and one for epitope residues of the pose (Figure 4B). A 3-layered CNN architecture extracts structural information from the paratope and epitope lattices in parallel, while a dense layer returns the binary class of the input pose. Although the CNN architecture is translation invariant, it is not invariant to rotation, which opens up the possibility to optionally augment training datasets by duplicating a pose into its many possible rotations (n_Rotations_) to reduce overfitting.

We compared the ranking of different ML strategies for pose classification on Absolut! datasets to the rankings observed on experimental datasets. First, using a strict threshold for positive poses (fnat = 1), we reproduced a finding in Ragoza et al. ^81,82^ on experimental PPI datasets where data augmentation by rotation was necessary to avoid overfitting (Figure 4C, Supplementary Figure 13A). The AUC of the classification accuracy was 0.5 with n_Rotations_ = 1 (no data augmentation) and increased to 0.75 with as few as 5 rotations per pose. This represents an aspect in which ML behavior is observed to be similar in the Absolut! world and the real world. Interestingly, with a less strict definition of positive poses (fnat ≥ 0.5), including rotations did not have a beneficial effect anymore, which suggests calibrating the extent of data augmentation by rotation in future ML methods. Second, we explored the impact of the type of negative examples included in the training dataset (Figure 4D). The accuracy of trained models is shown either on the problem with only “O” negative poses (Figure 4D), (“I” and “O”), or all possible negative poses (Supplementary Figure 13B-D). We reproduced a finding from Schneider et al. ^84^ where including the “I” negatives slightly increased the classification precision for the “O” training composition. Depending on the tested dataset composition, the choice of negative poses had a non-negligible effect on the classification accuracy and can assist future pose classification strategies on experimental datasets.

### Impact of structural information on paratope-epitope prediction accuracy

Since the rules of antibody-antigen binding lie within the paratope-epitope interface, we formulate the prediction of the epitope for a given paratope as the “paratope-epitope” prediction problem (Figure 1A, Figure 5A). Paratope-epitope prediction may be formalized as a multi-class prediction problem, where a paratope is labeled by the ID of the epitope (in which case shallow learning methods can be used ^5^) or as a sequence prediction problem; e.g., Natural Language Processing (NLP) problem, where a paratope needs to be “translated” into an epitope by text generation. The NLP formulation allows predicting the binding epitope of new sequences (named as “neo-epitopes”) and towards new antigens, which makes the NLP formulation a more difficult task than classification, but more useful for purposes such as antibody, vaccine design, or neo-epitope discovery (Figure 5A).

We used our framework to propose mathematical expressions for antibody-antigen binding prediction tasks (Supplementary Figure 5). We considered a diverse set of possible paratope and epitope encodings suitable for NLP (Figure 5A), each differing in the amount of sequence and structural information contained. Degree-free encodings only contain paratope or epitope residues (involved in the binding), with or without indication of gaps (“-”) for a group of non-interacting residues between them, and have already been used for ML prediction on experimental antibody-antigen structures ^5^. Residues can be encoded as the amino acid name (without gap: “Sequence”, with gaps: “Aggregate”), their chemical property (“Chemical aggregate”), or just an “X” to denote that binding occurs at this position (“Motif”). Additionally, each paratope residue can be annotated with its “binding degree” (i.e., the number of bound epitope amino acid residues), termed “Degree-explicit” encoding. Simplified encodings were also considered after preserving only residues of degree 2 and more (D ≥ 2). Of note, each encoding defines a different biological paratope-epitope problem (Figure 1A, Supplementary Figure 1).

Depending on the encoding, an epitope can be represented by a different number of matching paratopes (Supplementary Figure 15A, blue histograms). In the sequence degree-free encoding, one single epitope was associated with a median of 24 841 paratopes. Therefore, the epitope→paratope prediction problem may contain multiple correct answers (as it does in this dataset). In comparison, a paratope encoding was only paired with a small number of different epitopes (Supplementary Figure 15A, gray histograms). Therefore, the paratope→epitope prediction problem generally has only one or very few answers, which is compatible with NLP architectures, or architectures trained to predict one possible answer (epitope). We, therefore, decided to focus on the paratope→epitope prediction problem.

We considered two commonly used NLP architectures using attention layers (Figure 5A and Supplementary Figure 23), due to the existence of long-range positional dependencies (Supplementary Figure 6): an encoder-decoder with a Bahdanau attention layer ^85^ and a transformer architecture ^86^ (Supplementary Figure 23A). Shallow architectures (SVM, LR, RF and NB; with one-hot encoding) provided baseline accuracies using the multiclass problem formulation on the same datasets (Supplementary Figure 19).

We assessed the performance on the paratope-epitope prediction problem of the two NLP architectures with all proposed encodings (Figure 5A) and using from 1 600 to 200 000 unique paratope-epitope pairs for training. Performance was measured by the fraction of either exact prediction match (accuracy) (Figure 5B,C), correct predicted epitope size (Supplementary Figure 16A), or the LD between predicted and target (test-set) epitope (Supplementary Figure 16B,C). For each condition, a shuffled dataset control was included (i.e., where the causal link between paratopes and epitopes was removed) to quantify prediction accuracy on randomized data.

The accuracies of both NLP architectures were low using degree-free encodings including all binding residues irrespective of their degree (Figure 5B, D≥1). The encoder-decoder failed to predict the cognate epitope. The transformer achieved 0.109 median accuracy with the aggregate encoding and 0.048 for the chemical encoding, with 200 000 training paratope-epitope pairs. Median accuracy on paratope-epitope shuffled data (negative control) remained under 0.003 for all encodings. Altogether, we concluded degree-free encodings not filtered for residues with high binding degree (D≥2) did not allow for the investigated architectures to learn the patterns of antibody-antigen binding.

Degree-free encodings with high binding degree residues (D≥2) yielded higher prediction accuracies using both architectures (Figure 5B, lower facet), suggesting that degree 1 residues are less relevant for binding prediction. The aggregate encoding was most predictive with 0.456 and 0.396 median accuracies for the encoder-decoder and the transformer, respectively, with 40 000 training paratope-epitope pairs (shuffled accuracies were lower: 0 and 0.006 for the two architectures respectively). The transformer reached a 0.517 accuracy when trained on 200 000 sequences. Interestingly, the sequence encoding and the motif encodings yielded markedly lower accuracies using the transformer, 0.173 and 0.072 respectively, still higher than random (0.009 and 0.066 from the shuffled control, respectively). Therefore, encodings that combine both sequence and structural information (the “Aggregate” and “Chemical” encodings), filtered on residues with degree two or more, are more predictive than sequence-only or structure-only (“Motif”) encodings. Of note, the increase in prediction accuracies using the “Aggregate” encoding (compared to the “Sequence” one) was not due to a simplification of the task by a decrease in the number of possible output epitopes (2 128 epitopes associated with 16 276 paratopes each, on average for the “Sequence” encoding; and 2 154 epitopes associated with 16 252 paratopes each, on average for the “Aggregate” encoding (Supplementary Figure 15A)).

We asked whether including the binding degree in the encoding (degree-explicit encodings) would further improve accuracy. By retaining residues of degree 1 in the encoding (Figure 5C, upper facet), the transformer reached high accuracies for all the encodings (between 0.634 and 0.731 for 200 000 training pairs) except for the motif encoding (0.102). Accuracies on shuffled controls ranged between 0 and 0.005 across encodings. The encoder-decoder, however, only performed well on the sequence encoding (0.689 accuracy on 40 000 training sequences) while it failed to predict the epitopes with the other encodings (accuracy= 0), which was associated with improper size of predicted epitopes (Supplementary Figure 16A, the encoder-decoder entered infinite repeat loops). This suggests the transformer is better suited for longer encodings. Finally, discarding degree 1 residues in explicit degree encodings further increased the prediction accuracy (Figure 5C, lower facet) for all encodings except the motif one, and the encoder-decoder reached between 0.903 to 0.944 median accuracy on 40 000 training pairs, slightly outperforming the transformer architecture that reached 0.867 to 0.909 on the same training dataset size. We also introduced “Gapped Motif” encodings where gaps tokens “-” are replaced by a their size (as number), and which led to poor accuracies (Supplementary Figure 17A-C, maximum accuracy= 0.012) with both NLP architectures, suggesting that the size of gaps between binding residues does not contain sufficient information to predict paratope-epitope matching.

Taken all together, we have shown that datasets of as low as 8 000 training paratope-epitope pairs can be leveraged to obtain high prediction accuracy (∼0.8) if structural information is included within the sequence-based encodings in the form of gaps and/or binding degrees. The relative performance of NLP architectures on the Absolut! dataset validated our previous results ^5^ on experimental antibody-antigen structures, where the “Aggregate” encoding enabled higher prediction accuracy than the “Sequence” encoding alone on antibody-antigen structures from the AbDb dataset ^16^. Interestingly, structural information has recently been shown to be predictive and more data-efficient not only for the prediction of paratope-epitope binding ^5^ but also TCR-antigen binding ^88^.

### Performance of paratope-epitope prediction models on unseen data

We investigated the extent to which the aforementioned high accuracies (Figure 5C) reveal generalizable trained ML models, and specifically, whether the ML models have been biased to recognize similar paratopes or epitopes in the training and test datasets (data leakage problem).

First, on the epitope side, we analyzed the accuracy and LD of the trained models separately on epitopes that were either present in both training and test datasets (shared) compared to epitopes unique to the test dataset (hereafter called “neo-epitopes”, Supplementary Figure 18A). The likelihood to observe an epitope in both training and test datasets is shown in Supplementary Figure 15B,C). The accuracy on neo-epitopes was always zero (Supplementary Figure 18B) and the LD between predicted and true epitope was always very similar to the shuffled control, showing that neither architecture was able to predict epitopes that were completely absent from the training dataset. We observed a decreasing LD on the test-unique epitopes using the degree-explicit encodings in an encoding-specific fashion (e.g., using the chemical, degree-filtered, degree-explicit encoding the median LD went from 0.521 in the shuffled group down to 0.482 using the encoder-decoder, Supplementary Figure 18A). Despite a very low prediction accuracy, the LD of predicted neo-epitopes was slightly improved if the epitope was similar to training epitopes, for the “Chemical” and “Aggregate” degree-filtered encodings (Supplementary Figure 20A,B), suggesting that some patterns shared between similar neo-epitopes and training epitopes were learnt.

Second, on the paratope side, we quantified the similarity between paratope encodings matching the same epitope, by clustering encoded paratopes using their pairwise LD (Supplementary Figure 21A). Encodings that enabled high prediction accuracy generated clusters of paratopes with the same color (epitope), and multiple distinct paratope clusters matched the same epitope, possibly revealing different structural binding modes. We designed a train-test separation strategy, where the paratope clusters found for each epitope were split between the training or the test dataset, such that elements of the same cluster and same label (epitope) do not appear in both datasets (called “no data leakage” condition). This strategy minimizes the chance of high accuracy in the test due to the presence of encoding-similar examples in the training dataset. We assessed the performance of NLP architectures trained on the “no data leakage” datasets (Supplementary Figure 21B,C), which defines a more complex task. Interestingly, trained models reached higher prediction accuracy in the “no data leakage” condition compared to random splitting of examples between train and test. Of note, we used the same hyperparameters in all conditions, and therefore excluded the risk that models would be biased to perform better on a certain data processing strategy via hyperparameter optimization. Altogether, by creating a new data preprocessing condition where similarity of encodings is controlled, we could exclude the possibility that the NLP architectures used were reaching high accuracy because of similarity, which suggests that the trained models were able to learn generalizable patterns.

## Discussion

### Absolut! allows unconstrained generation of *in silico* 3D-antibody-antigen datasets for ML benchmarking

To overcome the lack of datasets to benchmark ML prediction methods and compare their relative performance, we developed a deterministic antibody-antigen simulation tool (Absolut!) that allows unconstrained generation of *in silico* antibody-antigen binding structures (Figure 1). The formalization of immunological antibody-antigen binding prediction use cases as ML problems imparts the capacity to generate datasets parametrized to antibody-antigen prediction problems as they would occur in the real world. Along with Absolut!, we provide an extendable database of 1.03 billion antibody-antigen complexes (Figure 2, greifflab.org/Absolut/). This database was used to compare the relative performance of ML strategies (differing by dataset design, encoding, or ML architecture) to three types of antibody-antigen prediction problems: classification of high-affinity antibody sequences (Figure 3), structural classification of binding poses (Figure 4), and paratope-epitope prediction (Figure 5). For these three use-cases, we have shown that the rankings of ML strategies were transferable to the rankings observed on experimental datasets, and predicted ML strategies that maximize prediction accuracy. In particular, we propose strategies to include structural information that improve paratope-epitope prediction performance (Figure 5).

Synthetic datasets are valuable as long as they provide features that overlap with experimental data as well as operate, at least to a certain degree, within a physiological parameter space. We have shown that Absolut! datasets recapitulate eight levels of complexity present in experimental antibody-antigen datasets (Figure 2, Supplementary Figure 6, Supplementary Figure 7). In comparison to previous widely used more or less abstract coarse-grained representations of antibody-antigen binding (reviewed in ^89^), ranging from probabilistic models ^90,91^, the shape-space model ^92^, to cubic ligand-receptors ^92,93^ or structural coefficients for antibody clones ^94^, Absolut! allows a substantially higher level of intrinsic structural complexity of antibody-antigen binding. Indeed, the VAE-generated latent space of monospecific binding sequences to different antigens did not cluster antigen-specific antibodies but rather by sequence similarity (Supplementary Figure 14E-H) ^34^, highlighting the complexity of binding rules beyond similarity that are present in Absolut! datasets ^34^. Absolut! has also been used as an oracle to compare stochastic and Bayesian optimization strategies that aim to iteratively increase antibody affinity to an antigen ^95^. Therefore, by generating simulated data that reflect different forms of sequence signals of antigen specificity, Absolut! enables the *in silico* evaluation of computational and ML strategies for antibody-antigen binding prediction prior to experimental exploration.

### Rankings of ML strategies on Absolut! datasets are transferable to ML rankings on experimental datasets

We have shown that Absolut! synthetic datasets are useful to predict the relative performance of ML strategies, and importantly, that the rankings of ML strategies on Absolut! datasets transfer to experimental datasets (transferability). ML architectures that performed best in a binary sequence classification task on Absolut! datasets (Figure 3) also performed best on an experimental HER2-binder/non-binder dataset ^39^. Paratope-epitope encodings containing both sequence and structure information performed better than sequence-based encodings (Figure 5), as shown in ^5^. Finally, reducing overfitting by data augmentation (Figure 4C) and increased precision after including additional types of negative examples (Figure 4D) for binary pose classification, mirrored analogous findings on experimental datasets ^81,82^. Using Absolut! datasets, we have also shown that the required size of datasets for generative models transferred to a published generative model on experimental dataset ^96^. Therefore, despite the simplifications taken in the Absolut! framework as compared to biological data, the biological levels of complexity preserved in Absolut! datasets suffice to assist ML design useful to ML investigations on experimental datasets.

### Classification of antigen-binding can be improved by negative class design

We leveraged Absolut!-generated datasets to show that the performance of binary classification of antibody sequences (Figure 3) or structures (Figure 4) depends on the definition of non-binding antibody sequences (Figure 3E,F). This is a helpful result in the context of experimental enrichment-based antibody screening ^21,97^, where each enrichment step defines “improved binders”. Our results indicate that the non-binders in a late enrichment step may represent a good negative class as opposed to the non-binders in the early enrichment steps, since they represent the low affinity negative sequences in D2. In the context of antibody-antigen pose classification, in agreement with findings on experimental antibody structures ^81^, we showed that it is also possible to generate negative datasets with Absolut!, and that the inclusion of incorrect poses of a high-affinity antibody-antigen pair increases classification precision while defining a more complex task (Figure 4D). Therefore, Absolut! enables the generation of different types of samples in datasets with various controlled levels of difficulty, which is currently unfeasible experimentally. For instance, by generating antibody sequence datasets with different modes of binding to an antigen, we have shown that Absolut! allows the prospective performance evaluation of predicted outputs from generative models across those different classes of sequences ^98^.

### Structural information improves sequence-based conformational paratope-epitope prediction

A major open question in the immune receptor ML field is to understand how beneficial it is to include structural information in binding prediction tasks ^33^. Several recent studies have reported improved paratope or epitope prediction ^49,50^ when including an attention mechanism to the antibody or antigen structure. Under the paratope→epitope prediction problem, we compared encodings with different levels of coarse-grained sequence and structural information, allowing us to regroup multiple paratope-epitope sequences under a shared pattern and embed binding rules within the encoding. We were able to delineate a hierarchy of predictive structural and sequence features within the paratope and epitope encodings (Figure 5B,C). The presence of the gaps in the encoding improved accuracy (compare “Sequence” and “Aggregate” encodings), which mirrors previous findings of us on 3D-structural experimental data ^5^. Solving the problem of pose classification would be a practical breakthrough for antibody discovery because it predicts both if an antibody binds to an antigen (as sequences or modeled structures), and where and how it binds (through classification of possible poses), and has the potential to combine information from both sequence-based and structural-based datasets as recently suggested by Schneider et al. in the DLAB-VS pipeline ^81^. Here, using Absolut!, we were able to reproduce and benchmark the behavior of DLAB-VS to poses generated from different types of antibody-antigen pairs. Therefore, Absolut! fosters the development of new methods before the data are available and enables the comparison of the predictive power of datasets with a varying richness of embedded structural information.

### The challenge of generalization and capacity to predict on unseen data

Another major question in antibody discovery is to what extent rules learned from a set of antibodies binding to a set of antigens are applicable to other antibodies or antigens (generalization). For instance, antibody structure or paratope prediction studies typically allow similar paratope or epitope sequences to be in both the training and testing dataset, and therefore it is challenging to disentangle whether their high prediction accuracy is due to the presence of a homologous antibody/antigen sequence or structure in the training dataset (“data leakage” problem). In the context of paratope-epitope prediction, we have shown that several encodings lead to high accuracy when predicting the target epitope of new paratopes (Figure 5B,C). However, the high accuracy was due to the successful prediction of epitope targets that were already described in the training dataset (Supplementary Figure 18, Supplementary Figure 15B,C) while the accuracy of unknown epitopes during training (here termed neo-epitopes) was very poor (accuracy close to 0 and LD similar to shuffled controls). Certain encodings provided a slight decrease in LD between predicted and target neoepitopes suggesting that a substring or subpattern could be predicted for some neo-epitopes (Supplementary Figure 18). Of note, the LD of predicted epitopes to their target was slightly better for neoepitopes sharing subpatterns with training epitopes (close neoepitopes) than for those that are more dissimilar to the training dataset (Supplementary Figure 20). The prediction of neoepitopes has also been proven to be difficult on the TCR binding prediction problem, for TCR sequences distant from the training dataset ^99,100^. Furthermore, the complexity of paratope→epitope prediction also depends on the number of possible answers for a paratope (Supplementary Figure 15A, gray histograms). Due to the dataset design of millions of CDRH3 to few antigens (as in the real-world), the possible answers were limited and the tested NLP architectures could reach high accuracy. In contrast, when studying either epitope→paratope prediction or datasets with mutant antigens, a many-to-many approach will need to be developed, a Multiple Instance Learning formalism ^101^ could help learn from datasets with higher levels of cross-reactivity ^101,102^. Absolut! datasets will be critical for optimal design of such models.

We also developed a strategy to assess the extent of data leakage on trained model performance, due to similarity of encoded paratopes in training and test datasets, by clustering similar encoded paratopes, and assigning the clusters to either the training or test dataset (Supplementary Figure 21, Methods). Interestingly, the prediction accuracy was slightly higher using the “no data leakage strategy”, suggesting that binding rules could be learnt beyond merely similarity. However, these findings do not exclude a potential influence of alternative types of similarity, e.g., conserved features reminiscent of structural similarity ^103^. Altogether, our results suggest that the analysis of similarity in antibody-antigen datasets as a confounder (instead of assuming that similar antigens bind the same target), and studying the capacity of different encodings to provide a functional similarity landscape in the encoded space, represent promising avenues to increase generalization.

### Limitations of the Absolut! framework in terms of reflecting the complexity of antibody-antigen binding

Although the Absolut! framework includes the above-mentioned levels of complexity, the main limitations of our model with respect to the current understanding of antibody-antigen binding are: (i) proteins do not conform to lattice representations based on fixed inter-amino acid distances and rigid 90-degree angles. (ii) Protein binding depends on additional factors not included in this model such as solvatation and 3D distribution of charges, and the possible flexibility of the antigen structure depending on the binding antibody. (iii) Only the CDRH3 loop (as opposed to VH-VL) was simulated, which is the most variable chain for antibody specificity ^5,6^, thereby neglecting the topological constraints with other CDR loops and their contribution to binding as well as the influence of the constant region on antibody binding ^104^.

### Potential improvements for the Absolut! framework to include antibody full chains

Possible improvements of the Absolut! framework are: (i) within the lattice framework, it may be possible to add local geometric ^105^ or angle constraints ^106^ within consecutive groups of amino acids, and thus, add an additional level of structural complexity on CDRH3 sequences binding an epitope. (ii) The 90-degree angles can be refined to a larger set of angles. This will impact the lattice resolution and the antigen discretization might be redefined for higher lattice resolution. (iii) Since the CDRH3 peptide is anchored on a conserved region between antibodies, geometric constraints could be included between the endings of a CDRH3, but this would require care since Absolut! only considered 11 amino acids that do not necessarily include both ends of the CDRH3. (iv) Including structural constraints between more than one loop could enable simulating the CDRH3 and CDRL3 chains together, or even all CDR antibody chains although the computational cost of evaluating each pair of compatible CDRH3 and CDRL3 structures without further optimizations seems unrealistic to achieve at the moment. (v) The top 1% binding thresholds used across the manuscript are an arbitrary choice of the user and can be adapted to any other simulation or experiment data-informed settings. Relative thresholds per antigen mimic the fact that experimental enrichment of high-affinity sequences is usually affinity threshold agnostic and therefore should not be compared across antigens. (vi) It will be possible to compare different energy potentials and to include non-linearities between the energetic contributions of neighboring amino acids once more experimental affinity distributions have become available. Docking-based affinities are not directly suitable to this end ^107^.

There is the risk that trained models on synthetic datasets would increasingly learn synthetic design biases with increasing size of these datasets, which we cannot exclude. However, learning the biases is only a problem if it decreases transferability. We believe the fact that transferability of ML method rankings was observed for a range of specific use cases as shown in this study (Figure 3C-D, Figure 4, Figure 5), supports the notion that biases are to a certain extent negligible for the use cases tested. Furthermore, the fact that sequence-encoded paratope-epitope prediction accuracy was low for the investigated dataset size and model complexity (Figure 5) suggests that the ML models were not able to learn lattice biases. Since experimental datasets also contain their own biases, these biases may be integrated in the future.

Finally, ML investigations shown will require follow-up studies. Beyond detailed benchmarking of already implemented encodings, investigations of graph-based encoding, physicochemical properties, full torsion angle, or 3D voxels, the addition of physics-based priors may be needed ^108^. Furthermore, comparative insights into different ML architectures such as Transformers, graph-based ML ^24,44,109^ as well as attribution strategies such as integrated gradients to map the ML models’ predictions to the underlying biology are warranted ^108^.

## Materials and Methods

### Representation of antibody-antigen binding

We used the Ymir framework that can represent binding between ligands and receptors in a 3D-lattice and has been used to model affinity maturation in germinal centers ^67^. Briefly, a protein is a set of consecutive (neighbor) amino acids on the 3D lattice (Supplementary Figure 2), meaning only 90 degrees angles are considered, and only one amino acid is allowed per lattice position. A protein chain is represented by its amino acid sequence, a starting point in space, and a list of relative moves in space to determine the next amino acid position (Straight (S), Up (U), Down (D), Left (L), Right (R)) according to a moving observer’s coordinates (similar to a plane pilot) ^67^. From the starting point, the first move is defined from the coordinate system (Ox for straight, Oz for up, etc.), and the “observer” moves or turns. The next move is relative to the observer. “Backwards” (B) is only allowed as the first move (otherwise, it would collide with the previous position). For instance, “(25, 30, 28), BSLDUSUSRS, RGAYYGSSYGA” or “(26, 30, 27), DLURSLDDUL, YYGSSYGAMDY” are two protein structures of the size “10 moves” or 11 amino acids. In particular, an antigen is a multi-chain Ymir protein (structure and amino acid sequence), and the antibody is represented either as an 11 amino acids sub-sequence of its CDRH3 (sequence only) or as with a 3D conformation as an Ymir protein (structure and amino acid sequence).

### Automatic discretization of antigens

The position of each amino acid within a protein’s PDB structure of a protein (the antigen) requires transformation into 3D integer lattice positions in order to use the Ymir framework (Figure 1B). Intuitively, this process is similar to rescaling the PDB structure, finding the best rotation and translation, and converting each amino acid position into a possible integer position around it with the best fidelity to the original structure. We used the software LatFit ^64^ to perform this task, using its dRMSD (Root Mean Square Deviation, providing a measure of the distance between experimental and lattice positions) minimization algorithm. LatFit iterates through multiple possible reconstructions of the protein structure on the lattice. It starts from the first amino acid of the protein and assigns it to (0,0,0) in the lattice. It iterates through all possible neighboring positions for the next amino acid, thereby generating multiple starting lattice structures of size two amino acids. Each structure is ranked according to its best fit to the underlying PDB positions, determined by the dRMSD, calculated on the best rotation/translation between this structure and the original PDB structure. LatFit iteratively stores a list of N best-ranked structures of lattice positions for the first K amino acids, and extends each structure with every possible position for the next amino acid, before ranking all the newly extended structures by their dRMSD and keeping only the N best ones. When residues are missing in the PDB file, or for instance when a protein loop was not structured enough to be reconstructed from a crystal or cryoEM, LatFit automatically starts a new chain starting with the next amino acid. Finally, since LatFit cannot parse residue insertions and can only process residues from a single chain in the PDB, we used pdb-tools ^110^ to rename amino acid insertions by shifting residue ID and merge all chains of interest into a single chain and adapted the LatFit C++ code to create new chains when necessary.

Altogether, the following LatFit parameters are considered during the discretization of the antigen:

i. *Type of discretization:* Which position do we consider for an amino acid (the position of the C Alpha “CA”, the centroid center “CoM”, or the fused center of positions of all the amino acid atoms “FuC” that we implemented);
ii. *Lattice resolution:* The distance between two lattice points in Å;
iii. *N:* The number of possible lattice chains that LatFit keeps at each iteration.

We compared the discretization types and lattice resolutions (Supplementary Figure 3) and reached the conclusion that a lattice size of 5.25 Å using the “FuC” discretization is optimal to minimize the dRMSD quality of discretization while avoiding an excessive number of empty cavities inside the protein when the lattice resolution is larger. As a tradeoff, values for N (retained lattice chain solutions) larger than 25 did not significantly improve the quality of discretization (dRMSD), although special cases of large proteins needed N=50 for LatFit to find a solution (PDB 3VI3 chains AB and 5JW4 chain A). Therefore, the discretization step preserves both the native antigen topology in 3D-space (typically <3.5Å, Supplementary Figure 3) as well as the native surface amino acid distribution (Supplementary Figure 7A).

### Removal of aberrant geometries

Discretization artifacts might leave empty positions (cavities) inside the antigen, or generate topologies that would not be physically accessible by antibodies in vitro/vivo (Supplementary Figure 3B). For instance, a free position surrounded by five amino acids would be a dead-end if an antibody bound there. Furthermore, an antibody should not occupy a free position surrounded by four antigen amino acids in the same plane (a “donut hole”) because the antibody’s simulated peptide end would “pass through” the antigen. We used a recursive algorithm that identifies “aberrant geometries” and blocks access to all positions surrounded by four planar (donut) or five or six (dead-end) inaccessible positions until all such positions had been blocked. We added a visual inspection step for complicated topologies where the antibody may pass through the antigen (manual curation by adding blocking positions).

### Assignment of glycans in the discretized antigens

Glycans show a high diversity of tree structures anchored on proteins as post-translational modifications and are of critical interest for antigen recognition because glycans on viral glycoproteins shield against antibody binding ^111^. However, the proteins used in PDB structures are often synthesized on expression systems that do not produce associated glycans, and crystal structures are more difficult to obtain in the presence of glycans ^112^. A glycan is typically a tree of sugar molecules anchored on a protein via a β-(1,4) linked N-acetylglucosamine (*NAG*) or N-acetylmuramic acid (*NAM*) stem. Here, we modeled the glycan by adding a “forbidden” position on the closest available lattice position to the stem position, around the discretized antigen (independently/after the LatFit discretization step was performed). The flexibility and weak binding of glycans ^113^ are therefore ignored. Starting from the PDB file, the following steps were carried out to assign glycan positions in the 3D lattice:

i. *identification of glycan stems:* Glycans are parsed from the PDB file by identifying NAM or NAG, and by finding which amino acids are bound to them on the chains of interest. Since the NAM and NAG are relatively big (similar size to an amino acid), we decided to consider the barycenter position of all atoms of the NAM or NAG as one amino acid-like element and to discard the remaining sugar ramified structure that is normally flexible and do not necessarily preclude antibodies to be at the original positions of the ramifications.
ii. *identification of root* amino acid*s for the glycans in the discretized antigen:* Since the preprocessing of insertions by the tool pdb-tools shifts the amino acid IDs in the initial PDB, we keep track of the amino acid bound by each glycan stem by noting their position in space instead of their ID. After discretization, we retrieve where these amino acids are (independently of their ID) positioned in space before and after discretization. The glycan position in the lattice should be attributed in the vicinity of this amino acid, and as close as possible to the real space position of the NAG or NAN barycenter.
iii. *definition of suitable positions to place glycans (“Atmosphere”):* In some cases, the discretization leaves empty space inside the antigen, which is not a suitable position for glycans that are typically located at the surface of proteins. We designed a recursive algorithm to identify “atmospheric positions” (Algorithm 1) in space that are defined as all positions that (i) may be accessed from the exterior by 90-degree moves, and (ii) that are either of Hamming distance one (nearest neighbor) or distance two (non-nearest neighbor) from an antigen residue. This excludes buried empty lattice points. In the algorithm, [global] means the variable should be a global variable shared between all calls of the recursive search, to label previously visited positions.

#### Algorithm 1

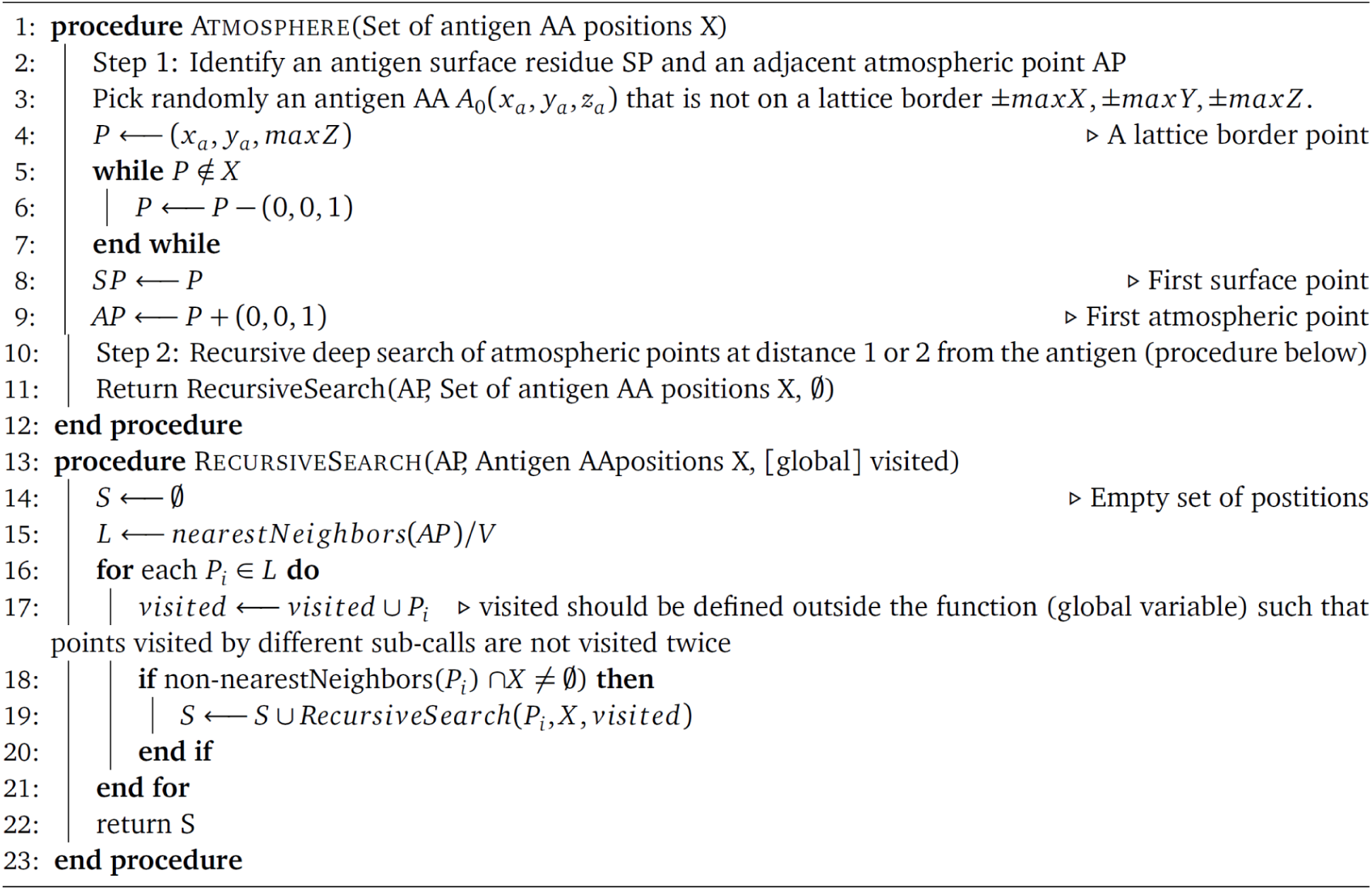
Atmosphere: returns empty lattice points around (but not buried inside) an antigen.
iv. *assignment of possible suitable free positions to each glycan:* to position a given glycan in the lattice, each glycan will be assigned a set of possible positions. Let *A*_i_ be the rooting antigen amino acids for a glycan *i*, and its position *(A_ix_, A_iy_, A_iz_)* in the lattice, from the discretization step. Let *G_i_* be the original glycan floating position (NAG or NAN barycenter) and *D_i_*the closest lattice integer position (rounding up all positions to integers). Among identified suitable positions (the “Atmosphere”, the neighbors of *A_i_* in the lattice are considered as potential positions for the glycan. If not, *D_i_*and its neighbor lattice positions are considered. If there are still no such positions, its non-nearest neighbors are also considered (Algorithm 2, possible positions).
v. *greedy assignment of positions to glycans:* The possible positions for each glycan are sorted by their distance to the original position *D_i_* and the glycans are iteratively assigned to a position in a greedy manner (Algorithm 2). For each glycan, its closest available position is assigned to it except if another glycan has only this choice (in which case the latter is assigned the respective position) or if another glycan has the same position with a smaller distance. The procedure is repeated until all glycans are assigned a lattice position. The greedy approach allows us to solve conflicts between spatially close glycans. Although it may be possible that there are not enough free positions for all glycans, in which case one position represents two or more glycans, we did not observe such a case for the antigens investigated.

#### Algorithm 2

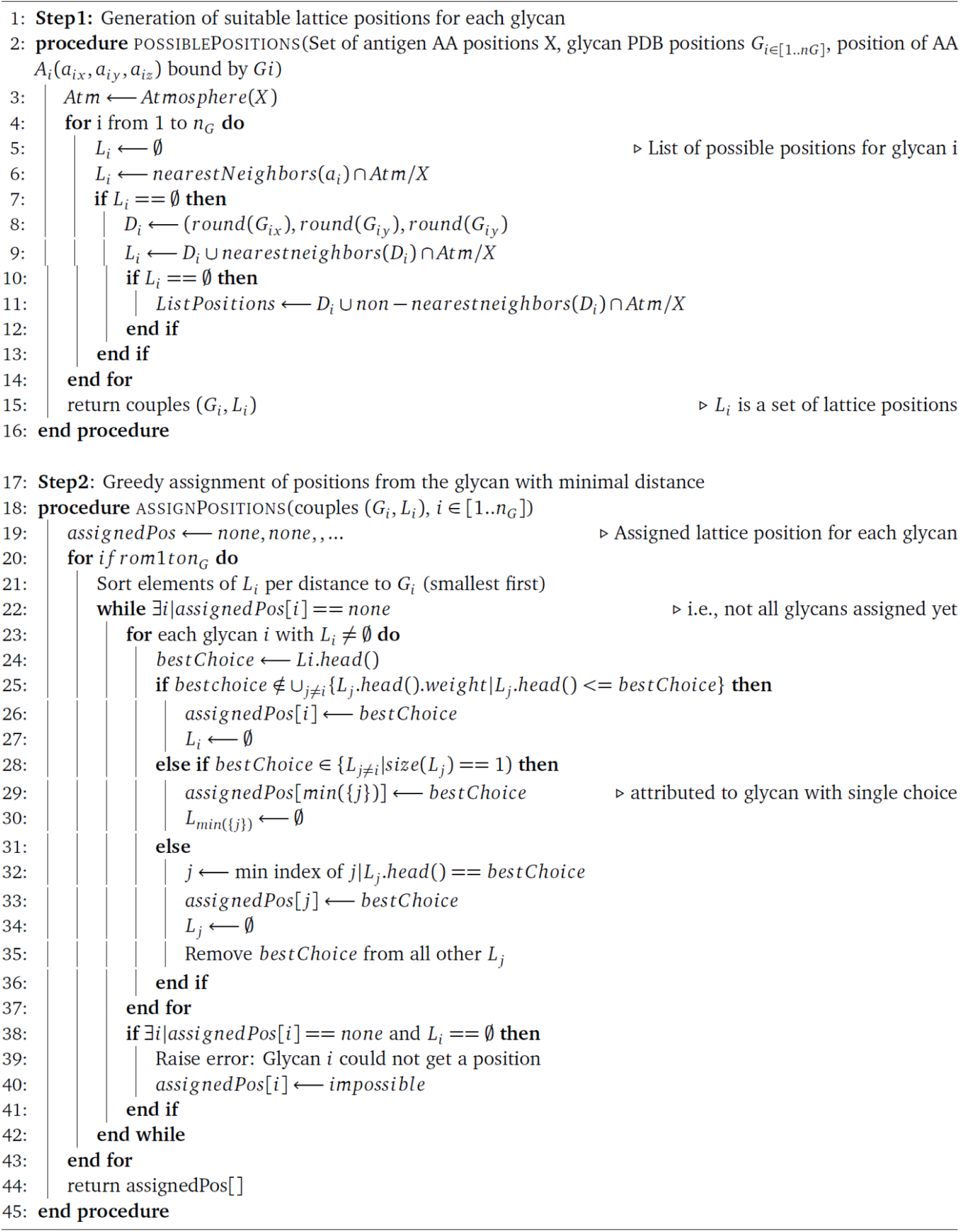
Glycan assignment: for each glycan, the closest atmospheric point in the lattice is chosen, starting from glycans with smaller distances to atmospheric points.

### Calculation of CDRH3 binding energy for a binding structure

The affinity of an antibody (CDRH3) binding to an antigen depends on its binding confirmation ^67^. The antigen structure is pre-defined by the user (see Section above on discretization), and the structure of the antibody is not known a priori (the considered 11-mer amino acid subsequence). In the Ymir lattice framework ^67^, only neighboring but non-covalently bound amino acids are considered interacting, and the strength of the interaction is defined by the Miyazawa-Jernigan ^114^ potential, an empirical, experimentally derived potential that gives a binding energy for each pair of interacting amino acids (Supplementary Figure 2A). For a particular conformation of the antibody on the antigen, by summing the binding energies of interacting amino acids, three energies can be derived: i) the interactions between residues of the antigen (folding energy of the antigen), ii) the interaction between antibody residues (folding energy of the antibody) and iii) the interaction between residues of the antibody and the antigen (the binding energy). The total energy of the binding is the sum of the three interaction types. Since the antigen structure is assumed constant, its folding energy is constant and ignored. In order to find how an antibody CDRH3 sequence binds to the antigen, the antibody-antigen binding structure with minimum total energy (the most stable) will be defined as the “energetically optimal binding conformation of the CDRH3 to the antigen”, and the binding energy will be recorded as “binding energy of this CDRH3 sequence to the antigen”. It is this binding energy that is used for all downstream analyses. Of note, the most stable binding is not necessarily the one with the lowest binding energy, due to trade-offs between folding and binding energies.

### Exhaustive search of the optimal binding (exhaustive docking)

The energetically optimal binding conformation is calculated using a previously developed algorithm ^67^ that performs an exhaustive search (Supplementary Figure 2B). By enumerating all possible binding structures (which is feasible in the lattice), we guarantee to find the optimal binding, while usual docking approaches rely on optimization procedures which can always be stuck in local optima, and therefore called our approach “exhaustive docking”. We restricted the calculations to peptides of 11 amino acids only as we could show that the lattice binding energy cannot directly be compared between binding sequences of different sizes ^67^ and the calculation for longer peptides was not computationally feasible. In brief, all possible 3D structures of 11-mers in the lattice were enumerated (independently of their amino acid content), provided they had a minimum predefined number of 11 contacts. Among the used antigen structures, an average of 6.8 million 11-mer structures was found per antigen (min 27 500, max 44.3 million) filtered according to a minimum of 11 contacts to the antigen (i.e., the sum of degrees of amino acids on the antibody 11-mer). Allowing a lower number of contacts led to an increasingly higher number of structures, and we noted that found optimal structures always had many more contacts than 11 (22 on average), suggesting that this contact threshold could further be increased). Finally, when treating a CDRH3 sequence, all its consecutive 11-mers amino acid sequences (called “antibody slices”, Supplementary Figure 9) are derived and assessed for binding and folding energies according to all possible binding structures on the antigens. The optimal binding conformation of the 11-mer with the lowest binding energy is kept as the structural binding of the CDRH3. CDRH3s of less than 11 amino acids were discarded (6 919 242 remaining out of 7 317 314, see next paragraph).

### Antigen and CDRH3 data source for the generation of high-throughput structural antibody-antigen binding structures

#### Antigen dataset

In order to build the Absolut! database, we discretized 159 antigens sourced from the AbDb database, which collects all non-redundant PDB structures of antibody-antigens complexes, as a selection of therapeutically relevant antigens, to allow for the comparison of our epitopes against experimentally known binding regions (Supplementary Figure 7E). The list of antigens and their species is shown in (Supplementary Table 1). *CDRH3 sequence dataset:* We used a total of 7 317 314 unique CDRH3 sequences of murine naive B cells ^63^. Only the 6 919 242 CDRH3 of size 11 or more were used for binding calculations. For comparison, we used 1 million healthy human sequences of size 11 amino acids or more of naive B cell BCRs from a set of 22.6 million CDRH3 amino acid sequences ^70^. Although our pipeline is valid for any CDR/FR sequence, antibody binding was computed for a single binding site, the CDRH3 of the antibody heavy chain, because it is the most important antibody site for antigen specificity ^5,6^.

### Dataset of protein-protein interaction (PPI) and Ab-Ag complexes (AbDb)

The dataset of protein-protein interaction complexes used to compare encoded interaction network topologies (Supplementary Figure 8) was obtained from 3did ^115^ (preprocessed data from Akbar et al. ^5^ was used) thus we followed the preparation, preprocessing, and filtering steps for the PPI data as described previously ^5^. Briefly, we annotated interactions between domains of different chain origin as interdomain and identical chain origin as intradomain, removed structures with domain description immunoglobulin and ig-like. We excluded the structures with non-interacting residues (gap) larger than 7 residues and finally we excluded protein-protein complexes larger than 300 residues long (to allow for a fair comparison against antibody-antigen complexes). The PPI dataset comprises 9 621 interdomain and 1 043 intradomain complexes. The dataset of Ab-Ag interaction comprises 1 200 of Ab-Ag complexes obtained from AbDb^16^.

### Absolut! computational consumption

The calculation of one binding energy/conformation of one 11-mer to one antigen takes on average one to two seconds on a CPU (depending on the number of structures in the exhaustive search), and 8 seconds per CDRH3 (Supplementary Figure 4F). In total, 16 000 CPU hours were needed per antigen on average, and approximately 3 million CPU hours were required to generate the Absolut! database.

### Clustering of antibody binding hotspots

From a structurally annotated list of CDRH3 sequences; i.e., the lattice binding structure and their associated epitope (set of bound antigen amino acids in the lattice). We defined a binding hotspot as a set (cluster) of epitope residues shared by multiple binding structures on the antigen, in the following way (see Supplementary Figure 5 for visualization of definitions and Supplementary Figure 22):

i. *Assign a list of binding epitope residues to each CDRH3.* Each CDRH3 binding structure was translated into a list of positions bound on the antigen (the epitope). Additionally, to differentiate only “important” positions on the antigen, we first selected residues on the CDRH3 that were a neighbor to at least *D* residues on the antigen, and then created the list of epitope residues A_i_ bound by these CDRH3 residues.
ii. *Separate sets of K antigen residues that explain the epitopes of all CDRH3s.* We aimed to identify binding hotspots as antigen residues bound by many CDRH3s. For each antigen residue, we created the list of all CDRH3 sequences that contain A_i_ in their fingerprint. A set of K residues {A_i_} “covers” the CDRH3 sequences whose epitope contains all the A_i_. The identification of binding hotspots translates into the following question: *“Can we find a minimal list of K-sets of antigen positions that cover all the CDRH3 sequences?”* This is a “set cover” problem, which is NP-complete ^116^. We decided to implement a greedy approximation solution by first taking the set of K positions that cover the most CDRH3s, then removing these CDRH3s and iteratively determining the set of K positions that cover the most remaining CDRH3s (Algorithm 3). Therefore, each set of K positions mirrors the binding pattern of a group of CDRH3s.
iii. *Translate each K-set into a binding hotspot:* For each K-set, we analyzed which residues are bound by this group of CDRH3s. The fraction of CDRH3 structures binding to each residue of the antigen was computed. The residues in the K-set were all bound by 100% of the CDRH3s. However, other positions might also be bound by 100% or a high enough percent and would be grouped together as an epitope if this was the case.

Therefore, in our framework, “a binding hotspot” is a cluster of antigen residues bound by a group of CDRH3s with K common positions in their binding epitope. Furthermore, a CDRH3 was named hotspot-specific if it contained the K common binding positions of this epitope. Note that a position can belong to two hotspots. It is not impossible that CDRH3s would have structures overlapping two hotspots (although very unlikely), in which case they will be classified as binding to both hotspots.

#### Algorithm 3

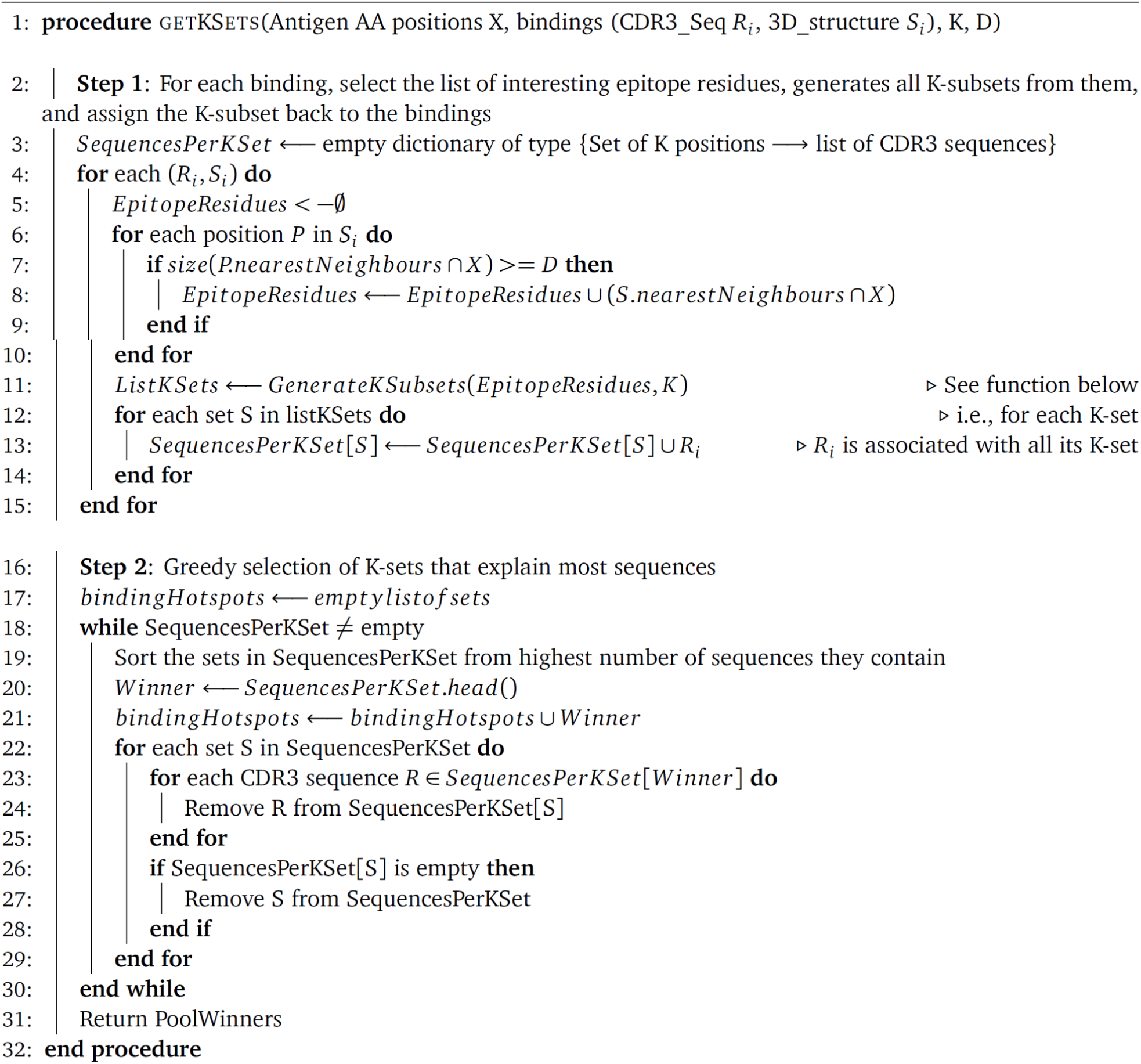
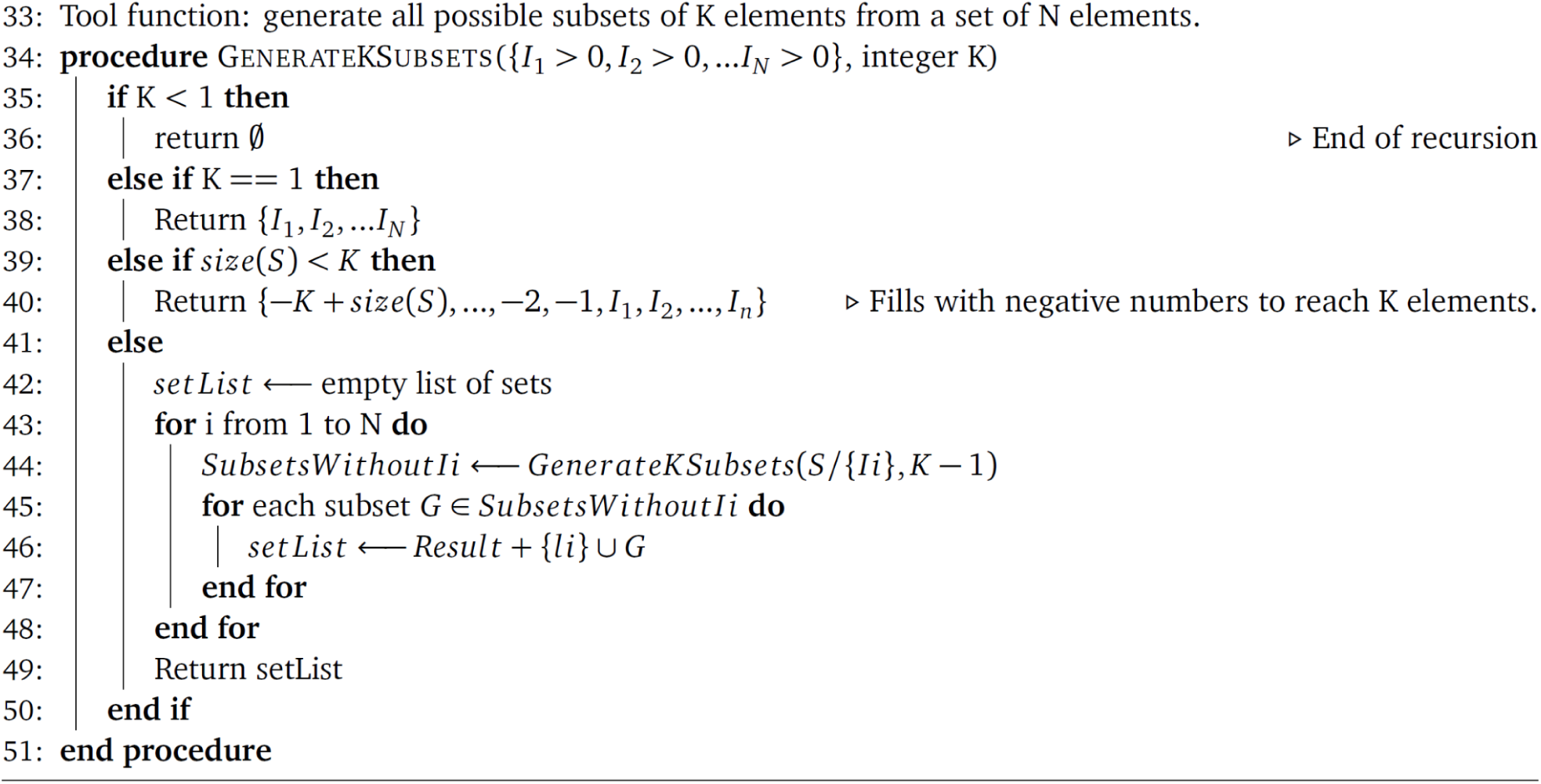
Clustering of binding hotspots as sets of K positions on the antigen, The epitope of each binding *i* is taken as a set of *N_i_* antigen positions binding antibody residues with minimal interaction degree *D*. All subsets of *K* positions of the *N_i_* are generated. The K-sets that represent the most binding epitopes become binding hotspots.

### Feature extraction from annotated antibody-antigen binding structures

Definition of affinity classes (Figure 2B): Starting from 7 317 314 unique murine CDRH3 sequences ^63^, 6 919 242 CDRH3s of 11 or more amino acids were kept and a total of 34 395 055 11-mers were generated (i.e., on average, 4.97 11-mers are generated per CDRH3). The same 11-mer can occur in different CDRH3 sequences, leading to 22 684 073 unique 11-mers. As explained above, the binding energy and structure of a CDRH3 to the considered antigen, are defined by its 11-mer with the lowest binding energy.

The Absolut! framework and database allows generating datasets with antibody-antigen binding sequences represented as a CDRH3 or a 11-mer sequence. Binding thresholds and affinity classes were defined based on the CDRH3 sequences, for each antigen separately. Binder CDRH3 sequences were defined as top 1% binders and ranged between 67 761–81 255 unique CDRH3 sequences (median 70 024). This number varied between antigens, due to multiple CDRH3 sharing the same binding energy as the threshold energy. Other CDRH3 binding classes such as the top 1%–5% binders, or the bottom 95% non-binders were defined analogously (Figure 2B).

However, at the 11-mer level, the top 1% binders were *not* defined as the top 1% of the 22 million unique 11-mers, because the binding energy of each CDRH3 was calculated among 4.97 different 11-mers on average, i.e., only keeping the energetically lowest 11-mer and discarding the other 3.97 11-mers. Therefore, an equivalent threshold would be around the top 0.2% best 11-mers. Instead of using a relative threshold proper to the 11-mers, we decided to define the “binder” or “top 1%” 11-mers based on the same absolute binding threshold as the top 1% CDRH3 sequences to the considered antigen, i.e, the list of 11-mers that would allow a CDRH3 to be a top 1% (CDRH3) binder to this antigen. We obtained between 28 486 and 97 618 unique 11-mers as “top 1%” binders per antigen (median 53 753), as shown in Supplementary Figure 9A. If the CDRH3 sequences did not share any 11-mer, we would have expected to obtain around 69 000 different binders for each antigen (exactly 1% of the number of CDRH3s). Lower numbers of top 1% 11-mers meant that the same 11-mer was isolated from multiple different CDRH3s, while higher numbers of top 1% 11-mers reveal a higher diversity of high-affinity 11-mers within the top 1% CDRH3s, i.e., that one CDRH3 contained more than one 11-mer over the binding threshold.

The strategy of defining affinity classes CDRH3 or 11-mers according to the same energy threshold allows interchangeably using 11-mer or CDRH3 binders for an antigen to build ML datasets. Throughout the manuscript, we used the 11-mers derived from CDRH3 sequences, to work on sequences of equal size.

#### Binary classification of binders (Figure 3A–F)

The “top 1%” 11-mers were isolated for each antigen (median 53 753 unique sequences, as explained above), and 25 000 of them were taken and labeled as “binders”. A random selection of 25 000 non-binding 11-mers (dataset 1, within the bottom 99% CDRH3 energies) or suboptimal binders (dataset 2, top 1%–5% CDRH3 binding energies) was taken. For both datasets, the non-binder 11-mers were sampled from CDRH3 of the same size distribution as the CDRH3 of origin of the binders, in order to avoid obvious amino acid composition bias, since CDRH3 sequences tend to start by “CAR” and shorter CDRH3s have, for instance, higher usage of C, A, and R amino acids.

#### Multi-class classification of antigen target (Figure 3G)

Among the 159 antigens, several antigen structures originated from the same protein (but different PDBs, see antigens with “*” in Table S1). The binding structures of the CDRH3s to the remaining 142 non-redundant antigens were kept. When K antigens were randomly picked to perform classification, only the top 1% binders to at least one of these K antigens were kept (we discard 11-mers that bind none of them, and perform the classification among binders only). Further, since we opted for multi-class techniques, we discarded any 11-mer that bound more than one antigen among the K-selected antigens, explaining the saturation in the number of available sequences as a function of the number of antigens in Supplementary Figure 9. In the future, multi-label techniques may also be tested without discarding these cross-reactive sequences.

#### Paratope-epitope prediction (Figure 5)

The structural annotation of all the binding 11-mers to the 159 antigens was transformed into epitope and paratope encoding, as described in Figure 5. For each paratope and epitope encoding, each encoded pair was kept only once.

### Assessment of physiological relevance of binding structures

Surface residues in Supplementary Figure 7A were identified using the module `findSurfaceResidues` in `Pymol ver. 2.1.0` ^117^. For Supplementary Figure 7B, for each of the 159 antigen-antibody complexes that were used to generate the library of *in silico* antigens, the position of the antibody was kept unchanged during the discretization of the antigen using Latfit, therefore the lattice antigen is shown with the experimental structure of the antigen.

### Machine learning

#### Data processing of sequence-based datasets

For the binary classification tasks (Figure 3A-F), 11-mer amino acid sequences were encoded as one-hot encoding (binary vector, dimension 220), as input features for the architectures shown in Figure 3B. Dataset 1 (binary classification) was generated with 40 000 unique binder 11-mers (Supplementary Figure 9A), and the same amount of unique non-binders (defined differently in D1 and D2). The number of used training sequences is indicated in the figure legends, while testing was performed on 40 000 sequences. For 9 antigens out of 159 antigens, only 56 900 to 78 000 sequences (binder + non-binders) were available in total, and the training ratio was downscaled to 80% training and the remaining sequences for testing (see Supplementary Figure 9A for the distribution of available binding 11-mers per antigen). For multilabel classification (Figure 3), the amount of available sequences for Dataset 3 is shown in Supplementary Figure 9B. When at least 300 000 sequences were available, 100 000 sequences were used for testing and 200 000 for training. For the ML use case of 10 antigens or less, less than 300 000 sequences were available and 80% of them were used for training and 20% for testing.

#### Data processing of encoded paratope-epitope pairs

For the paratope-epitope prediction problem, a list of unique paratope-epitope encoding pairs was generated by taking the structural annotation of all 11-mers from the top 1% affinity class. Therefore, the same encoded paratope-epitope pair can represent the binding of multiple 11-mer sequences to different antigens and is only repeated once in the paratope-epitope dataset. The number of available unique paratope-epitope pairs is shown in Supplementary Figure 15A. The number of used training pairs is indicated in the figure legends (between 1 600 to 200 000 pairs) while 100 000 pairs not present in the training dataset were used for testing. For the “motif” encodings, only 2 262 to 9 065 pairs were available, and for the “Max” or “8000” condition, 80% of all available pairs were used for training and 20% for testing. The encoded paratope-epitope pairs are processed for NLP processing by cutting the paratope and epitope in words of size 1 (degree-free) or size 2 (degree explicit) encodings while gaps were considered as one word (see Supplementary Figure 23A). Processed pairs were completed by adding a <START> and <END> word tokens, tokenized word by word, and padding. Words of size 1 in degree explicit encodings led to lower accuracies and we restricted ourselves to words of size 2.

#### Encoder-decoder

We leveraged a model based on Neural Machine Translation ^87^ to “translate” an epitope to a paratope and vice versa at motif, sequence, and aggregate levels. The architecture comprises two components: encoder and decoder with gated recurrent units (see Supplementary Figure 23). Within the decoder, a Bahdanau attention layer ^85^ was used to capture relevant input-side information necessary for the prediction of the output. Utilizing the context vector, the decoder generates each paratope or epitope motif/sequence character by character. The dataset was split into 80% training and 20% test set. An embedding layer of dimension 6 was used to learn numerical representations of the input. 516 cells in the GRU were taken, otherwise known as the length of the hidden dimension. The training procedure was carried out for 150 epochs with Adaptive Moment Estimation (Adam) optimizer ^118^ and was replicated ten times. 20 additional epochs were used where the loss function was letting the decoder predict the full output sentence instead of predicting the next word from the true previous word. The deep learning framework, `Tensorflow ver. 2.0` ^119^ was used.

#### Transformer

A transformer architecture was used as a comparison and implemented in `Tensorflow ver. 2.0.` Specifically, 8 heads, an embedding dimension of 16, and feed-forward layers of dimension 512 were used, together with a dropout rate of 0.1. The training dataset was separated into 40 batches of equal size and 250 epochs were used.

#### Shallow learning models for classification (including paratope-epitope prediction as a classification problem)

Naive Bayes (NB), Support Vector Machine (SVM), Random Forest (RF), and Logistic Regression (LR) models were defined as shallow models. All shallow learning models were parameterized with their baseline (default) parameters (see Table 1). We note that the shallow learning models, namely LR and SVM, yielded comparable performance (Supplementary Figure 11) in comparison to the single-layer neural network (Figure 3) and the performance may further be improved by hyperparameter optimization. Prior to training, the data were randomly split into training and test datasets (80:20), each amino acid was encoded as a one-hot vector (of size 20), consequently, each sequence was encoded as a one-hot vector of size 220 (11×20). Following training, the test dataset was used to obtain the model’s accuracy. The ML framework `Scikit-learn ver. 0.21.3` ^120^ was used to construct, train and evaluate the models. The comparison of ML architectures from Mason et al. ^39^ in Figure 3H, was performed using hyperparameters from their study instead of the ones used for classification in the rest of the manuscript (see *Ranking of ML architectures for sequence classification* below).

**Table 1 (refers to Methods, Figure 3 and Figure 5).** Parameters of the shallow learning methods. During training, unless stated otherwise, the default parameters for each model were used. We note that we did not perform any hyperparameter optimization. Therefore, the performance of these models may further be improved by additional steps of hyperparameter optimization.

| Shallow learning model | Parameters used |
| --- | --- |
| Naive Bayes (NB) | priors=None, var_smoothing=1e-09 |
| Support Vector Machine (SVM) | penalty="l2", loss="squared_hinge", dual=True, tol=0.0001, C=1.0, multi_class="ovr", fit_intercept=True, intercept_scaling=1, class_weight=None, verbose=0, random_state=None, max_iter=1000 |
| Random Forest (RF) | n_estimators=100, criterion="gini", max_depth=None, min_samples_split=2, min_samples_leaf=1, min_weight_fraction_leaf=0.0, max_features="auto", max_leaf_nodes=None, min_impurity_decrease=0.0, min_impurity_split=None, bootstrap=True, oob_score=False, n_jobs=None, random_state=None, verbose=0, warm_start=False, class_weight=None, ccp_alpha=0.0, max_samples=None |
| Logistic Regression (LR) | penalty="l2", dual=False, tol=0.0001, C=1.0, fit_intercept=True, intercept_scaling=1, class_weight=None, random_state=None, solver="lbfgs", max_iter=100, multi_class="auto", verbose=0, warm_start=False, n_jobs=None, l1_ratio=None |

#### Performance metrics

Accuracy is defined as 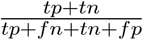 where *tp* is true positive, *tn* is true negative, *fp* is false positive and *fn* is false negative.

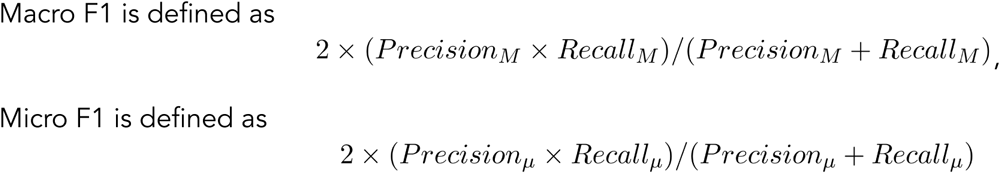

where macro averaging *M* is the per-class average and micro averaging *μ* is the cumulative average as defined in ^78^. Specifically, macro averaging splits the multi-class problem (*k* classes) into a binary classification for each class (*k_i_*), measures the performance metric for each class and finally averages the results from all classes (average of *k* metrics). In contrast, micro averaging treats the entire dataset as one aggregate and measures one single metric (instead of *k* metrics). Thus, micro-averaging is more sensitive to imbalance in the dataset since the class with a large number of samples dominates the average. Macro- and micro-averaging will return an identical metric when the number of samples in each class is identical (balanced dataset).

#### Integrated Gradients

(IG) ^80^ were computed with PyTorch ^121^ on the SN10 architecture to identify the importance of input dimensions towards the binder / non-binder prediction. The SN10 model (Figure 3) was trained on 40 000 sequences from dataset 1 for antigen 1ADQ_A (i.e., containing 50% binders and 50% non-binders to antigen 1ADQ_A). IG were only calculated on binder sequences from the training dataset (positive class), to identify which positions were predicted to be important for binding, or to reveal clustering of sequences based on their predicted IG. For each sequence, linear interpolations of 100 points between the one-hot encoded input and a baseline were created (we used the “zero” baseline, i.e., a zero vector with the same dimensionality as the encoded input). Gradients for each interpolation were computed with respect to the output of the neural network. The IGs are computed by averaging over all 100 gradients. Thus, each sequence is associated with a 11×20-dimensional IG matrix.

#### VAE

We used a standard VAE implementation ^122^ using fully connected layers, batch normalization between layers (for numerical stability and to allow for higher learning rates, we used the Adam optimizer with a learning rate of 1e-04), and a module for sampling from standard normal distribution. The architecture is described in Table 2. The encoder takes one-hot encoded 11-mers (220 input units) while the decoder contains 220 output units. We trained the model with 233 605 samples. The VAE optimizes Evidence Lower Bound (ELBO), which consists of reconstruction error and the Kullback-Leibler (KL) divergence to keep the posterior probability close to the latent prior. The reparameterization trick was used to allow backpropagation through non-deterministic nodes in the neural network. Specifically, our implementation uses negative log-likelihood for reconstruction loss and KL divergence as a closed formula, and was built using the PyTorch deep-learning framework. Regarding parameter beta, we tried the following (1.05, 1.1, 1.2, 1.5, 2.0). The purpose of parameter beta was to encourage learning disentangled representation where each dimension of the latent space (orthogonal directions) corresponds to a single variation factor in the input space ^123^. We noticed that while increasing the value of parameter beta, the reconstruction loss increased as well on average (resulting in lower reconstruction quality, also described by Dupont et al. ^124^), and did not observe any changes in the clustering. We, therefore, used beta=1.0.

**Table 2 (refers to Supplementary Figure 14):** VAE architecture layer by layer. The number of neurons in each layer is shown, with the type of activation function. As a comparison, other architectures with one less layer in both encoder and decoder were tested and yielded less clear delineation of clusters, written as [Encoder dense layer sizes (latent space dimensions used to represent mean and variance and first decoder layer size). Other decoder dense layer sizes]: [220-100-(50-50,40-40,30-30, or 20-20)-100-220]; [220-200-(50-50,40-40,30-30, or 20-20)-200-220]; [220-300-(50-50,40-40,30-30, or 20-20)-300-220]; [220-400-(50-50,40-40,30-30, or 20-20)-400-220]; [220-500-(50-50,40-40,30-30, or 20-20)-500-220]; [220-500-(50-50,40-40,30-30, or 20-20)-500-220] and [220-500-(20-20)-500-220].

| Encoder | Decoder |
| --- | --- |
| Dense (220) - BatchNorm1d - ReLU | Dense (40) - BatchNorm1d - ReLU |
| Dense (300) - BatchNorm1d - ReLU | Dense (100) - BatchNorm1d - ReLU |
| Dense (100) - BatchNorm1d - ReLU | Dense (300) - BatchNorm1d - ReLU |
| Dense (40) for mean and dense (40) for variance - BatchNorm1d | Dense (220) - Softmax |

#### ESM-1b embedding layer

In order to leverage biochemical attributes pre-learned on 250 million protein sequences, we added to the VAE the pre-trained ESM-1b transformer embedding architecture ^125^, using the default parameters to generate per-sequence latent space representations of 11-mer sequences.

#### UMAP visualization

The latent space of the VAEs was clustered using UMAP with the following parameters: n_neighbors_=(2, 5, 10, 20, 50, 100, 200), min_dist=(0.0, 0.1, 0.25, 0.5, 0.8, 0.99), n_components_=(2,3), metric=(“euclidean”,”correlation”).

### DLAB-VS pose classification pipeline

#### Generation of poses using Absolut!

During exhaustive docking, Absolut! enumerates all possible conformations (binding poses) of a 11-mer CDRH3 sequence to a predefined antigen, and calculates two energy components for each pose: the binding energy and the folding energy. Their sum is the total energy of the pose, which represents its probability of being stable. Instead of returning only the structures with optimal total energy (the most stable ones), we modified Absolut! to rank all the enumerated poses (6.8 millions on average during one exhaustive docking) based on their total energy, and to return the top N=500 poses with minimal total energy. This step replaces the docking step in DLAB-VS ^81^.

#### Selection of antibody-antigen pairs

In order to create a database of antibody-antigen poses for pose classification, we selected sequences with various energy thresholds to Absolut! antigens. For each antigen *A_i_* of the library, we randomly selected: i) 500 high-affinity 11-mers (top 1% affinity), 2 500 low-affinity 11-mers (top 1%–5% affinity), iii) 2 500 non-binders (bottom 95% affinity), and iv) 4 500 11-mers from the pool of sequences that were high affinity to any of the 142 antigens, as “other pairs”. We included these 11-mers irrespectively of whether they would also have high affinity to the considered antigen *A_i_*, to mimic the bias of the random pair matching strategy as used in PPI^82^. It is however possible to annotate them with Absolut! as false negatives prospectively during performance analysis if necessary. The numbers of high-affinity and “other” pairs were calibrated to the 1:9 ratio reported in DLAB-VS (thereby 500 and 4 500). We therefore created a database of 500 poses from 10 000 antibody-antigen pairs for each antigen in the library (700 M poses in total). For DLAB-VS, due to computation time constraints, we randomly picked 10 antigens for each simulation and only kept *GroupSize*=10 randomly selected poses per antibody-antigen pair from each antigen before dataset preprocessing. Up to 142 antigens and 500 kept poses per pair were available within the generated pose dataset.

#### Calculation of fnat and annotation of poses

The poses from low affinity pairs, non-binding pairs and other pairs were labeled with “L”, “N”, and “O” and were all associated with the class 0 (negative). Each pose from a high affinity pair needed to be compared to the target binding structure, that is, the energetically optimal pose for this pair (based on total energy, as used in the other figures of the manuscript). For a pose (the pose to be compared or the target binding structure), the set of binding pairs was enumerated (containing each time a residue in the CDRH3 and a residue in the antigen, that are neighbors in the lattice, i.e., interacting). The intersection of the two binding pairs was calculated (and represents the list of “conserved native contacts”). Subsequently, this number was divided by the total number of binding pairs in the target pose, thus ranging between 0 (no shared contacts) and 1 (all contacts from the target pose are present in the pose). Finally, depending on the positive and negative thresholds threshold_P_ = 0.5 or 0.7 or 0.9 or 1, and threshold_N_ = 0.1, any pose with fnat higher than threshold_P_ was labeled “P” and classified 1 (positive), while poses with fnat lower than threshold_N_ were labeled “I” (incorrect) and classified 0 (negative). Of note, poses with a fnat between the two thresholds are considered as a “no man’s land” and were discarded from the datasets, because the fnat is not directly mirroring the relative energy (binding or total energy) between the poses. In Absolut!, it would also have been possible to classify “P” and “I” poses based on an energy threshold but this information is not available experimentally at high throughput and the predicted Absolut!-based results would therefore not be transferable to application on experimental datasets.

#### Encoding of poses

Starting from a pose, only paratope and epitope residues were kept. The barycenter of paratope residues and epitope residues was calculated separately, then averaged as the “interaction center” (IT), and the full pose was translated to put the IC at position (2.5,2.5,2.5). Since the IC is a floating number, the translation was rounded randomly up or down for each direction to end up in integer positions, therefore similar poses are sometimes translated between each-other by a maximum of 1 grid step in each direction (this creates a data augmentation by translation without increasing the number of poses). Finally, only the integer positions [0;5] for each axis are kept, generating a 6×6×6 cubic lattice. The paratope and epitope residues are then separated in two 6×6×6 cubic lattices, as input to the CNN, and each AA in the lattices is converted into a one-hot encoding with 20 possible values, and empty positions are encoded as a 20-dimensional zero vector.

#### Preprocessing of training and test datasets

The list of *GroupSize*=10 randomly selected poses per antibody-antigen pair were gathered, annotated with their label (P, I, N, L, O) and class (0/1) from 10 randomly selected antigens. To avoid data leakage, it is important that the poses from the same antibody-antigen pair should not appear both in training and test datasets. Therefore, the poses were shuffled by groups of *GroupSize,* and the first nPosesTrain = 8000 poses after shuffling are assigned to the training dataset, and the next nPosesTest=2000 poses are assigned to the test dataset. Importantly, since there are much less positive poses than negative poses, and as performed in DLAB-VS, the positive poses are repeated in the training dataset to reach the number of negative poses (i.e., the training dataset is balanced with 50% of positive poses and 50% of negative poses altogether).

#### Data augmentation by rotation

In order to rotate poses, a uniform list of rotations are enumerated (Algorithm 4). In order to be independent on the axes (rotationally invariant), a list of uniformly distributed unit vectors (defined as the phi (=0 at the equator) and theta (=0 at the z axis) angles in spherical coordinates) on a sphere are enumerated according to the following formula ^126^, where resolution=1000 is the number of unit vectors on the equator (this number decreases at higher latitudes).

#### Algorithm 4

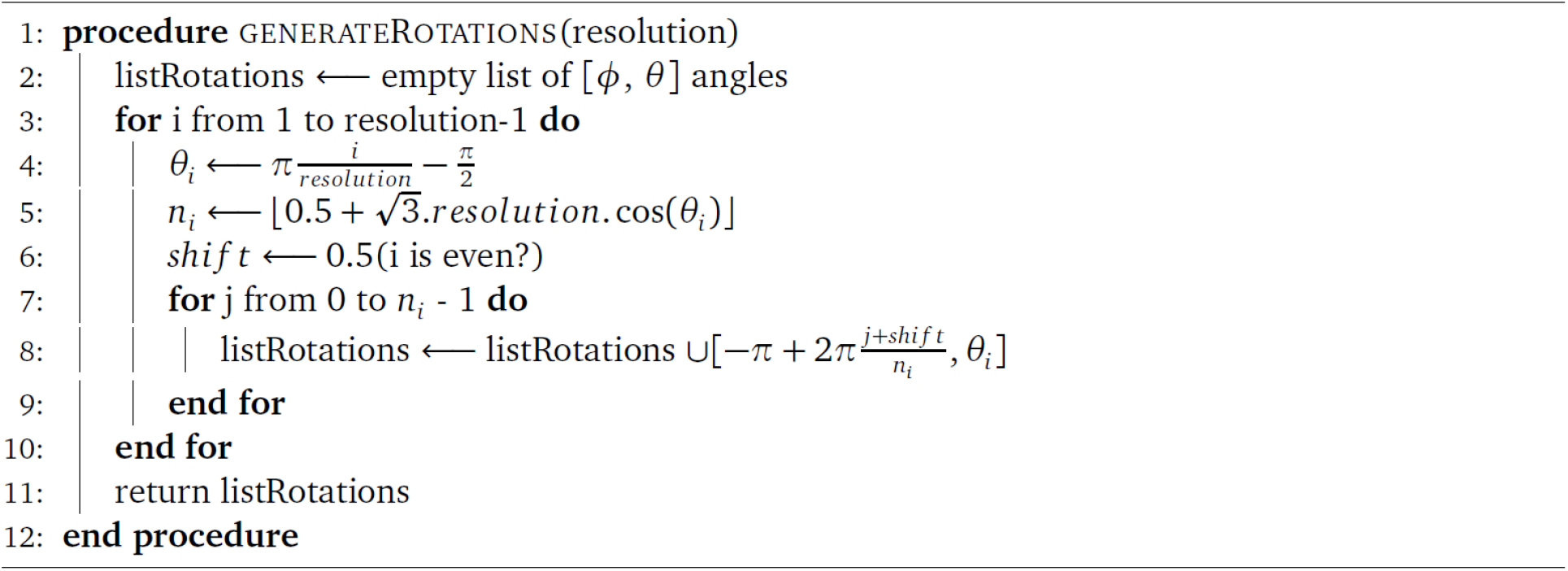
Function that generates a list of uniformly distributed unit vectors on a sphere.

When a pose needs to be rotated, during preprocessing, a random (phi, theta) rotation is picked from the list of rotations, and a rotation matrix is applied to each position of both paratope and epitope 6×6×6 cubic lattices that contained a residue (using the picked phi and theta, i.e., same rotation for both lattices). The new position of each residue is rounded up to the closest integer coordinate in each axis, and bounded within [0;5] to generate two new, rotated, paratope and epitope 6×6×6 cubic lattices. For each pose in the train or test datasets, n_Rotations_ were performed, and one of the rotations was performed with angles (phi=0, theta=0), preserving the original pose. This step multiplies the training and test datasets sizes by nRotations.

#### CNN architecture

The configuration of the CNN used for pose classification is shown in Table 3.

**Table 3.** Parameters of the CNN layers. *CNN architecture:* The input shape of the pose classification architecture is ([6×6×6×20], [6×6×6×20]). Two CNNs with the same architecture (but different parameters) process each cubic lattice in parallel. Each CNN has the following layers:

| Layer name | Layer type | Filter size | Kernel size | Padding type | Stride dimension | Kernel initializer | Activation | Batch normalization layer |
| --- | --- | --- | --- | --- | --- | --- | --- | --- |
| Layer 1 | 3D convo-lutional layer | 32 | 3 | Same (preserves the original 6x6x6 dimension) | (1,1,1) | Glorot uniform | Relu | Yes |
| Layer 2 |  | 64 |  | Valid (reduced dimensions at borders) |  | Glorot uniform |  |  |
| Layer 3 |  | 128 |  | Valid (reduced dimensions at borders) |  | Glorot uniform |  |  |
| Layer 4: Dropout layer: The output of the layer 3 of each CNN is flattened, and concatenated into a 256-dimensional feature vector followed by a dropout layer of value 0.2. |  |  |  |  |  |  |  |  |
| Layer 5: Dense layer that converts the 256 dimensions into one logit for the binary classification, with sigmoid activation function and kernel_initializer="glorot_uniform". |  |  |  |  |  |  |  |  |

#### Evaluation of model performance on a comparable basis

The inclusion of different types of negative poses generates datasets with different ratios of positive and negative examples, and also with a different distribution of poses with different labels (I,N,L,O) within the negative examples. It is therefore not straightforward to compare the performance of a trained model under different compositions of types of negative poses. When comparing trained models with the same type of negative poses (for instance in Figure 4C), the same distribution was kept for train and test datasets, except that positive poses were repeated during training to reach 50% positives. When comparing trained models under different dataset compositions, we decided to evaluate the trained models separately on test datasets with the same composition (Figure 4C, Supplementary Figure 13B-D), i.e., the models are evaluated in an out-of-distribution setting (when the distribution of negative poses in the training dataset was different than in the test dataset), that is, more “fair”. We consider only three test dataset compositions with predefined types of negative poses (“O”, “IO”, and “INLO” in Supplementary Figure 13B, C and D, respectively). Specifically, we first take the type of negative poses requested during training, then create a train/test split, then further process the test dataset by either removing unwanted types of negative poses, or by adding 2000 poses of a requested type of negative poses that were not in the training dataset composition. Finally, in both the training and test datasets, the positive poses were repeated to reach the 50% of positive poses, and therefore reach final balanced training and test datasets. This explains why the F1 score of models in Supplementary Figure 13A was low (the test dataset was unbalanced), while the F1 score of Supplementary Figure 13B–D was higher (the datasets were class-balanced, i.e., 50% positive poses). The summary of simulation parameters is given in Table 4

**Table 4.** List of parameters for a DLAB simulation. Altogether, a DLAB simulation can be parameterized according to the following parameters (the values used in the manuscript are given).

| Parameter name | Parameter value | Description |
| --- | --- | --- |
| nAntigens | 10 | Number of antigens |
| nPosesTrain | 8 000 | Number of poses in the training dataset (prior to data augmentation by rotation and prior to balancing by overrepresentation of positive poses) |
| nPosesTest | 2 000 | Number of poses in the test dataset (prior to data augmentation by rotation and prior to balancing by overrepresentation of positive poses) |
| nRotations | 1, 5, 20, 50, or 200 | Number of times of pose is repeated with a different random rotation |
| strategyNegatives | I, O, INO, ILO, INLO, or NL | Type of negative poses in the training dataset |
| strategyNegativeTesting | I, IO, or INLO | Type of negative poses in the test dataset |
| condition | one-hot or shuffled | one-hot encoding or shuffled data |
| groupSize | 10 | Number of retained poses for each antibody-antigen pair |
| fnatNegative | 0.1 | Maximum threshold of fraction of native (Fnat) for high-affinity antibody-antigen pairs to be classified as incorrect (classified as negative) |
| fnatPositive | 0.5, 0.7, 0.9, or 1 | Minimum threshold of fraction of native (Fnat) for high-affinity antibody-antigen pairs to be classified as positive |
| nEpochs | 10 | Number of epochs |
| batch_size | 2 000 | Number of samples in each batch |

### Ranking of ML architectures for high affinity antibody sequence classification on experimental and Absolut! datasets (refers to Figure 3H and Supplementary Figure 24)

#### Experimental and Absolut! data sets

We obtained CDRH3 sequences of antibody sequences with experimentally determined high affinity (n= 11300 positive sequences) and low affinity (n= 27539 negative sequences) to HER2 from Mason et al. ^39^. The sequences were of size 10AA and had an average LD of 8 to the original Trastuzumab sequences. Since the sequences were not especially similar to each other, we used the Absolut! dataset D1 (for 10 randomly selected antigens: 1ADQ_A, 1FBI_X, 2ARJ, 2B2X_A, 3B9K_A, 3BGF_S, 4AEI_A, 4CAD_C, 5B8C_C, 5BVP_I) containing ∼50 000 top 1% binding and bottom 99% non-binding 11-mers as synthetic dataset to compare the ranking of ML architectures between experimental and synthetic data. In order to check whether the predicted performance of ML methods was also valid in other data design settings with mutated sequences that are more similar to each other, in Supplementary Figure 24, we also Absolut!-generated mutated sequences (50 000 with 1 point-mutation, 25 000 with 2 point-mutations, and 25 000 with 3 point-mutations) around the CDRH3 sequence “CAPNLLFITTVVAPFDYW” that was a top 1% binding sequence to antigen 1ADQ_A. Mutated CDRH3 sequences above the top 1% threshold were considered as positive and the remaining were labeled as negative.

#### Data preprocessing

Analogously to Mason et al. ^39^, eight data preprocessing conditions were designed with different ratio of binding and non-binding sequences in the training dataset, by random sampling of 7 894 binders and from 7 237 (condition “A”) to 17 237 (condition “K”) by incrementing the number of negative sequences by 1 000 between conditions. The test dataset was generated with 1 622 binders and the same amount of non-binders, distinct from those in the training dataset. Therefore, evaluation metrics are balanced while the training dataset is more or less balanced depending on the condition.

#### ML architectures

We used the same ML methods and ML architectures as Mason et al. ^39^ (Table 5).

**Table 5.** ML methods and architectures. The sklearn package was used for shallow models (LSVM, SVM, LR, RF) and keras was used for the ANN, LSTM, and CNN architectures, using the following hyperparameters, taken from Mason et al.

| ML model/DL architecture | Parameters |
| --- | --- |
| LSVM | sklearn.svm.LinearSVC, Loss function = “squared_hinge”, Tolerance = 1e-4, C (Regularization) 1.0 |
| SVM | sklearn.svm.SVC, kernel=“rbf” (Gaussian RBF), degree=3, gamma=“scale”, tolerance=1e-3 |
| LR | sklearn.linear_model.LogisticRegression, solver=“lbfgs”, C=1e-4, penalty=“l2” |
| RF | sklearn.ensemble.RandomForestClassifier, n_estimators=150, criterion=“gini” |
| ANN | tf.keras.Sequential input: flattened one-hot encoding [220] Dense, units=70, kernel_initializer=“uniform”, activation=“relu” Dropout, rate=0.1 Dense, units=70, kernel_initializer=“uniform”, activation=“relu” Dropout, rate=0.1 Dense, units=70, kernel_initializer=“uniform”, activation=“relu” Dropout, rate=0.1 Dense, units=1, kernel_initializer=“uniform”, activation=“sigmoid” loss = “binary_crossentropy” |
| CNN | tf.keras.Sequential input: 2D one-hot encoding [10x20] Conv1D, filters=400, kernel_size=3 Conv1D, filters=400, kernel_size=3, strides=1, activation=“relu”, kernel_regularizer=None, bias_regularizer=None, padding=“same” Dropout, rate=0.5 MaxPool1D, pool_size=2, strides=1 Flatten Dense, units=50, activation=“relu”, kernel_regularizer=None, bias_regularizer=None Dense, units=1, activation=“sigmoid” loss = “binary_crossentropy” |
| LSTM | tf.keras.Sequential() input: 2D one-hot encoding [10x20] regularizer=tf.keras.regularizers.l2(1e-4) LSTM, units=40, return_sequences=True, bias_regularizer=regularizer Dropout, rate=0.1 LSTM, units=40, bias_regularizer=regularizer, return_sequences=True Dropout, rate=0.1 LSTM, units=40, bias_regularizer=regularizer, return_sequences=False Dropout, rate=0.1 Dense, units=1, activation=“sigmoid” loss = “mean_squared_error” |

\* For ANN, CNN and LSTM: optimizer = tf.keras.optimizers.Adam() nEpochs = 20 batch\_size = 16

### Quantification of data leakage in paratope-epitope prediction

In order to assess whether similar instances in the train and test datasets were responsible for the high accuracy of paratope-epitope prediction trained models in certain encodings (Figure 5B,C), we created new train and test datasets (for each encoding separately). Briefly, two similar paratopes matching the same epitope should preferably be both in the train or the test dataset, but not one in each dataset because the model would predict correctly for the test instance even by overfitting to the similar training instance. However, two similar paratopes matching a different epitope do not need to be grouped in one dataset, because they represent “difficult” instances to learn.

#### Creation of train and test datasets with reduced data leakage

First, the paratope-epitope pairs were balanced by keeping the same number of paratopes for each epitope. For visualizing paratope similarity (Supplementary Figure 21), 20 epitopes were randomly chosen, and 500 paratopes for each epitope were randomly kept. The LD between any pair of paratopes was calculated and UMAP was used to cluster the paratopes, while their machine epitope (label) was used as coloring. For separating train and test instances, for each epitope separately, 10 000 paratopes were kept and a UMAP between paratopes using their LD as metrics was performed. The clusters generated by the UMAP were then separated and annotated with an ID using HDBSCAN and each cluster was randomly assigned either to the train or the test dataset. In this way, similar paratopes will appear in the same cluster and be segregated into the train or the test dataset. However, similar paratopes matching different epitopes were treated independently since the clustering and separation was performed for each epitope separately, and such paratopes would sometimes be both in train dataset, or both in test dataset, or one in train and one in test datasets.

### Graphics

Plots were generated using the R package ggplot2 ^127^ as well as the python packages seaborn ^128^ and matplotlib ^129^. Plots were arranged using `Adobe Illustrator 2020` (Adobe Creative Cloud 5.2.1.441). Logo plots were made with R package ggseqlogo ^130^ and networks using Cytoscape 3.0.

### Hardware used

Computations were performed on the Norwegian e-infrastructure for Research & Education (NIRD/FRAM; https://www.sigma2.no) and a local server.

### Code and data availability

The Absolut! package is freely available on https://github.com/csi-greifflab/Absolut/ and the Absolut! database is available on https://greifflab.org/Absolut/.

## Acknowledgements

We acknowledge generous support by The Leona M. and Harry B. Helmsley Charitable Trust (#2019PG-T1D011, to VG), UiO World-Leading Research Community (to VG), UiO:LifeScience Convergence Environment Immunolingo (to VG, GKS, and IHH), EU Horizon 2020 iReceptorplus (#825821) (to VG), a Research Council of Norway FRIPRO project (#300740, to VG), a Research Council of Norway IKTPLUSS project (#311341, to VG and GKS), a Norwegian Cancer Society Grant (#215817, to VG), and Stiftelsen Kristian Gerhard Jebsen (K.G. Jebsen Coeliac Disease Research Centre) (to LMS and GKS). This work was not funded by Marie Skłodowska-Curie Actions while grant writing was supported by the German Arbeitsamt. This work was carried out on Immunohub e-Infrastructure funded by University of Oslo and jointly operated by GreiffLab and SandveLab (the authors) in close collaboration with the University Center for Information Technology, University of Oslo, IT-Department (USIT). We acknowledge Thérèse Malliavin (Institut Pasteur, Paris, France) for her comments and suggestions that helped in the analysis of the results, and Constantin Schneider, for helping us reproduce the DLAB-VS pipeline.

## Supplementary Figures

**Supplementary Figure 1 (refers to Figure 1A).**
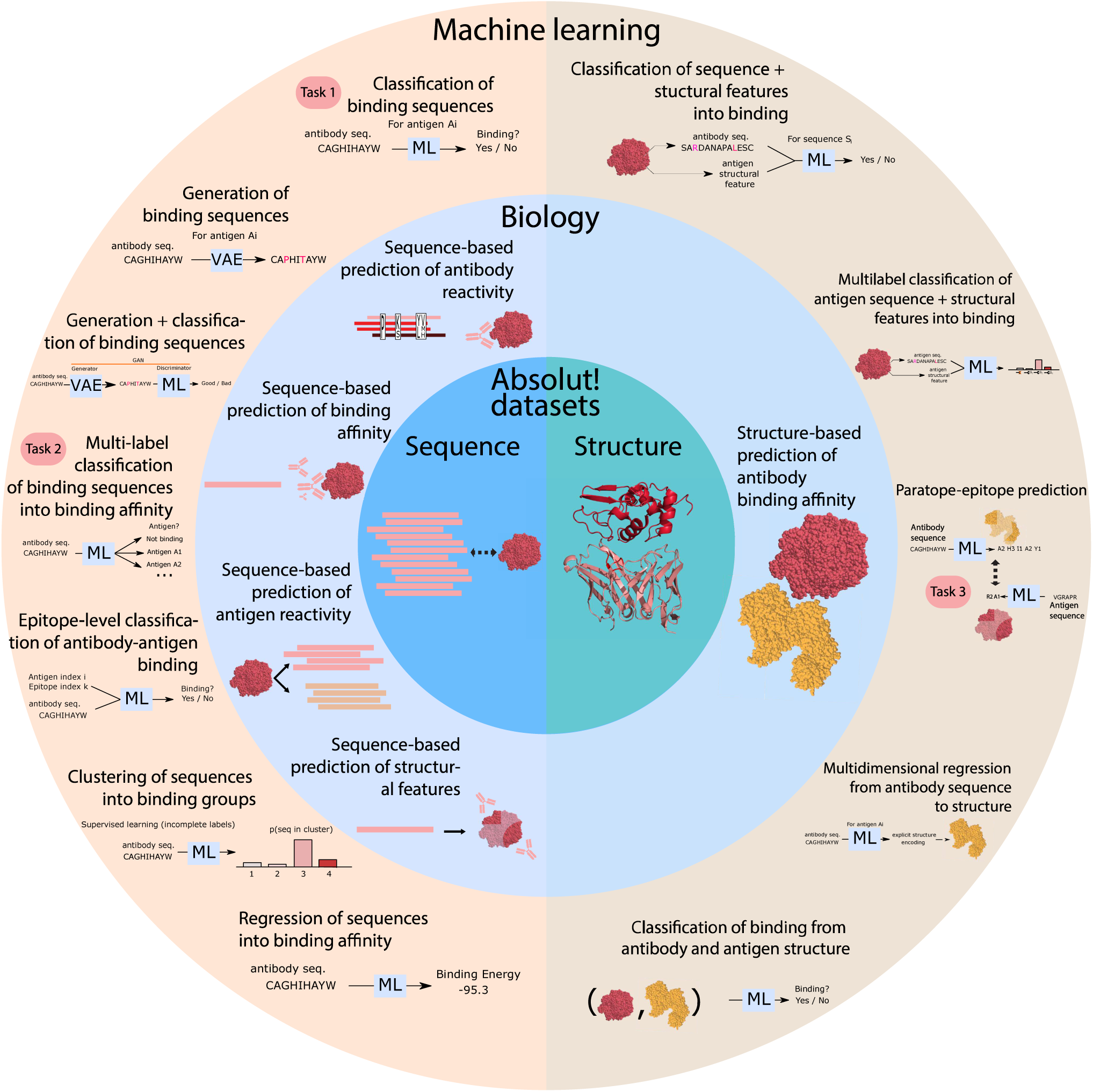
Immunological antibody specificity prediction problems formalized as machine-learning tasks for which Absolut! can generate synthetic benchmark datasets. Biological problems of antibody-antigen binding are very diverse and require translation into ML tasks performed on a specific type of experimental dataset. Datasets branch into two main categories: sequence- and structure-based (central ring). A large proportion of biological questions (middle ring) such as the prediction of antibody reactivity (to a defined antigen), binding affinity, antigen reactivity (to a defined antibody), and prediction of antibody-antigen structural features may be addressed (partially) by relying on sequence data alone, however, structural data contains the highest amount of information in terms of resolution and fidelity for the prediction of antibody-antigen binding (antibody and antigen structures are depicted in salmon and red respectively). Each biological problem can be converted into ML problems (outer ring) with a diverse panel of formulations, such as classification/generation of binding sequences, clustering of sequences into binding groups, multi-label classification of sequences into binding affinity class, prediction of structural features, and classification of binding from antibody and antigen structure. Most formulations typically fall into classification, regression, or paratope-epitope prediction (Figure 1A), and Absolut! is suited to generate *in silico* datasets for all of them.

**Supplementary Figure 2 (refers to Figure 1B).**
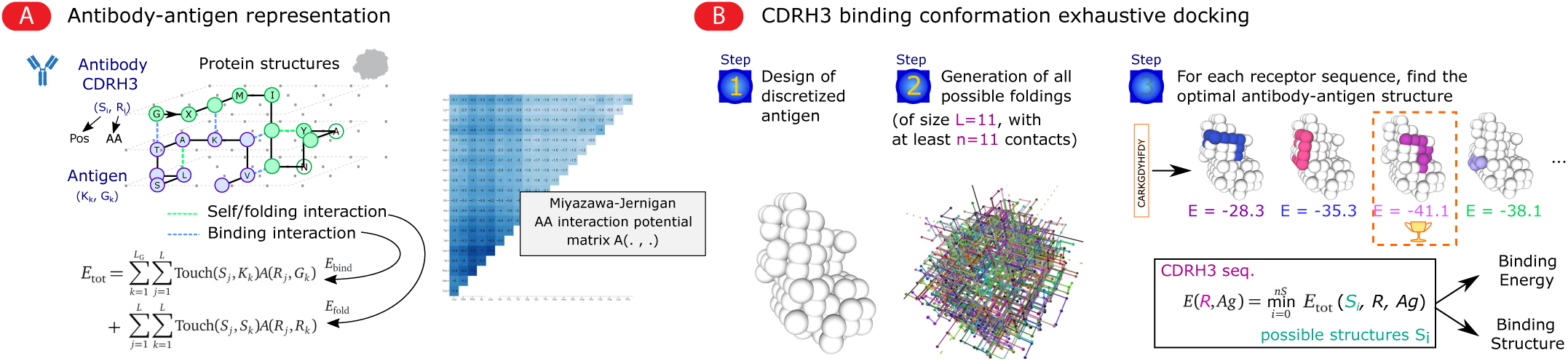
Representation of antibody-antigen binding and deterministic calculation of the binding structure by exhaustive docking. (A) Synthetic antibody-antigen binding structures are represented as lattice peptides whose neighboring amino acids interact (non-covalently) via the Miyazawa-Jernigan interaction potential ^68^. The stability of an antibody-antigen complex is assessed by its total energy (E_tot_) that sums two types of energies are defined from the interaction potential: the binding energy (E_bind_), summing the energy of bonds between the two proteins, and the folding energies (E_fold_), summing the energy of bonds within the antibody. The folding energy of the antigen is neglected since its structure is fixed and this energy is therefore constant. (B) Exhaustive docking: The energetically optimal binding conformation of an antibody chain sequence to an antigen is found by exhaustive search to minimize the energy of the binding complex (see Methods for details).

**Supplementary Figure 3 (refers to Figure 1B).**
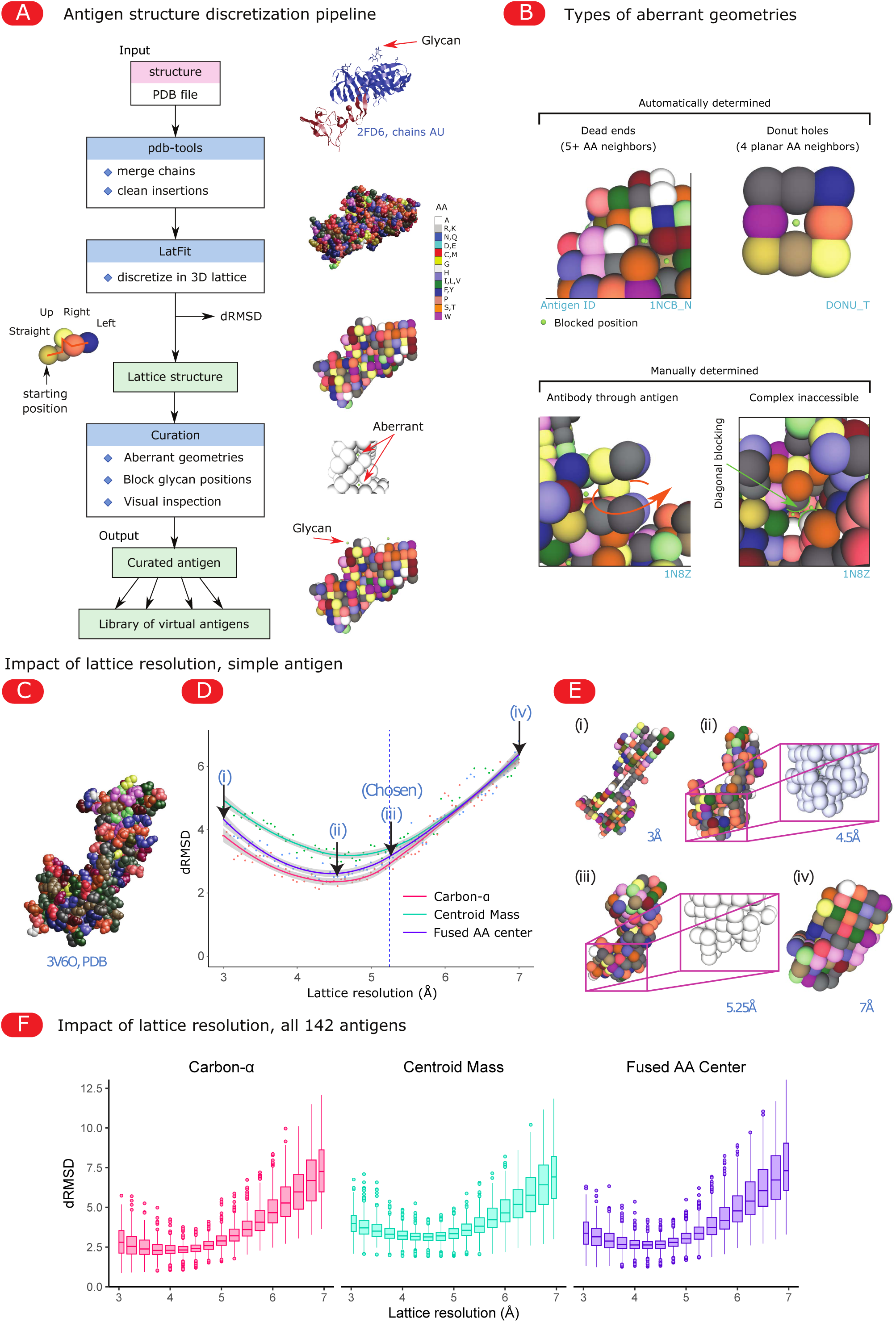
Steps of the antigen discretization pipeline and optimization of the lattice resolution. **(A)** Antigen discretization pipeline. Antigens are filtered for the chains of interest and removal of insertions by pdb-tools ^110^ and are further discretized into a 3D lattice using LatFit ^64^, which outputs a new PDB file with both lattice and real-world amino acid positions side-by-side. These positions are transformed into a lattice structure (see Methods), where each chain is defined by a starting position and a string of letters representing moves in the lattice (Straight, Up, Down, Left, Right). The presence of glycans on the protein is parsed from the original PDB and added at the vicinity of the designated residues that are closest to the original position (see Methods). Finally, aberrant geometries are removed (curated antigen, see Methods). (B) Definition of “aberrant geometries” that are blocked from antibody access during discretization. Dead-ends and donut-shaped holes are automatically detected and blocked recursively. Subsequently, more complex holes are manually determined by visual inspection, such as larger-than-donut holes that would allow the antibody to go through the antigen, or more complex pockets where the antibody loop would be “diagonally” blocked to enter. (C–F) Choice of the optimal lattice resolution for discretization. (C–E) Discretization of the antigen of PDB entry 3V6O with different lattice resolutions (distance between two lattice points), from 3Å to 7Å. (C) Original PDB structure of antigen 3V6O with amino acid coloring (shapely palette). (D) Quality of the discretization of 3V6O depending on the lattice resolution, measured as the dRMSD, quantifying the average distance between pairs of amino acids from the original PDB and their lattice position. The color represents the method of discretization: using the carbon alpha, the mass center of the centroid, or the fused center of the whole amino acid. (E) Result of discretization using the four lattice resolutions depicted in panel (D) using the fused center method, and showing the zoom on a concave area. (F) Discretization efficacy (dRMSD) of 130 antigens of the database, as a function of the lattice resolution and discretization method. In this manuscript, we opted for 5.25Å resolution (arrow C) in panel D, using a “fused amino acid center” method.

**Supplementary Figure 4 (refers to Figure 1C and Figure 2).**
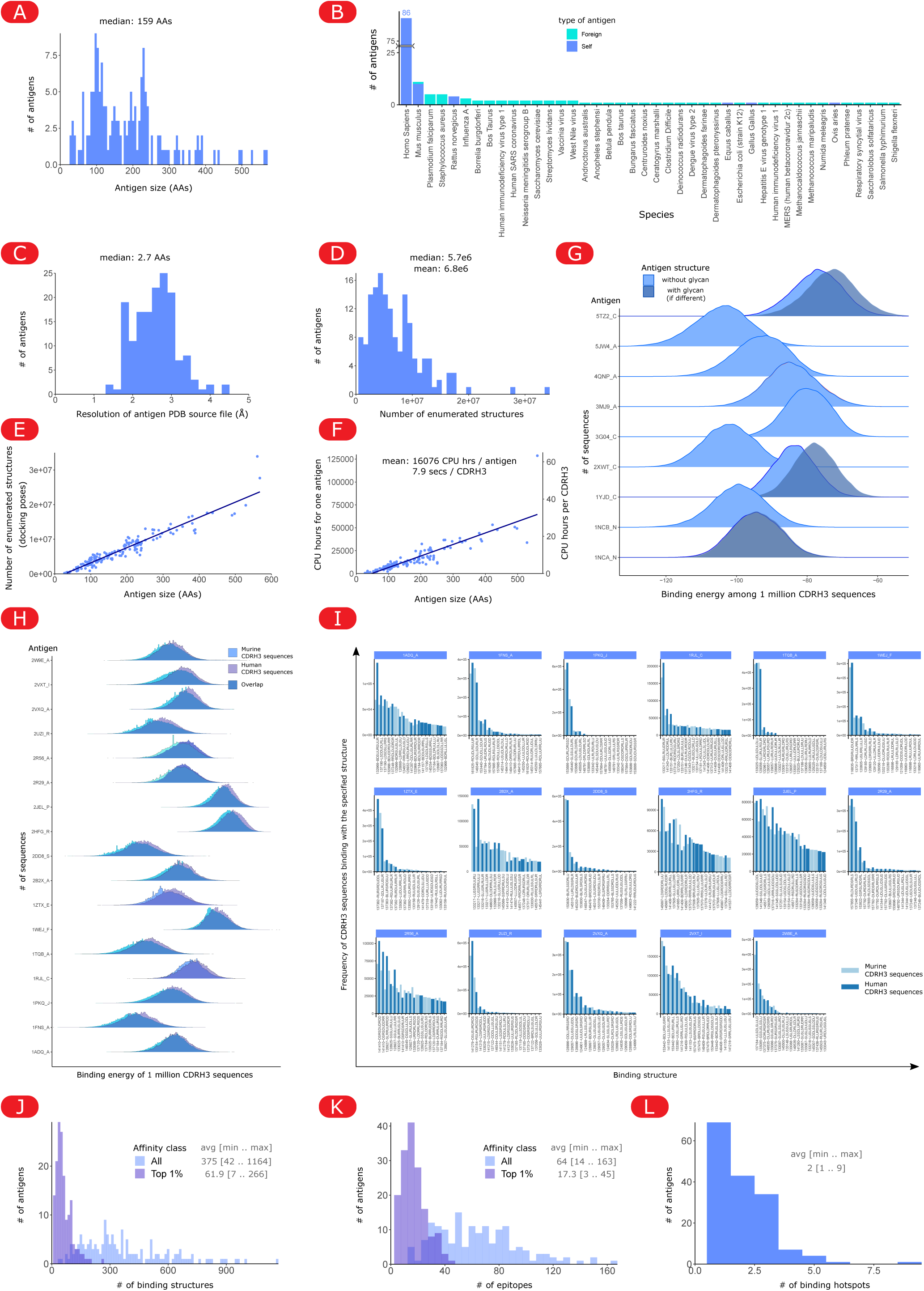
Computational complexity and binding diversity of the Absolut! database. (A) Amino acid size distribution of the 159 antigens (amino acid). (B) Species of origin of the antigens (blue: foreign and turquoise: self). (C) Resolution of the PDBs of the 159 antigens. (D-F) Computational complexity: (D) Distribution of the number of enumerated structures per antigen (tested docking poses for each CDRH3 affinity calculation). (E) Number of enumerated structures as a function of the antigen size. (F) Computational resources needed to generate the Absolut! database. Left axis: CPU hours to calculate the 6.9 million CDRH3 sequences for each antigen (as function of the antigen size). Right axis: average time to compute the affinity of one CDRH3 sequence to an antigen depending on its size. (G) Effect of glycans on binding profiles. Distribution of affinities of 1 million murine CDRH3 to antigens that contain glycans (See Table S1), when calculated for binding with (gray) or without (blue) these glycans. The gray curve is only shown when it is different from the blue one. (H,I) Affinities and binding conformations between 1 million murine CDRH3 and 1 million human CDRH3 sequences share the same ranges of affinities. (H) Distribution of binding energies (affinity). Human and murine sequences follow similar distributions of affinities for each antigen. (I) Distribution of the top 15 most used binding structures for each antigen. Human and murine CDRH3 sequences share the same usage of binding structures. Structures with only one color denote those that were not in the top 15 most used structures in the respective other category (mouse or human). (J–L) Structural diversity of binding for the CDRH3 sequences (either high affinity purple, top 1% or all sequences, blue) to each antigen. (J) Number of distinct antibody-antigen binding structures. (K) Number of epitopes. (L) Number of binding hotspots.

**Supplementary Figure 5 (refers to Figure 1).**
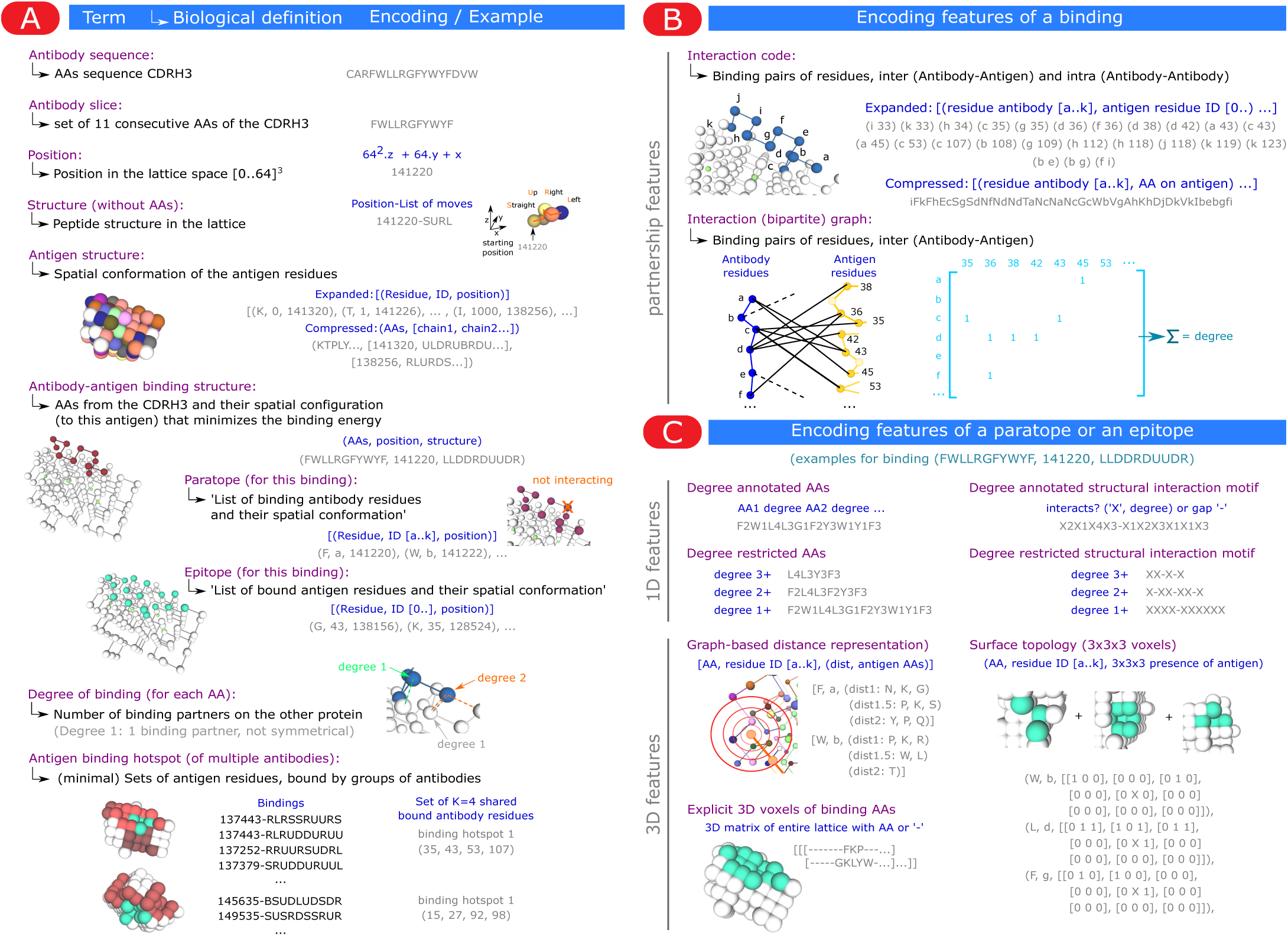
*In silico* formalization of antibody-antigen binding features and their encodings for ML. **(A)** Definitions. An antibody *slice* refers to 11 consecutive amino acids of the CDRH3, which is the size used to compute binding (see Methods) ^67^. Amino acids can only occupy integer 3D lattice *positions* encoded as a single integer number. A *peptide structure* is defined as a starting position and a list of moves in space, without knowing which amino acids are in this peptide. An *antibody binding* is then represented as the best spatial conformation of a slice that minimizes its energy (according to the interaction of neighboring lattice residues from the Miyazawa-Jernigan potential, see Methods and Supplementary Figure 2), and is stored as the amino acids with their structure in 3D. The *paratope* refers to the spatial configuration of only the antibody amino acids participating in the interaction, while the *epitope* is the spatial configuration of the antigen amino acids that are involved in the binding. Therefore, we define “paratope” and “epitope” as referring to the full structural information of the binding from either side. In a binding structure, an amino acid’s *degree of binding* refers to its number of binding partners on the opposite protein. Note that amino acids of degree 2 on the antibody can bind to residues of degree 1 on the antigen (residue degree asymmetry). The preferential binding of many antibodies to certain regions of the antigen can be analyzed by clustering the epitope residues as *“antigen binding hotspot”*, using a minimal set cover algorithm (see Methods) **(B)** Representations of the binding interface. The list of binding partners or the bipartite graph of interaction can conveniently describe some properties of the binding interface, while losing the spatial configuration of the epitope or paratope (surjective mapping). **(C)** Encodings of the paratope or epitope. Paratope and epitope may be represented with different resolutions, to represent the type of available experimental information, or as encoding features for ML pipelines. The binding amino acids can be annotated with their binding degree, or as “structural interaction motifs” ^5^ that only encode binding positions and gaps. The 3D environment of each amino acid may be encoded as a graph of neighbors for each range of distances ^71^, or as explicit presence of amino acids around (local surface topology). Finally, the full conformation of a paratope or epitope can be provided as a 3D matrix of voxels (3D cubic units) describing the type of amino acid at each position in the lattice, as used for protein docking strategies ^47^.

**Supplementary Figure 6 (refers to Figure 2C).**
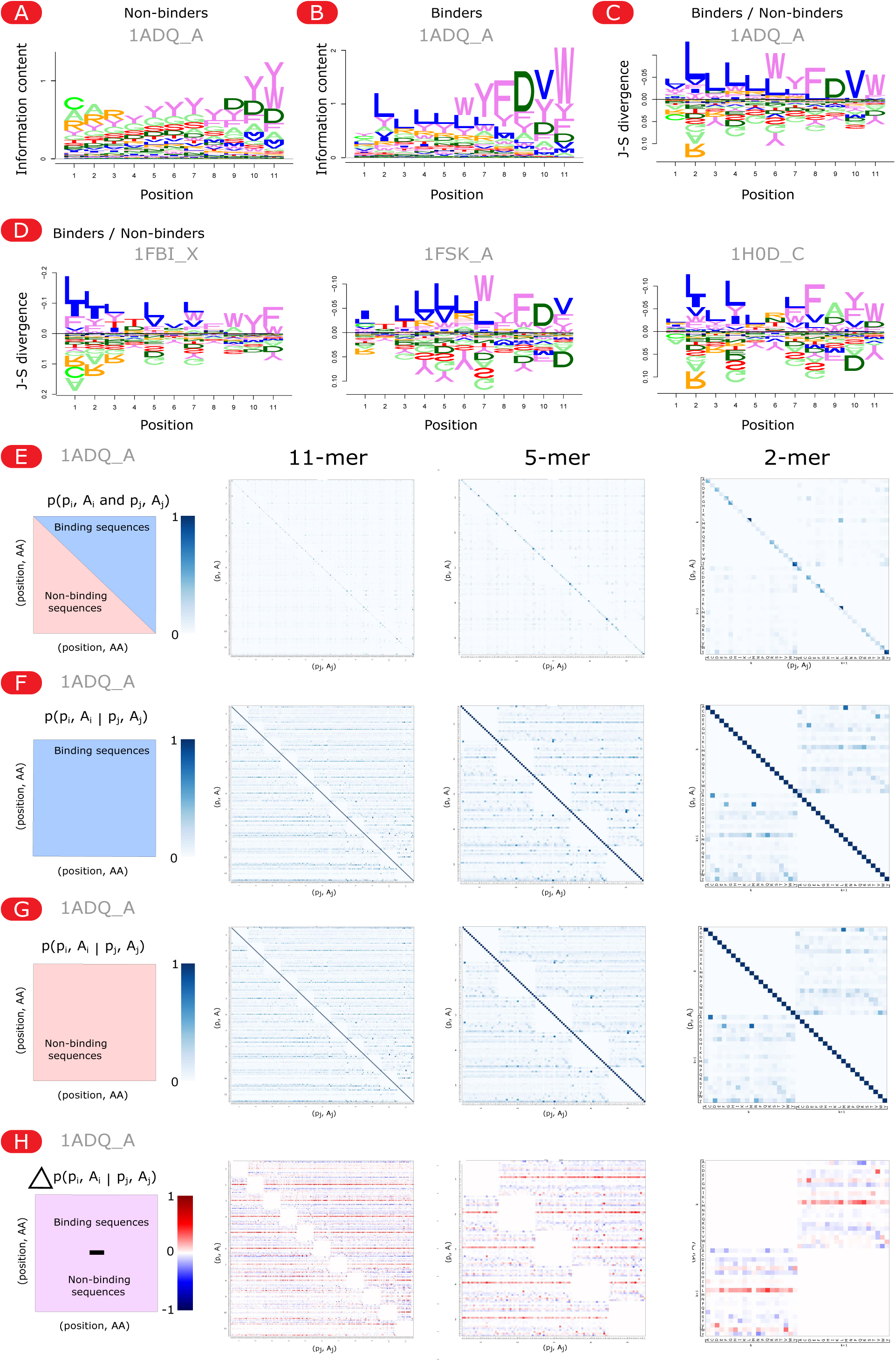
Positional amino acid frequencies and positional dependencies of binder and non-binder CDRH3 sequences to antigen 1ADQ_A. The binding CDRH3 sequences to antigen 1ADQ (as an example) were assessed for positional dependencies. (A–C) Amino acid positional bias between binder (from the top 1%) and non-binder (from the bottom 99%) sequences (11-mers) to 1ADQ_A (see Figure 2B for details). For each CDRH3 (binder or non-binder), its 11-mer with highest affinity was taken to build the aligned logo plot. (A) Positional amino acid usage is shown separately for non-binders, representing the sequence bias of CDRH3 sequences generated by VDJ recombination, as seen by the “CAR” sequence when the 11-mer is close to the beginning of the CDRH3 or the DYW sequence when the 11-mer lies close to the sequence’s end. (B) Positional amino acid usage of non-binder sequences, showing remaining biases from CDRH3 sequences but with a preferential usage of the end of the CDRH3 as seen by the DYW sequence. (C) Comparison of binder and non-binder positional amino acid usage shown as the Jensen-Shannon divergence between the distribution of amino acids at each position between binders and non-binders. (D) Comparison of binder and non-binder positional amino acid usage analogous to panel C for three different antigens, 1FBI_X, 1FSK_A and 1H0D_C. (E–H) The positional dependencies are further illustrated by the conditional probabilities of having an amino acid at a position knowing the amino acid at another position, assessed on all subsequences of size 2 (right), 5 (middle) or 11 (left) amino acids of the binders and non-binders. (E) The simultaneous appearance p(p_i_, A_i_ and p_j_, A_j_) of two amino acids A_i_ and A_j_ at two different positions p_i_ and p_j_ are quantified in binders and non-binders separately. (F–G) The conditional probability p(p_i_, A_i_ | p_j_, A_j_) to observe a certain amino acid A_i_ at a position p_i_ knowing the presence of another amino acid A_j_ at a position p_j_ is shown, which normalizes the correlations (simultaneous appearance) by the amino acid composition difference between binders and non-binders shown in panel (C). Residues A_i_ (y axis) that show consistently higher conditional probability across all positions (blue horizontal lines in F,G and blue or red horizontal lines in H) reveal only a higher amino acid usage of A_i_ while local colored points show complex positional dependencies. (H) The difference Δp(p_i_, A_i_ | p_j_, A_j_) = p(p_i_, A_i_ | p_j_, A_j_) (binder) - p(p_i_, A_i_ | p_j_, A_j_) (non-binder) between the conditional probabilities in binders and non-binders reveals binder-specific positional dependencies. The right panel (2-mer) is the one shown in Figure 2C.

**Supplementary Figure 7.**
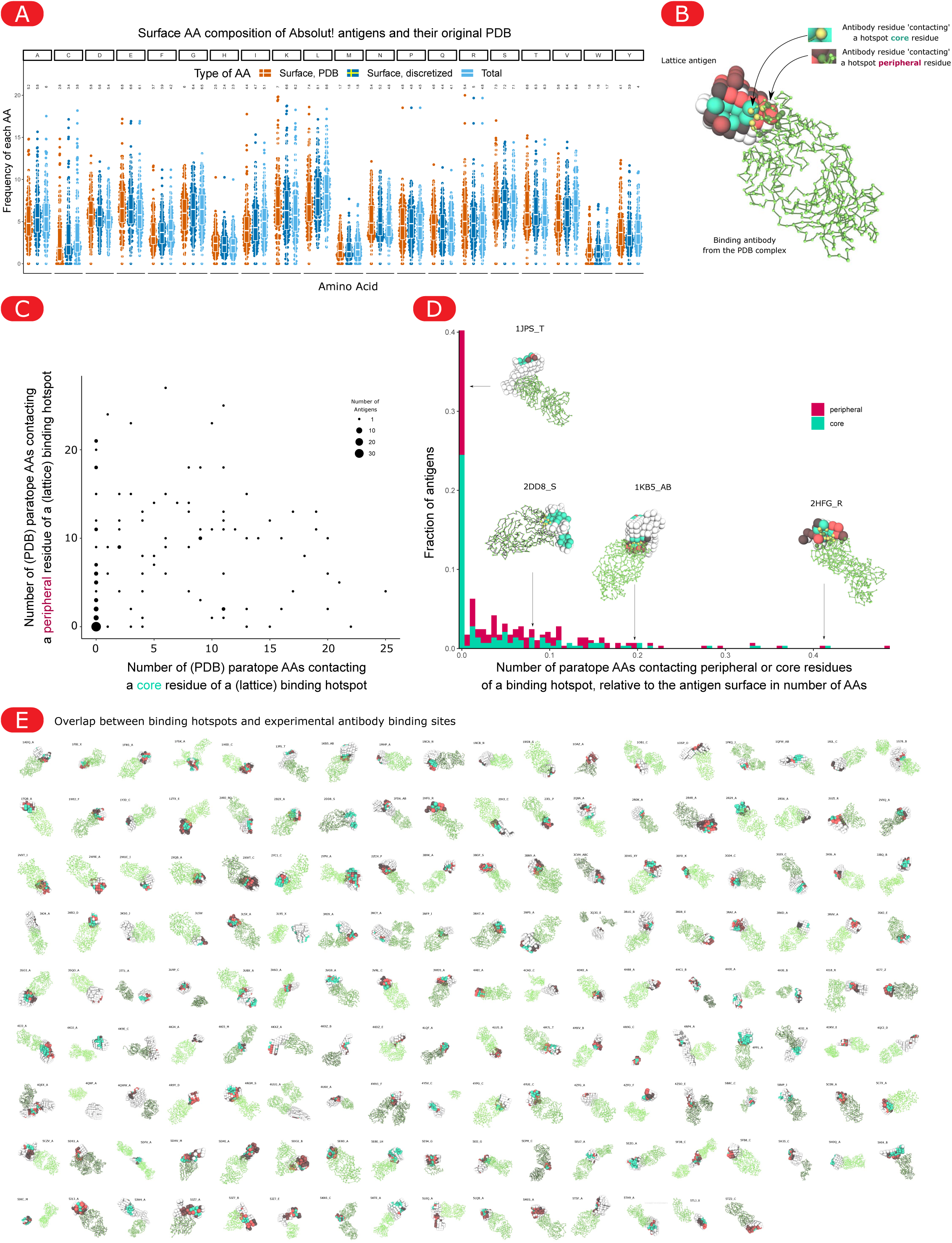
Comparison of Absolut! antibody-antigen binding structures with experimental crystal structures. **(A)** Surface amino acid composition of discretized antigens, compared to the experimental amino acid distribution in their original PDB file, or to the entire amino acid composition (not only surface, which is therefore identical between the PDB and lattice antigen). **(B)** Each antigen was built from PDB structures containing both the antigen and at least one binding antibody. Description of the overlap between the experimental antibody binding structure in the original PDB, and the predicted binding hotspots (see C). For a PDB antibody-antigen binding structure, paratope residues in the PDB, contacting a shared binding hotspot position (core, green) or a peripheral residue of a binding hotspot (dark red) in the lattice can be quantified. A PDB paratope residue is considered as contacting the antigen if it lies within a radius of 8Å to a lattice antigen residue, to mimic the lattice resolution of 5.25Å and the antigen discretization error RMSD around 3.5Å (Supplementary Figure 3). The different possible hotspots are all “merged”, meaning that in the lattice, by definition of the hotspots, each structure of a binder sequence in the Absolut! dataset was binding at least 4 green positions in one of the hotspots. **(C)** The number of PDB paratope residues contacting a core or peripheral residue of a binding hotspot were quantified. **(D)** Distribution of the number of paratope residues contacting a core and peripheral (side) epitope residues normalized by the number of amino acids at the surface of the lattice antigen. 75% of PDB antigen-antibody complexes were in contact with the lattice hotspot core residues identified from the 6.9 million murine CDRH3 sequences, while the remaining structures show a high variation of overlap, from few paratope residues (relative to the total antigen surface) such as antigen 2DD8, up to a strong overlap in the case of antigen 2HFG. When inspecting peripheral residues of binding hotspots, 85% of antibody-antigen complexes overlapped with peripheral hotspot residues. **(E)** Experimental PDB antibody binding versus predicted hotspots for all the Absolut! antigen-antibody complexes. In a few cases, the PDB antibody was binding to another protein chain than the one of the discretized antigen, in which case the PDB antibody and discretized antibody chain are not shown.

**Supplementary Figure 8 (refers to Figure 2E).**
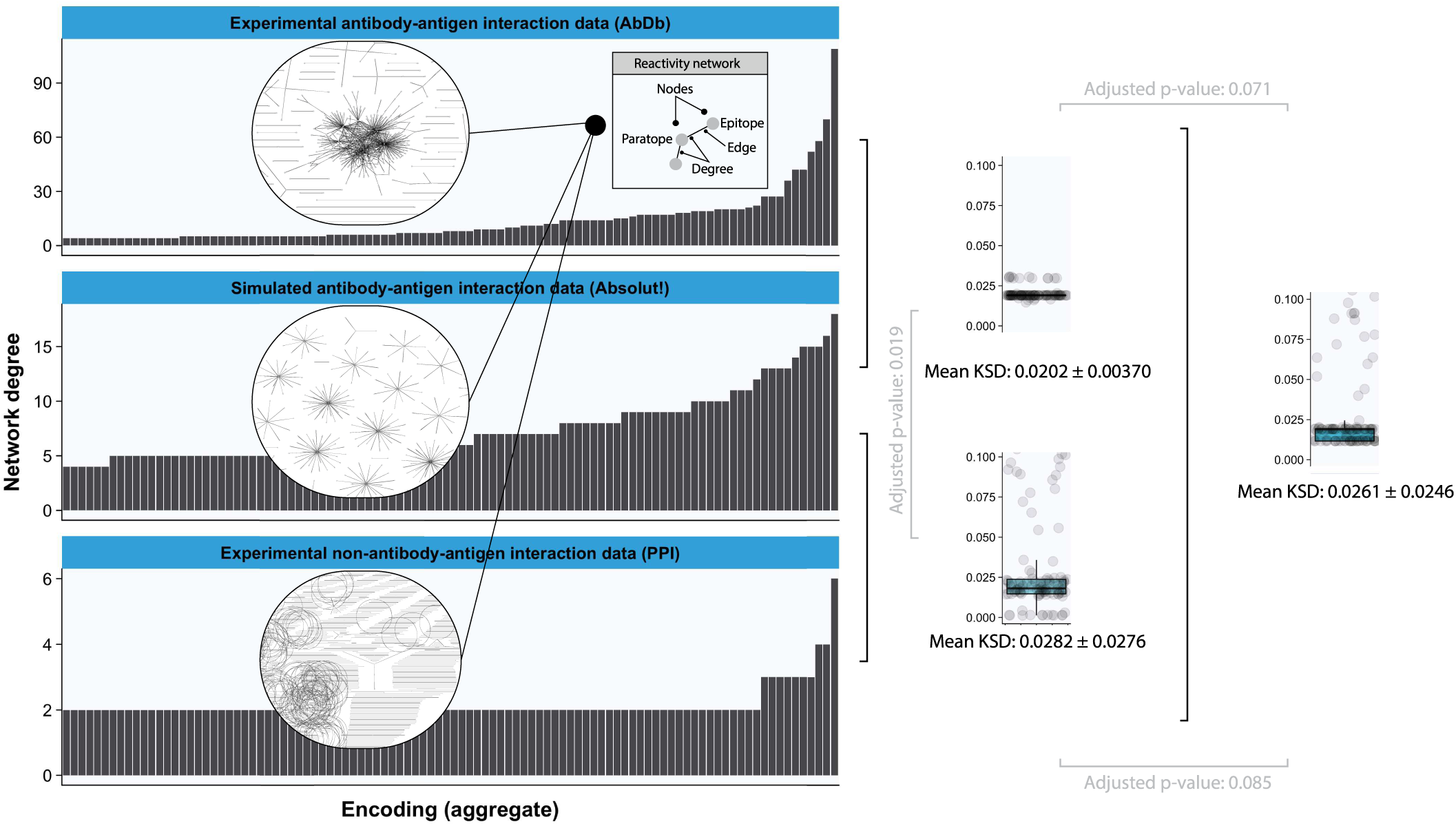
Reactivity networks of experimental (AbDb) and simulated antibody-antigen interaction data (Absolut!) are similar whereas the reactivity network of experimental non-immune protein interaction data (PPI) differs from both experimental and simulated immune interaction data. To compare the experimental antibody-antigen complexes (AbDb; data preparation and preprocessing were done as described in ^5^), Absolut! antibody-antigen complexes (Absolut!), and PPI (non-immune protein-protein interaction) complexes, we subsampled randomly the Absolut! and PPI datasets to match the size of the experimental dataset (n_pairs_=7 353 paratope-epitope pairs; n_subsampling_=100, non-overlapping subsampling), constructed the reactivity network (each paratope is connected to its epitope by an edge, see inset) for each subsampled dataset, plotted the resulting network degree distributions (here shown only for one subsample and for the top 200 aggregate encodings; see Figure 5 for detailed explanation on aggregate encoding) and calculated the KSD between empirical distribution functions of the subsamples. We compared the KSD distances in three groups (group 1: KSD between experimental vs Absolut! subsamples, group 2: KSD between experimental vs PPI subsamples, group 3: KSD between Absolut! and PPI subsamples) using pairwise paired t-tests and corrected for multiple testing using the Bonferroni method. We failed to reject the null hypothesis that the average KSD of network degree distributions are equal in group 1 (experimental vs Absolut!) vs group 2 (experimental vs PPI) and in group 3 (PPI vs Absolut!) and group 2 (experimental vs PPI) (adjusted p-values are 0.071 and 0.085, respectively). But we detected a statistical difference in group 3 (PPI vs Absolut!) vs group 1 (experimental vs Absolut!) (adjusted p-value 0.019).

**Supplementary Figure 9 (refers to Figure 3).**
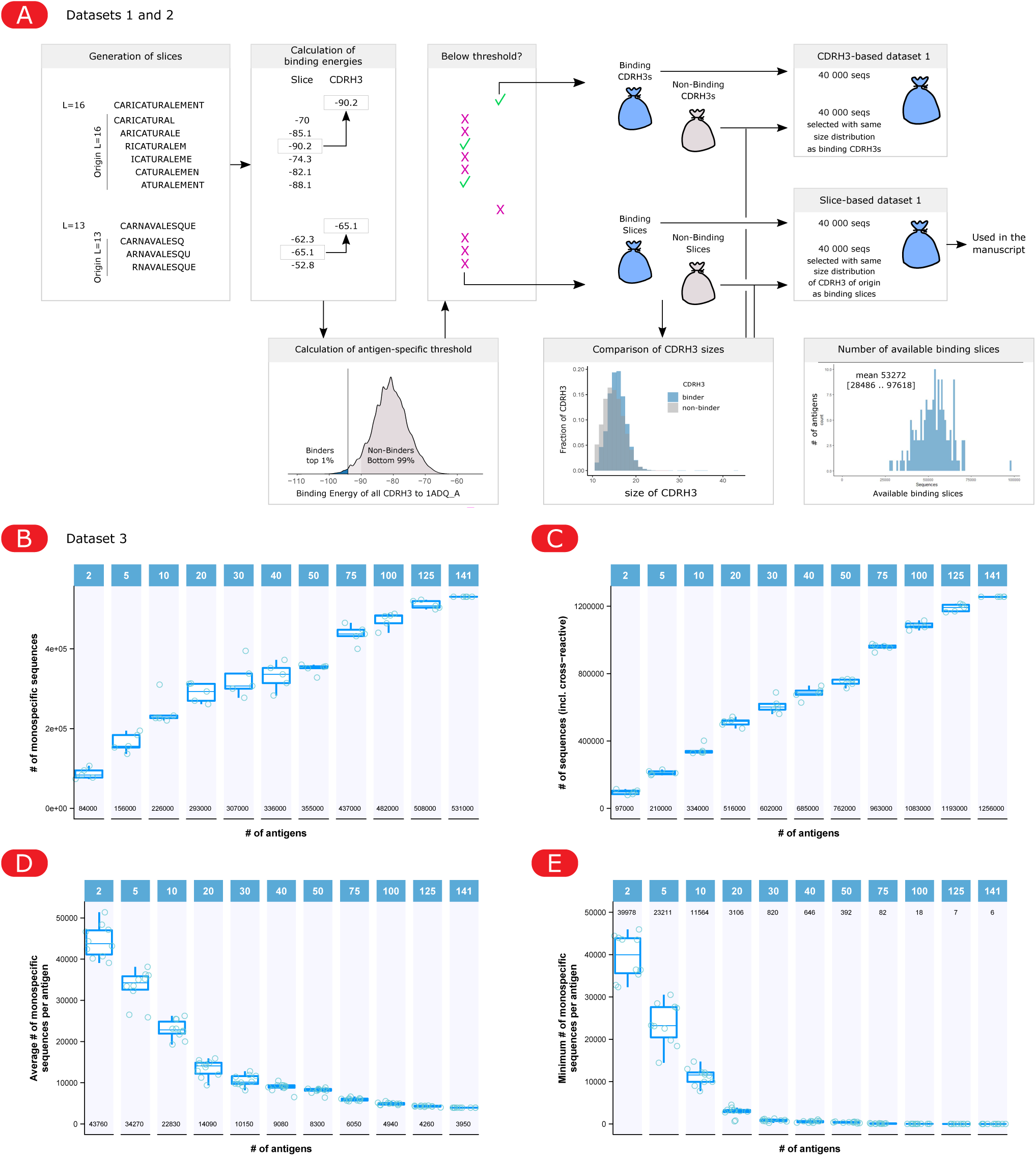
Generation and properties of classification datasets 1, 2 and 3. (A) Pipeline for the generation of Dataset 1 and 2 for a chosen antigen (see Methods). 11-mer slices were generated from the CDRH3 sequences and assessed for binding affinity to the antigen. The lowest energy among slices defines the CDRH3 energy. A (top 1%) binding threshold was calculated and each CDRH3 (or each slice) is labeled with a “binder” or “non-binder” tag according to the CDRH3-defined top 1% threshold. The binding CDRH3s have a bias to be longer (and the binding slices tend to stem from longer CDRH3s), which creates an amino acid composition bias between binder and non-binder CDRH3s (or slices), due to the frequent “CAR” sequence in the beginning of CDRH3s. Therefore, for building a CDRH3-based or slice-based-dataset 1, 40 000 binders are taken and 40 000 non-binders are sampled that follow the same CDRH3 size distributions as the selected binders. For dataset 2, an additional 5% threshold is defined per antigen to take non-binders within the lowest 1% to 5% energies. (B–E) Number of available sequences for Dataset 3 as a function of the number N of considered antigens. (B) Number of total monospecific sequences to one of the N antigens (of note, these sequences can be cross-reactive to antigens other than the N selected ones). (C) Number of total sequences (including cross-reactive sequences within the N selected antigens) to one or more of the N antigens. (D) Average and (E) minimum number of monospecific sequences recognizing each antigen (i.e., belonging to each class).

**Supplementary Figure 10 (refers to Figure 3).**
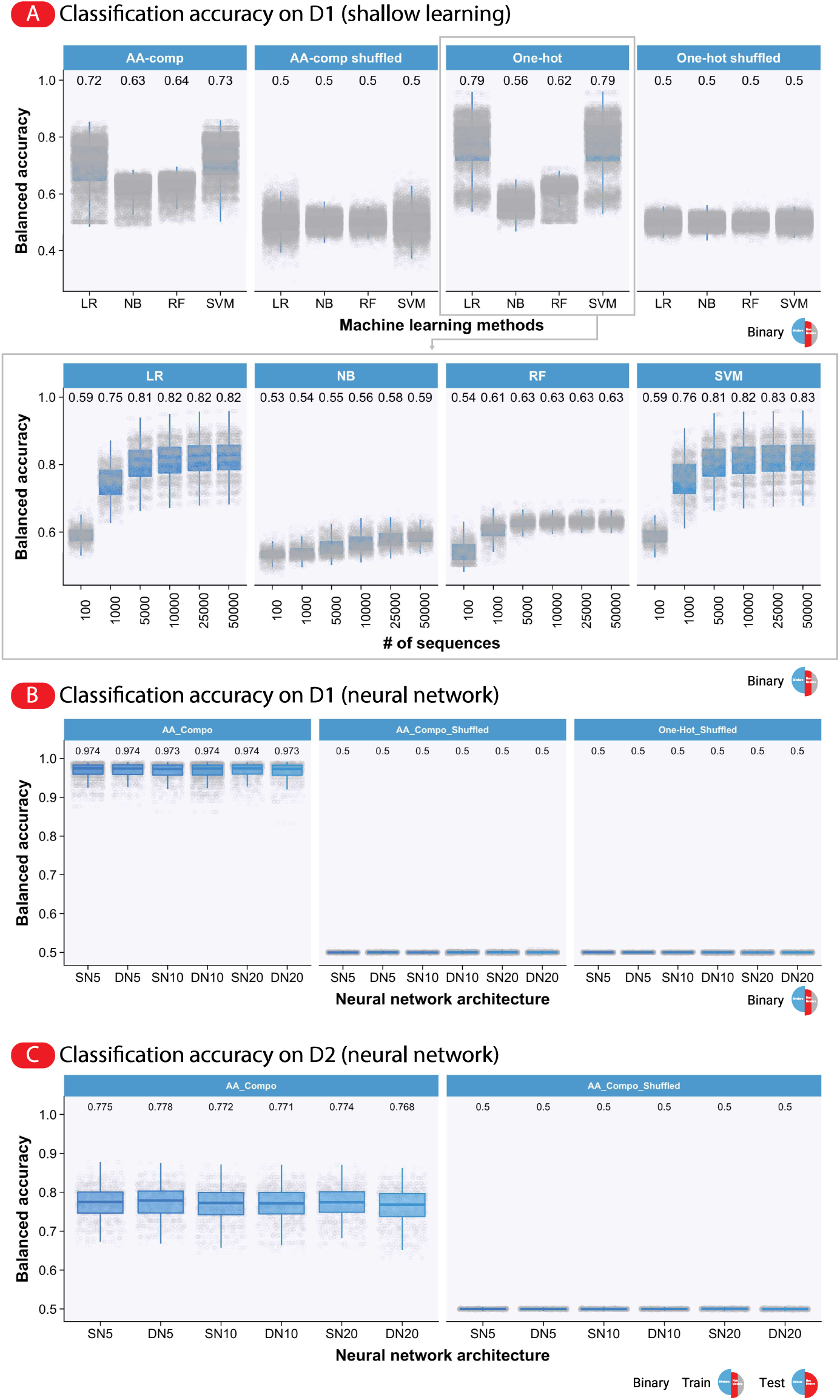
Classification accuracy on D1 and D2 for amino acid composition (AA-comp) and one-hot encodings. (A) Shallow learning. In addition to one-hot encoding, we used the frequency of the amino acids (AA-comp) to encode CDRH3 sequences and as a control we trained the models on label-shuffled datasets (shuffled). Binary classification median balanced accuracy (shuffled accuracy) ranged between 0.64–0.73 (0.5) and 0.56–0.79 (0.5) for AA-comp and one-hot encoding, respectively. Maximum accuracy (0.83) was obtained by SVM trained on the largest training dataset (50 000 sequences). Of note, ML methods were not optimized with respect to hyperparameters. (B) AA-comp encoded neural network binary classification median balanced accuracy ranged between 0.816–0.82 and shuffled median balanced accuracy for one-hot encoding 0.5 (see Figure 3 for accuracy values of models with one-hot encoding). (C) Median classification accuracy of neural network trained on D1 and tested on D2 for AA-comp ranged between 0.768–0.778 (shuffled accuracy at 0.5). For each antigen, model, encoding and n_CDRH3_training_, training and test were replicated 10 times.

**Supplementary Figure 11 (refers to Figure 3).**
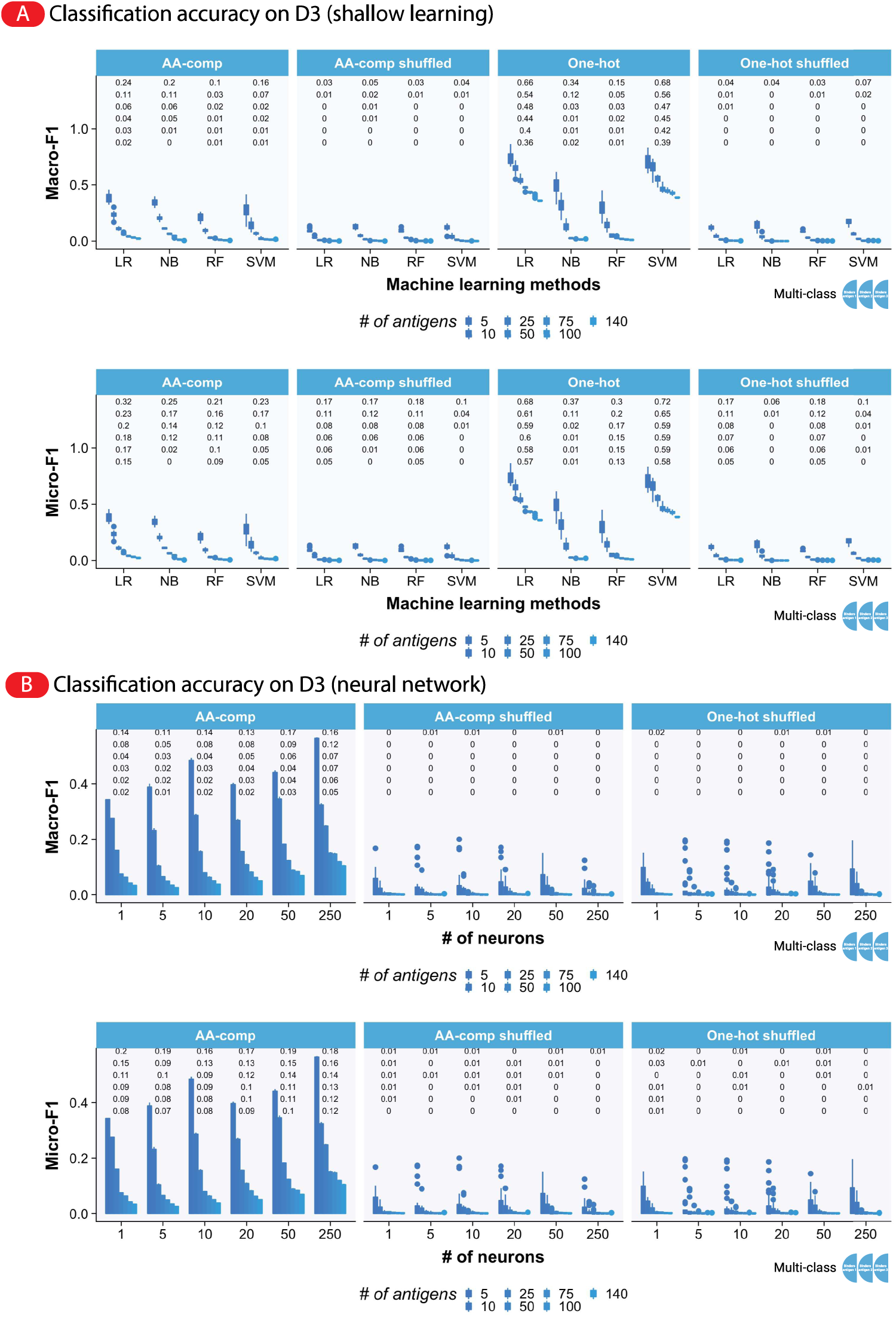
Classification accuracy on D3 for amino acid composition (AA-comp) and one-hot encodings. Accuracy of shallow architectures, shuffled control and amino acid composition encoding are assessed on D3 generated for different antigen numbers as described in Figure 3. (A) Shallow learning. Macro F1 multi-class classification median accuracy (shuffled accuracy) ranged between 0–0.24 (0–0.05) and 0.01–0.68 (0–0.07) for AA-comp and one-hot encoding respectively. Micro F1 multi-class classification median accuracy (shuffled accuracy) ranged between 0–0.32 (0–0.18) and 0.01–0.72 (0–0.18) for AA-comp and one-hot encoding respectively. (B) Neural network. Shuffled Macro F1 multi-class classification median accuracy ranged between 0–0.01 and 0.02 for AA-comp and one-hot encoding respectively. Shuffled Micro F1 multi-class classification median accuracy ranged between 0–0.01 and 0.03 for AA-comp and one-hot encoding respectively. See Figure 3 for unshuffled accuracy. For each model, encoding and # of antigens, training, and test were replicated 10 times.

**Supplementary Figure 12 (refers to Figure 3).**
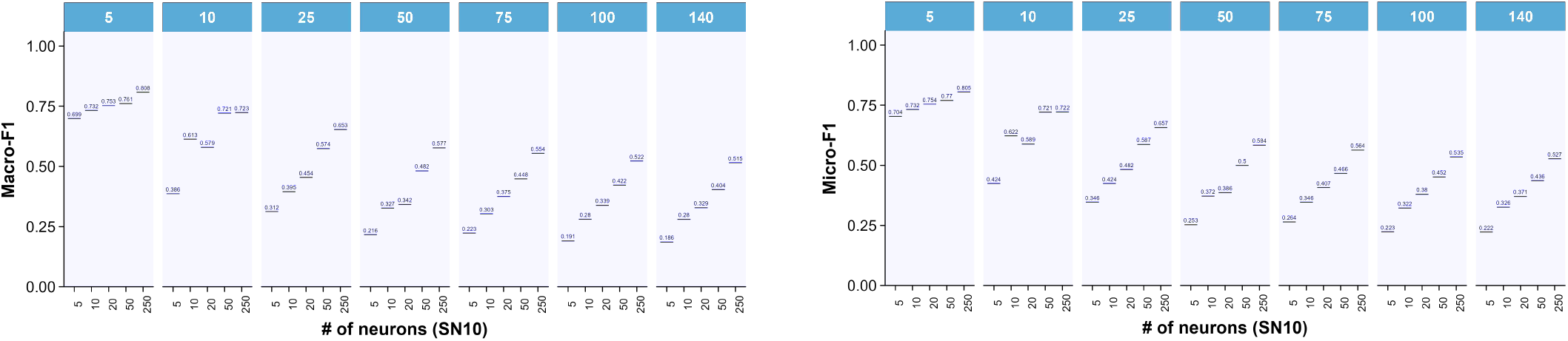
Prediction accuracies of multilabel classification (instead of multiclass in Figure 3), for antibody sequences to bind a set of predefined antigens.

**Supplementary Figure 13 (refers to Figure 4).**
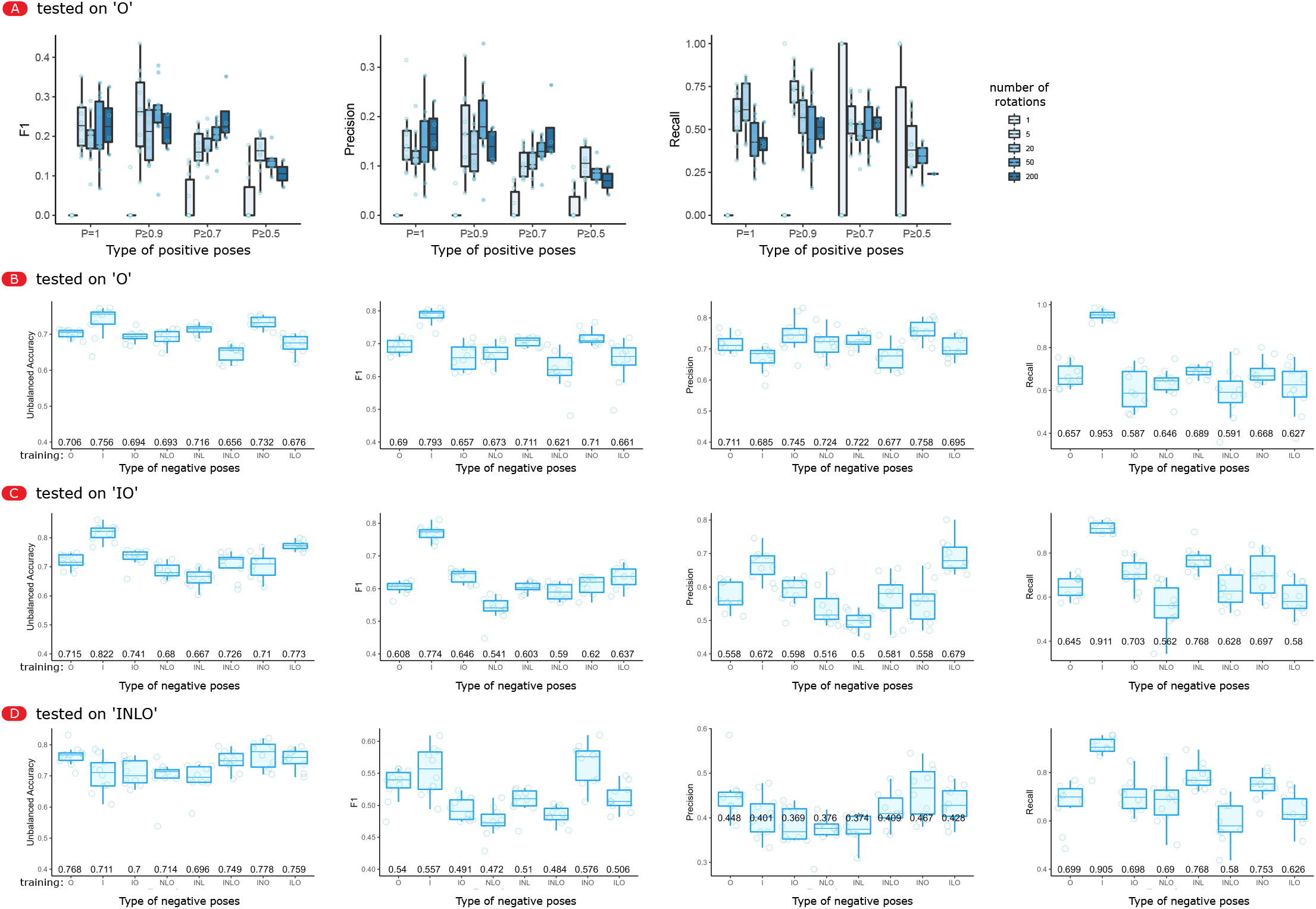
Effect of data augmentation, and definition of positive and negative poses on pose classification prediction performance. (A) Additional metrics of pose classification performance depending on the number of times a pose is replicated by rotation in the dataset (complementing Figure 4C). All trainings are based on the problem “O” (only “other Ab-Ag pairs” as negatives), and are tested on the same dataset design “O”. (B-D) Pose classification prediction performance with different definition of negative examples (complementing Figure 4D). The x axis represents the negatives included in the training dataset. The positives always represent 50% of the instance, while the different negatives are kept within the remaining 50% (see Methods). Since every choice of negative example defines a different problem, it is more fair to compare the train models on the same dataset composition (i.e., the same problem). Therefore, the classification performance is shown on problems “O” (B), “IO” (C), or “INLO” (D).

**Supplementary Figure 14 (refers to Figure 3).**
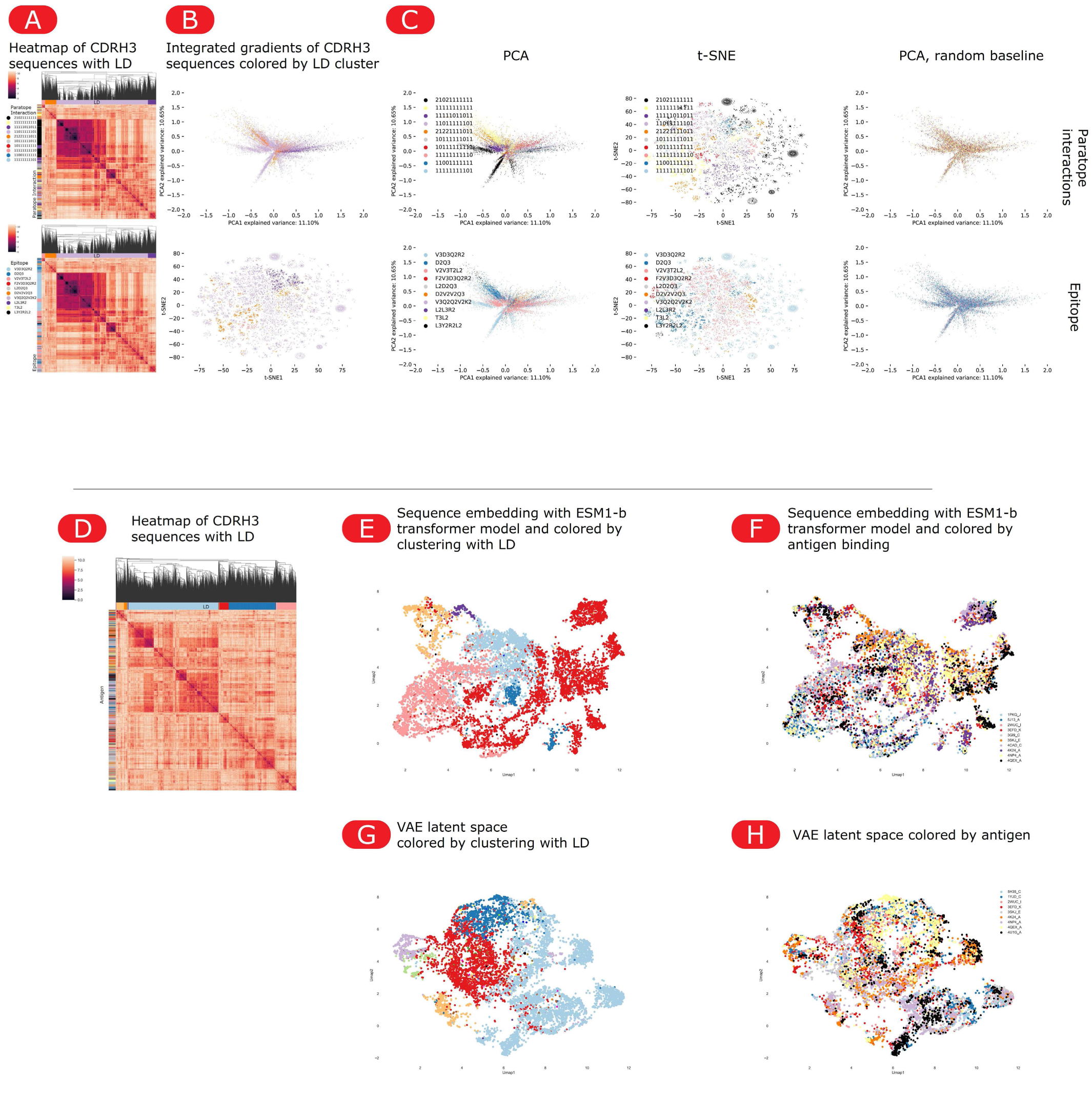
Structural properties of antibody binding can be discriminated using sequence-trained model integrated gradients, but not using sequence similarity, sequence clustering, nor VAE-based latent space representations. (A) Similar sequences do not cluster according to paratope (degree-explicit motif encoding, top) nor epitope encoding (degree-explicit sequence encoding, bottom). 25 000 binder 11-mer sequences to 1ADQ_A from D1 (Supplementary Figure 9) were clustered according to their LD, identifying 10 clusters (horizontal color bar) in the heatmaps. The two heatmaps are identical but showing as vertical color bars the annotation of sequences with either their paratope (top, 10 colors) or epitope (bottom, 10 colors) encoding. (B–C) Integrated gradients can inform whether CDRH3 sequences share a paratope or epitope encoding PCA and t-SNE of integrated gradients of each binder sequences, computed on the SN10 architecture trained with 40 000 training sequences (dataset D1, antigen 1ADQ_A, containing both binders and non-binders Supplementary Figure 9A), colored according to different properties of the sequences: (B) colored according to the 10 clusters generated by agglomerative clustering on the LD between amino acid sequences (see horizontal color bar in panel A); **(C)** colored with associated paratope encoding (top) or epitope encoding (bottom); or colored with shuffled paratope/epitope labels as a comparison (right) for detecting confirmation bias. Sequence coloring by paratope (top) is well separated in the integrated gradients-generated PCA and t-SNE, while agglomerative clustering based on LD sequence similarity could not recover these binding-specific informations. This suggests that taking a new sequence, calculating the integrated gradients of this sequence from the trained model, and integrating the gradients vector into the t-SNE or PCA could be predictive of its binding paratope encoding. In comparison, sequences colored by epitope encoding show more than one cluster for the same color, reminiscent of different modes of binding (paratopes) to the same epitope. (D) Similar CDRH3 sequences bind to different antigens. Sequences from D3 with mono-specific binding sequences to 20 different antigens were clustered according to their similarity (LD), leading to the delineation of 23 clusters (horizontal color bar), and colored according to which antigen they bind (vertical color bar, 20 colors). Clusters of sequence similarity were not predictive for the sequences’ target antigen. (E–H): Neither VAE architecture nor ESM1-b transformer embeddings can isolate CDRH3 sequences by antigen specificity. To save computational time, 233 605 monospecific sequences binding monospecifically 10 different antigens from panel (D) were embedded in the latent space of a VAE (lower row) or through the ESM1-b transformer model ^125^ (upper row) and reduced with UMAP (see Methods). Sequence embeddings were colored according to their LD cluster identified from (D) (left) and by epitope recognition (right). 10 000 embedded sequences are shown for the VAE and 9 000 for the ESM1-b.

**Supplementary Figure 15 (refers to Figure 5).**
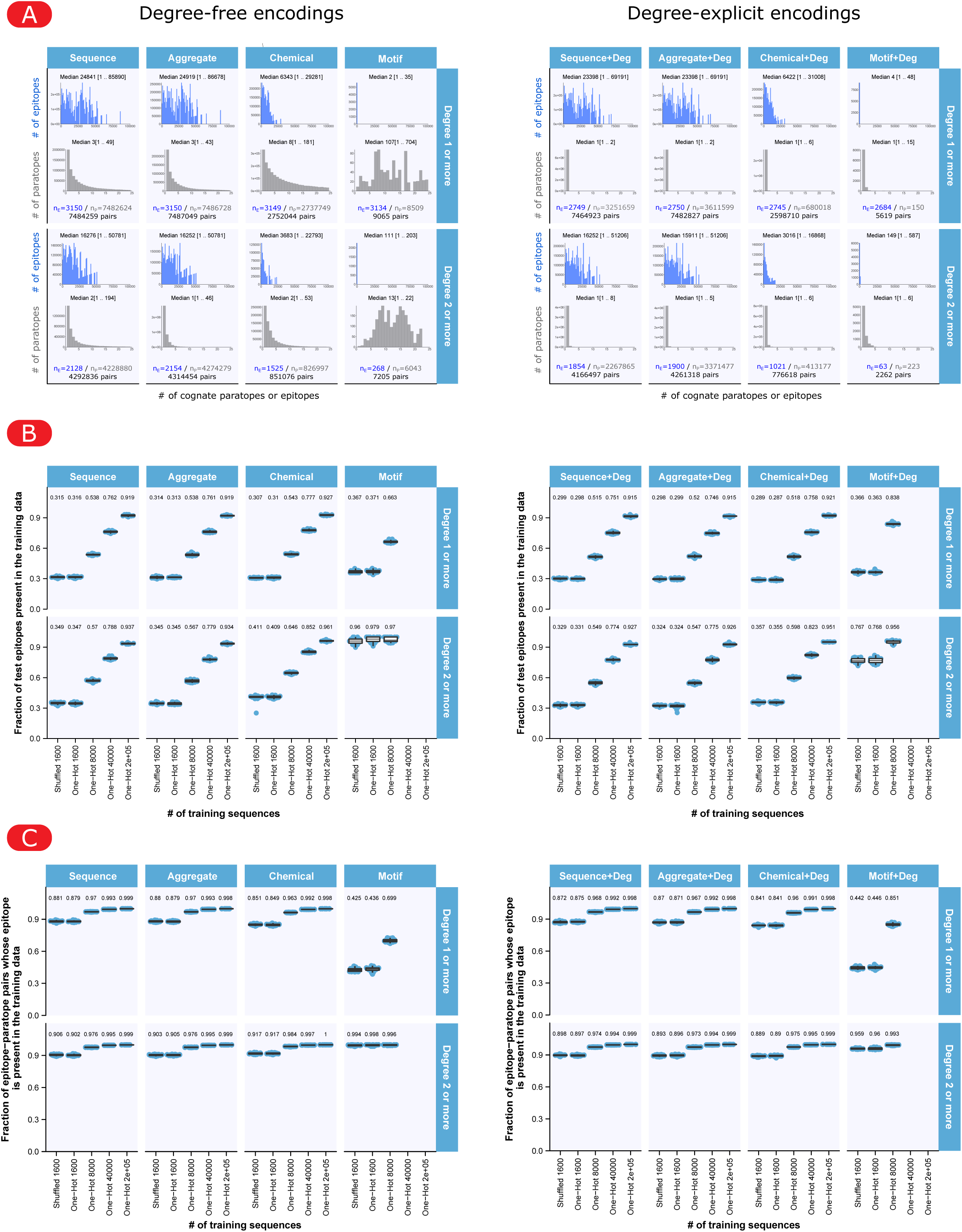
Attributes of the paratope-epitope dataset depending on the encoding. (A) Numbers of cognate paratopes and epitopes with respect to encodings and degree filtering for degree-free (left) and degree-explicit (right). The largest number of pairs was observed for the sequence encoding (4 292 836 and 4 166 497 for degree-free and degree explicit) and the smallest number of pairs was observed for the motif encoding (7 205 and 2 262 for degree-free and degree explicit respectively). (B) Fraction (relative overlap) of test epitopes that were also present in the training data. Broadly the overlap between test and training epitopes increased as a function of the number of training sequences. (C) Fraction of paratope-epitope pairs whose epitopes were present in the training data. As in (B), the overlap increased as a function of the number of training sequences. 10 replicates are shown in each panel/condition.

**Supplementary Figure 16 (refers to Figure 5).**
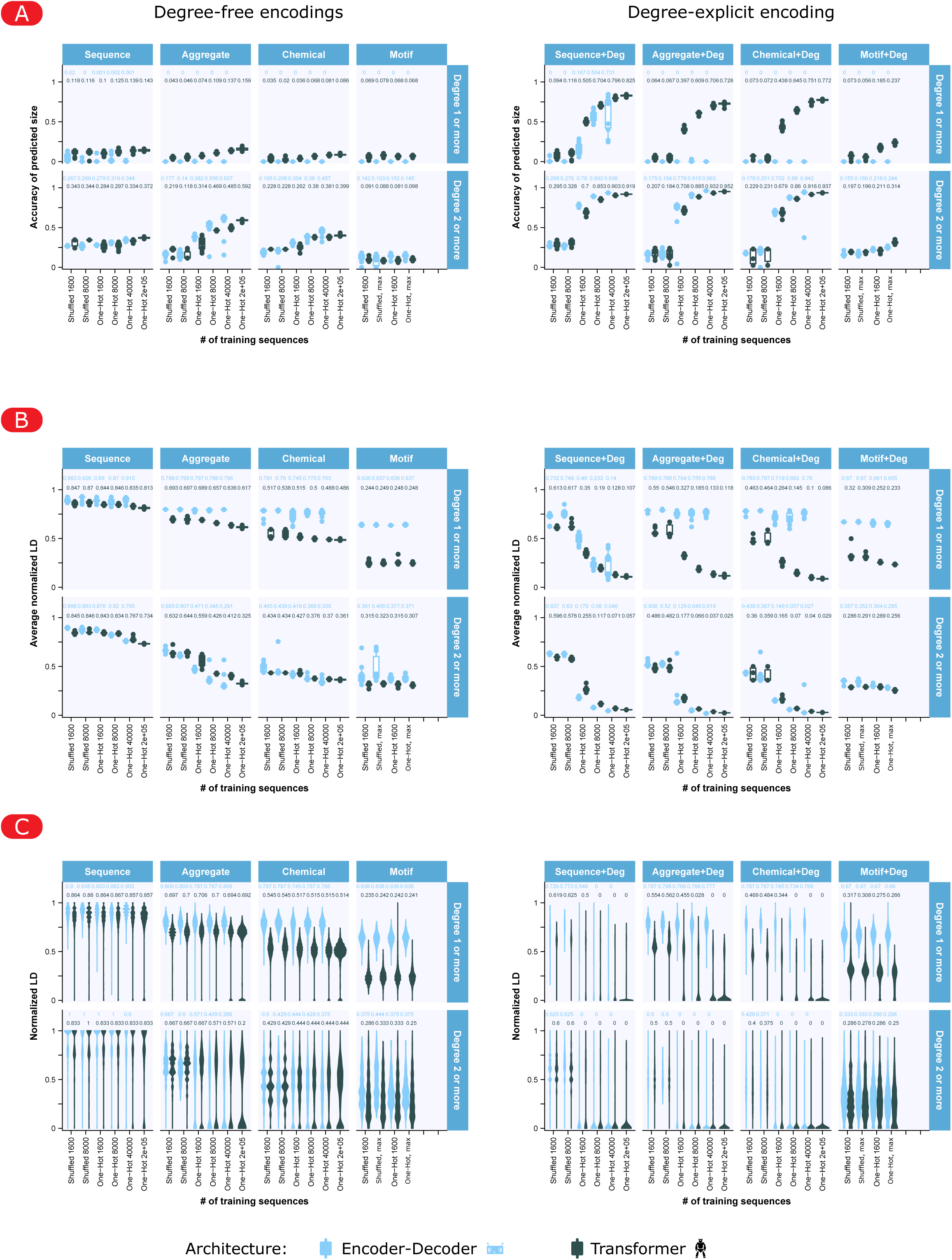
Performance of the transformer and encoder-decoder architectures on the paratope-epitope prediction problem with different encodings. (A–C) Additional accuracy measures for both architectures depending on a large panel of encodings, for degree-free (left) or degree-explicit (right) encodings, with (upper facet) or without (lower facet) residues binding with degree 1. Each point is an independent training, 10 different replicates per condition were performed. Learning on a shuffled dataset (on 1600 and 8000 unique paratope-epitope pairs) is compared with learning on 1 600–40 000 (and 200 000 only for the Transformer architecture) unique paratope-epitope pairs, while all conditions are tested on 100 000 unique pairs when available, or downscaled to reach 80% training and 20% testing (see Methods). The “motifs” encodings only allowed for 2 262 to 9 065 unique paratope-epitopes pairs (see Supplementary Figure 15A), which are referred to as One-Hot (Max) conditions, and higher dataset size is therefore not available). (A) Accuracy of predicting the proper epitope size: fraction of predicted epitopes with the same length as the target epitope. (B) Average Levenshtein distance between predicted and target epitope, normalized to the length of the largest of the two. The shuffled controls show a different reference level for the LD depending on the encoding. Indeed, degree-explicit encodings are longer while degree-filtered ones are shorter (Figure 5A), therefore leading to different possible LD values. (C) Distributions of normalized Levenshtein distance between predicted and target (test set) epitopes. Performance on shuffled pairs was identical with different training sizes (1600 and 8000 pairs).

**Supplementary Figure 17 (refers to Figure 5).**
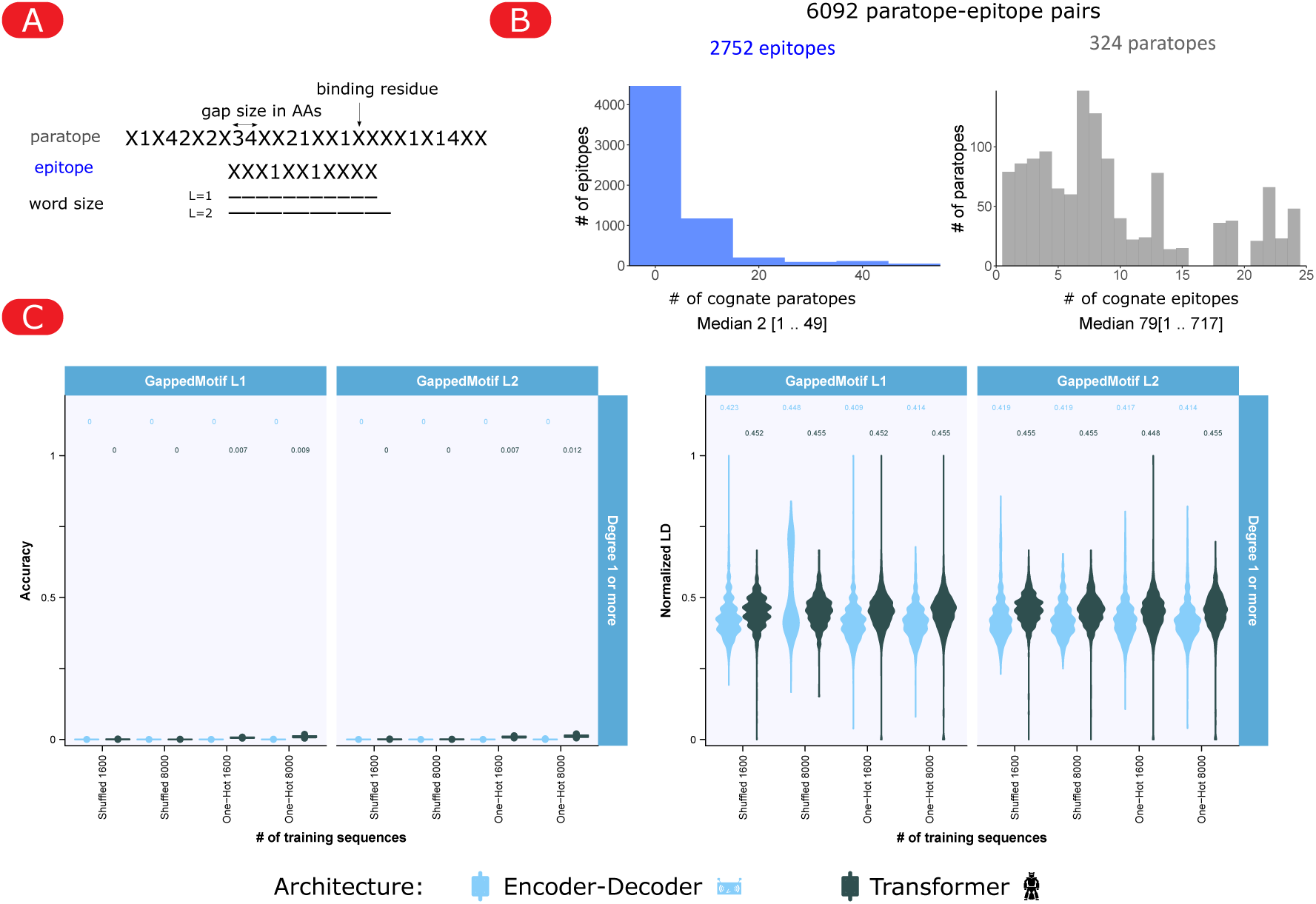
Prediction accuracy of the paratope-epitope prediction problem with the transformer and encoder-decoder architecture, using the “Gapped Motif” encoding. (A) Illustration of the gapped motif encoding. Here, interacting amino acid residues were encoded as the string X and the size (number) of non-interacting residues as integers. (B) The distribution of the number of cognate paratopes (left) and cognate epitope (right). A total of 6092 paratope epitope pairs were observed with 2752 and 324 unique paratope and epitope respectively. (C) Prediction accuracy (left) and LD (right) for encoder-decoder (blue) and transformer (black). Broadly, both architectures yielded poor prediction accuracy (∼0) and no increased LD as compared to the shuffled controls (LD ∼0.4).

**Supplementary Figure 18 (refers to Figure 5).**
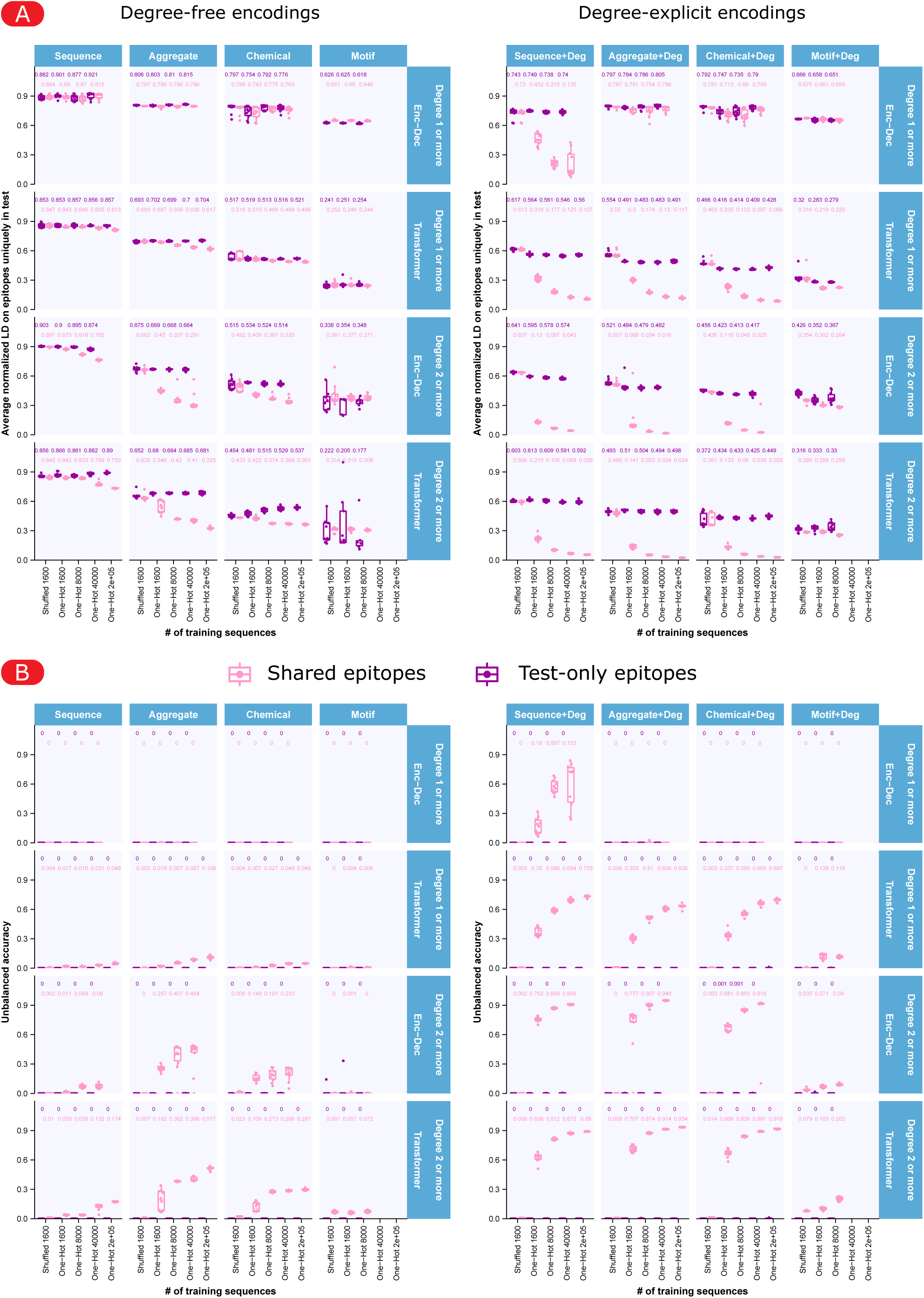
Paratope-epitope prediction accuracy on paratope-epitope pairs whose epitope was present or absent in the test dataset. To examine the prediction accuracy of epitopes that were not present in the training set (test-only epitopes, called “neo epitopes” here), we calculated separately the prediction accuracy (B) and LD (A) of shared (present in both training and test datasets) and test-only epitopes. Broadly the test-only epitopes yielded markedly lower accuracy (B) and larger LD (A) in comparison to shared epitopes across all encodings and degree filtering.

**Supplementary Figure 19 (refers to Figure 5).**
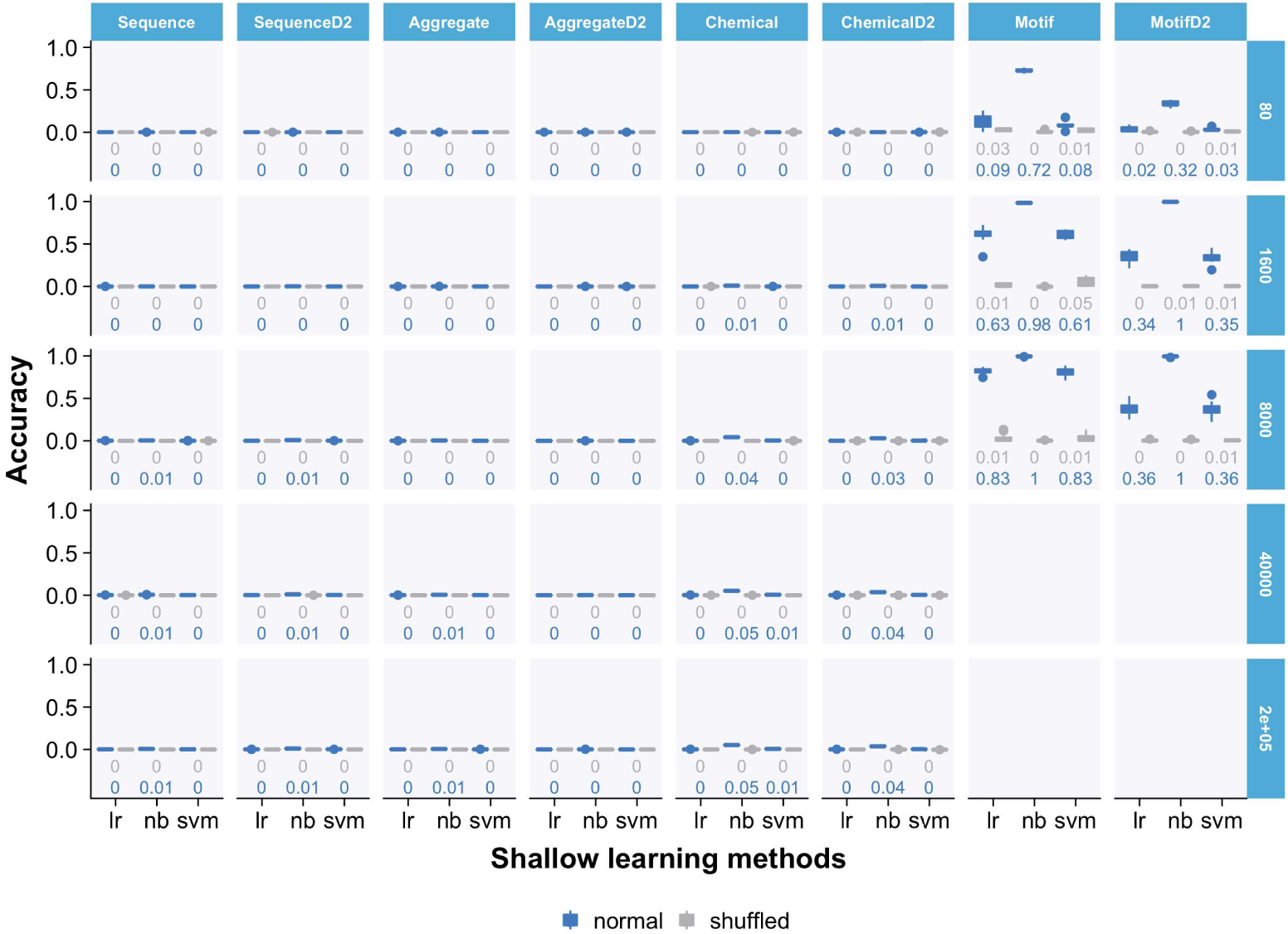
Paratope-epitope prediction with shallow learning methods. As in Figure 5, we examined the performance of shallow learning methods in predicting paratope-epitopes pairs (degree-free encoding). The highest accuracy (shuffled accuracy) values ranged between 0.36–1 (0–0.01) for the model trained on the largest training dataset (n_CDRH3_training_=8 000) and motif encoding (MotifD2). We note that for the motif encoding the maximum n_CDRH3_training_ are 4 495 and 1 809 but we show them in the 8 000 row. In contrast to Figure 5, aggregate and chemical encodings did not improve the prediction accuracy even with increased size of training datasets indicating that shallow learning methods were unable to capture long-range sequence dependencies in the data. Due to the large memory footprint required to train shallow learning models with 200 000 training sequences, the RF classifier was excluded in this analysis. For each model, encoding and n_CDRH3_training_, training and test were replicated 10 times.

**Supplementary Figure 20 (refers to Figure 5).**
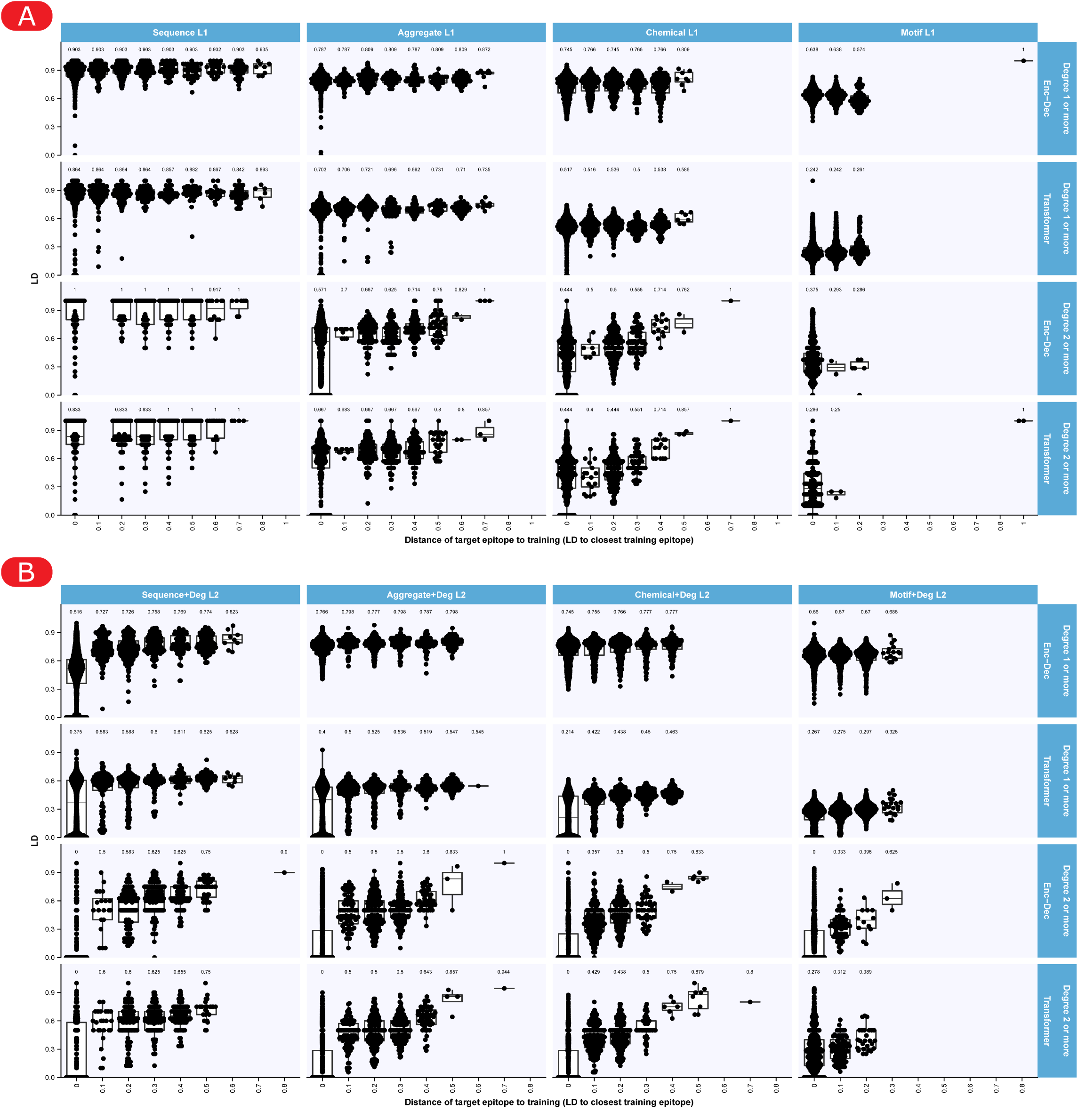
Levenshtein distance of predicted epitopes depending on their similarity to epitopes in the training dataset (x-axis). For each test paratope-epitope pair, the closest epitope in the training dataset (lowest LD) was identified, defining a distance between the epitope and the training dataset (x-axis). The LD of the predicted epitope to the target epitope (y axis) is shown for 5000 pairs in each condition (or all the test pairs for the motifs encoding). 1600 sequences were used for training (enabling to have a high fraction of test-specific epitopes compared to larger training datasets, see Supplementary Figure 15B,C). The fact that accuracy was better in conditions for similar epitopes in certain conditions suggests that the trained model can learn patterns of sequence-wise close epitopes in those conditions (degree-explicit encodings, with degree at least 2 and chemical degree-implicit also with degree at least 2), and that trained models may be applied to unseen epitopes (neoepitopes) provided they are not too different from the training epitopes. (A) Degree-free encodings. (B) Degree-explicit encodings.

**Supplementary Figure 21 (refers to Figure 5).**
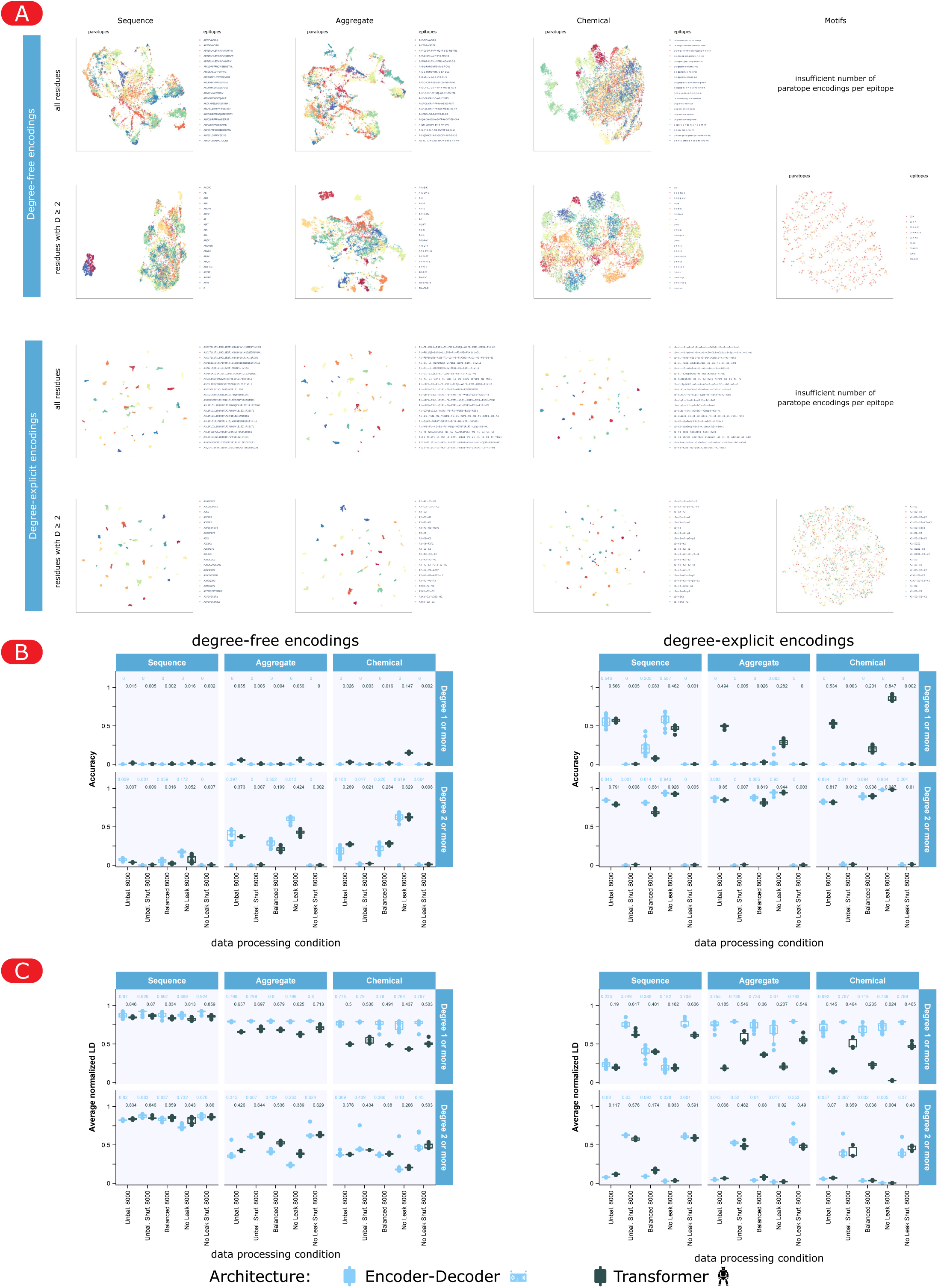
Evaluation of data leakage due to paratope similarity and its impact on the performance of the two NLP architectures. **(A)** To quantify the potential similarity between paratopes sharing the same label (epitope), the paratopes were clustered according to each encoding, and colored by their matching epitope encoding. The clustering was performed based on the LD between paratope encodings using UMAP, and 10 000 paratopes were considered per epitope, to ensure the same amount of datapoints per color (see Methods). Of note, in different encodings, the same LD may have different interpretations: for example, in “aggregate” substitution can mean a difference in one non-interacting residue, while in “sequence” it is always a difference in the interacting one. Motif encodings contained less different paratopes encodings for each epitope. For degree-explicit motif encodings, only 9 epitopes (colors) were matched by 1000 paratopes or more. For degree-free encodings, there were not enough different paratopes encodings per epitope to show clustering. In degree-explicit encodings, one epitope (color) is represented by more than one similarity cluster, possibly representing different modes of binding. Visible clusters with a single color show the risk of data leakage if some sequences from this cluster appear in the training while other sequences from this cluster are in the test, since they are likely very similar. The encodings that led to highest accuracies in Figure 5 seem to generate more visible clusters than those with low accuracy. **(B,C)** Evaluation of paratope-epitope performance after minimizing similarity between train and test data points. **(B)** Paratope-epitope prediction accuracy and **(C)** LD of predicted epitopes compared to the matching epitope of the two NLP architectures after balancing and after removing data leakage. The data processing conditions are shown on the x-axis. 8000 paratope-epitope pairs were used for training, 800 for testing. For each condition, the training and test datasets have the same data distribution. The “unbalanced” condition represents the settings shown in Figure 5 as a control. The “balanced” conditions represent datasets generated with 10 000 paratopes per epitope, before sampling 8000 sequences for the training. The “no leakage” condition represents datasets where paratope similarity clusters (as shown in (A)) were split such that CDRH3 sequences from the same cluster and matching the same epitope are either in the train or test dataset but not both (see Methods), thereby preventing that the trained models learn from similarity, and possibly increasing generalization. In the “no leakage” condition, for each epitope, some clusters were assigned to train and some to test, therefore most of the time the same epitope appears in both training and test (although with dissimilar paratopes). Therefore, to provide a fair comparison, the results of conditions “unbalanced” and “balanced” are only shown for test epitopes that were also present in the training dataset. Altogether, the ranking of encodings according to their accuracy was preserved both after balancing and removing data leakage. Balancing induced a reduced accuracy in some conditions, meaning a part of the unbalanced accuracy was achieved by accurate prediction on larger classes. Removing data leakage generally increased accuracy and decreased the LD of predicted epitopes to their target, compared to both balanced and unbalanced conditions, showing that high accuracy could be achieved despite the reduced similarity between train and test pairs. Therefore, the trained models were able to learn generalizable patterns to some extent.

**Supplementary Figure 22 (refers to Methods).**
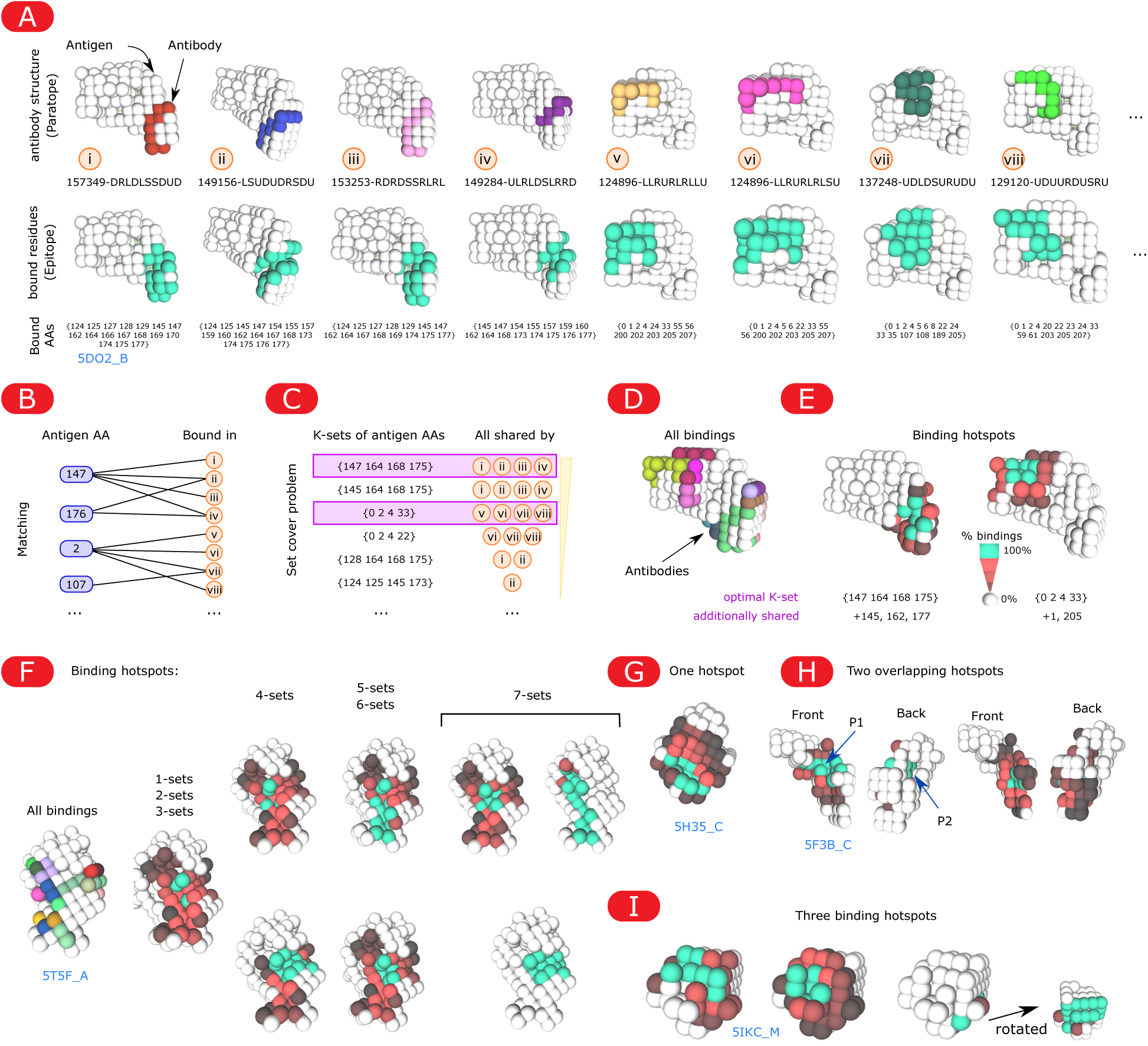
Generation of binding hotspots from a list of antibody binding structures. **(A)** The epitope residues of each binding (i to viii, upper panel) are extracted (turquoise residues, lower panel), provided they bind to an antibody residue of degree at least D (D=1 in this example). The IDs (numbering) of the epitope residues are shown below the images. **(B)** Each antigen residue is matched with all corresponding binding structures where it belongs to the epitope. **(C)** All possible K-sets (example with K=4 here) of residues are associated with the binding structures that share these K residues. The K-set covering the most structures is taken as the first binding hotspot; these structures are removed, and the next K-set that covers the most remaining structures is taken as the second binding hotspot, etc. The two hotspots for 5DO2_B are highlighted in purple. **(D)** Example of all binding structures of the 6.9 million murine CDRH3 sequences around the antigen with PDB ID 5DO2, chain B. Each antibody (11-mer) structure has a different color and potentially overlaps with other antibodies. **(E)** The two binding hotspots associated with all binding structures to antigen 5DO2_B. Each hotspot represents a group of epitopes (/binding structures). Among a group, the turquoise residues are 100% shared, and are K amino acid or more by definition. The level of red is a 3D heatmap on the percent of hotspot associated antibodies that cover this residue (with the degree constraint), and therefore shows the diversity of epitope residues from different binding structures within a binding hotspot. **(F)** Impact of the size of K-sets on the number of hotspots and their shape for antigen 5T5F_A. Smaller values of K cluster binding structures with a too small overlap in their epitopes. Higher values of K lead to duplicate overlapping hotspots, and K=4 was chosen as a good trade-off among 50 inspected antigens (not shown) **(G–I)** Example of the diversity of hotspots for the 6.9 million sequences, with two non-overlapping ones in E, a unique one in G, two non-overlapping hotspots in H, or three overlapping hotspots in I. Although the hotspots tend to form concave topologies, the antigen 5F3B_C in panel H shows a more “immunogenic” pocket (P1) and a “non-immunogenic” pocket (P2), demonstrating that antigen pockets are not necessarily binding hotspots in the Absolut! framework.

**Supplementary Figure 23 (refers to Methods and Figure 5).**
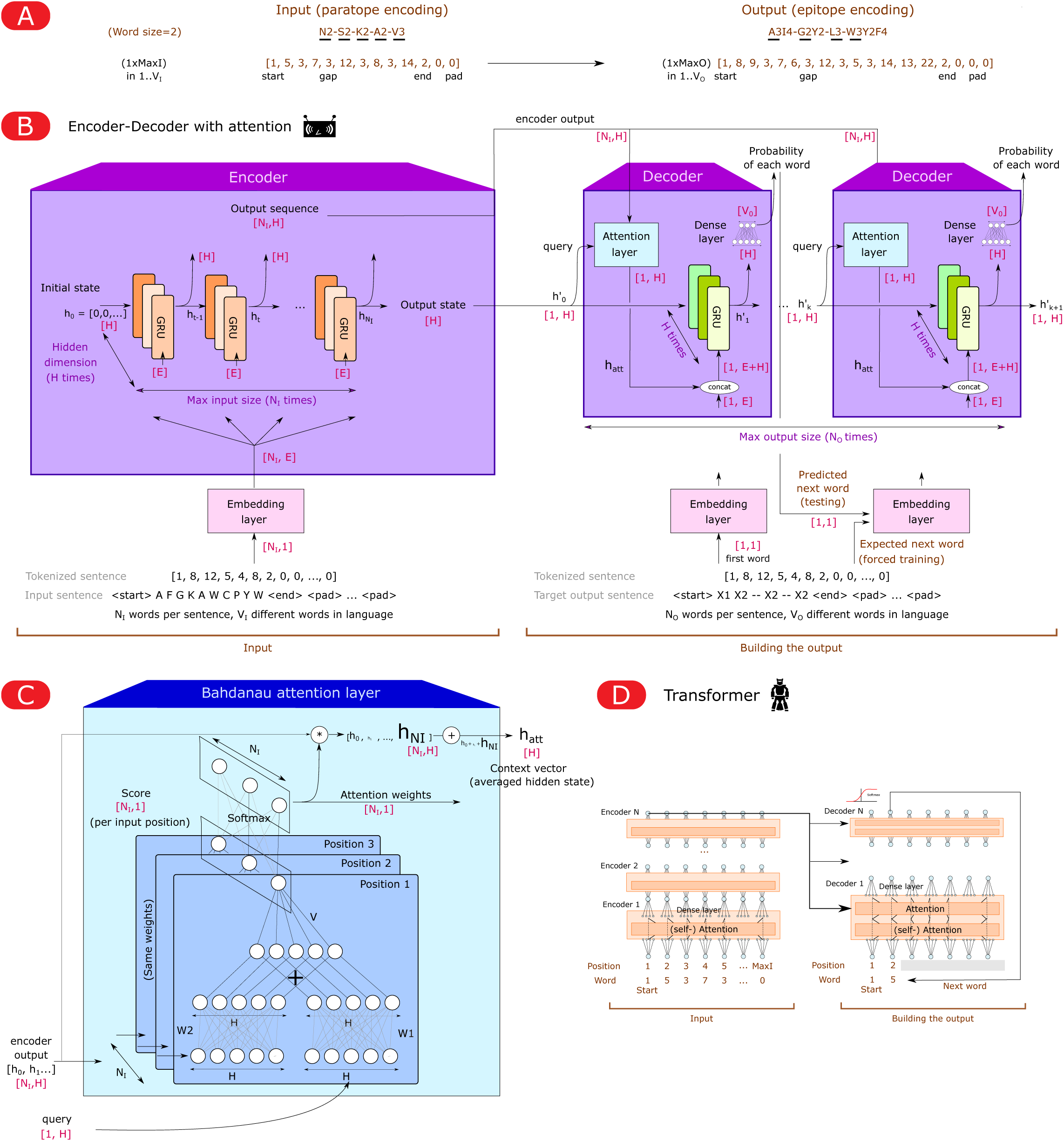
Architecture of the encoder-decoder (NMT) and transformer used for the paratope-epitope prediction task. (A) Input paratope-epitope pairs were encoded according to a range of sequence-structures encodings as described in Figure 5A, start and end positions were encoded by the start and end tokens, and sequences were padded with zero-padding to match the length of the longest sequence in the dataset. Degree-explicit encodings are tokenized by words of size two (with gaps as one word, as shown here), while degree-free encodings are tokenized by words of size one. (B) Illustration of the encoder-decoder architecture (neural machine translation, NMT ^87^). Numerical representation of the input (paratope) sequence was learned by the embedding layer, the output of the embedding layer was passed to the encoder layer (gated recurrent unit, GRU of hidden dimension 512; see Methods) and finally a decoder layer with attention was used to translate the encoded input into an epitope. A Bahdanau attention layer (see panel C) weighs the encoder outputs according to the previous hidden state of the decoder. Numbers in brackets denote tensor dimensions. N_I_: maximum input size; N_O_: maximum output size; E: Embedding dimension (6); H: Hidden layer dimension (512); V_I_/V_O_: number of possible tokens in input/output tokenized language. GRUs with the same color share the same parameters. Word by word, the output of the decoder is transformed into the next token of highest weight (softmax). Forced training denotes when the next target word (not the predicted one) is used to generate the next token. (C) Illustration of the Bahdanau attention layer. (D) Illustration of the transformer architecture ^86^. The encoder layer comprises N stacked-encoders with self attention and the decoder layer comprises N stacked-decoders with attention and self-attention layers (see Methods).

**Supplementary Figure 24 (refers to Figure 3).**
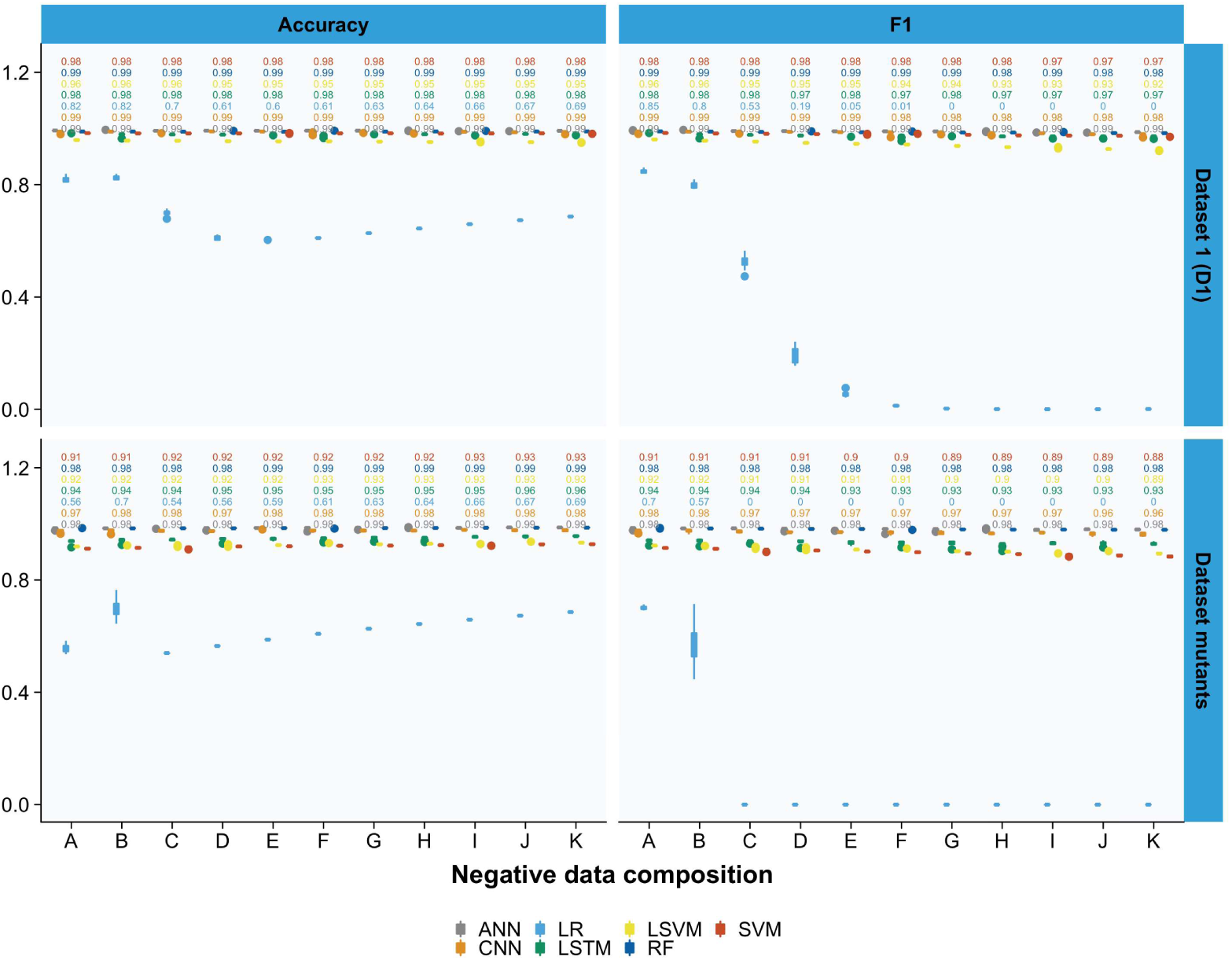
Impact of data imbalance on ML performance. To examine the impact of dataset imbalance (binders/non-binders ratio) on prediction accuracy across different ML methods, we created 11 datasets (A-K) wherein binder to non-binder ratios decreased in the increment of 0.03 from 0.52–0.31 as described in Mason et al ^39^. In agreement with the findings by Mason et al., we found that the accuracy and F1 score decreased as the imbalance in the datasets increased (non-binder>binder) ^39^. The overall performance ranking of Methods reported in Mason et al. was also preserved. To be consistent with other results in the manuscript, the type of hyperparameters in the RF and LR architectures were kept the same as in other tasks, (i.e., not taken from Mason et al.). The low performance of the RF architecture suggests that RF is particularly sensitive to training data class imbalance.

**Supplementary Table 1 (refers to Figure 2A).**
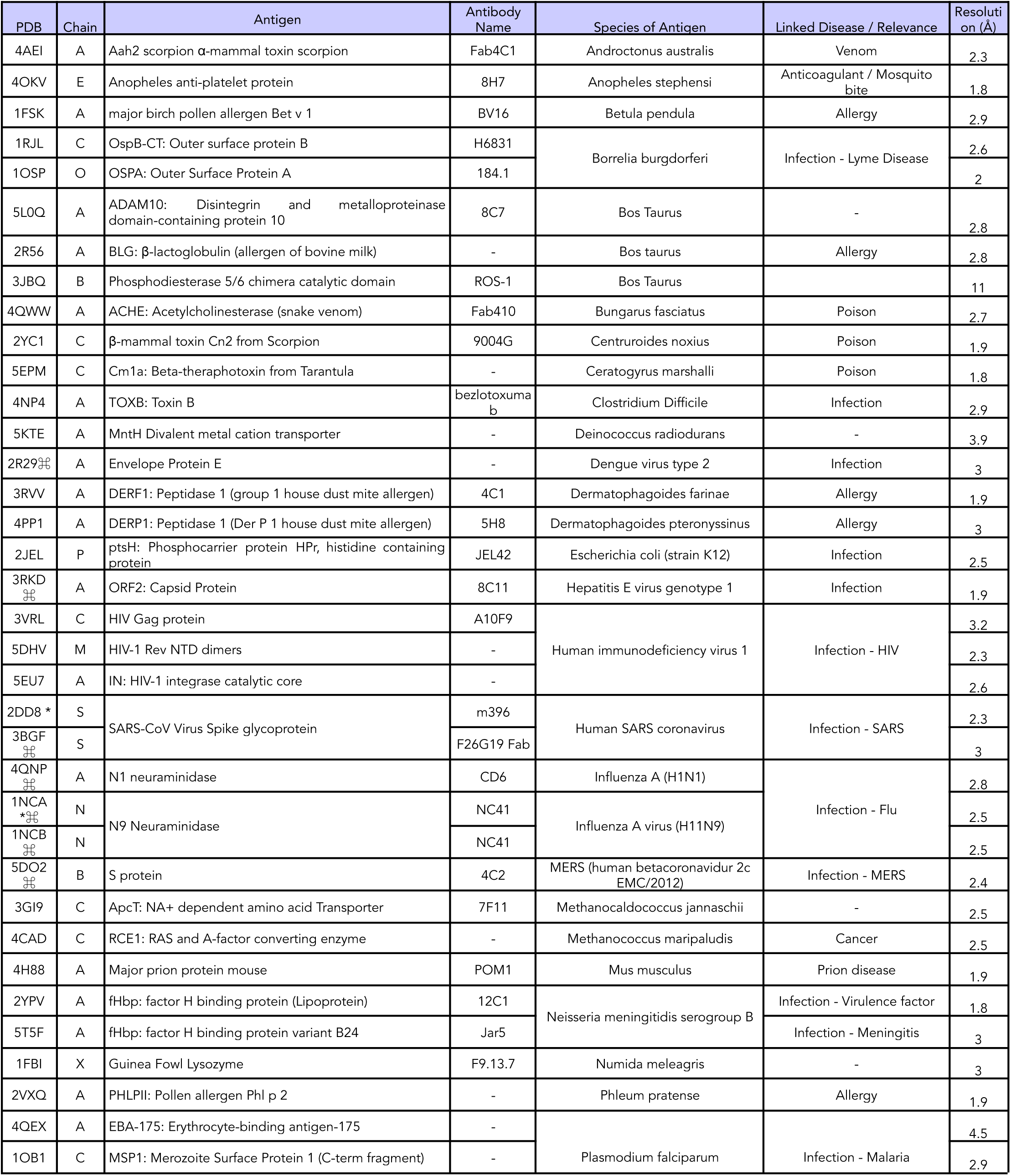
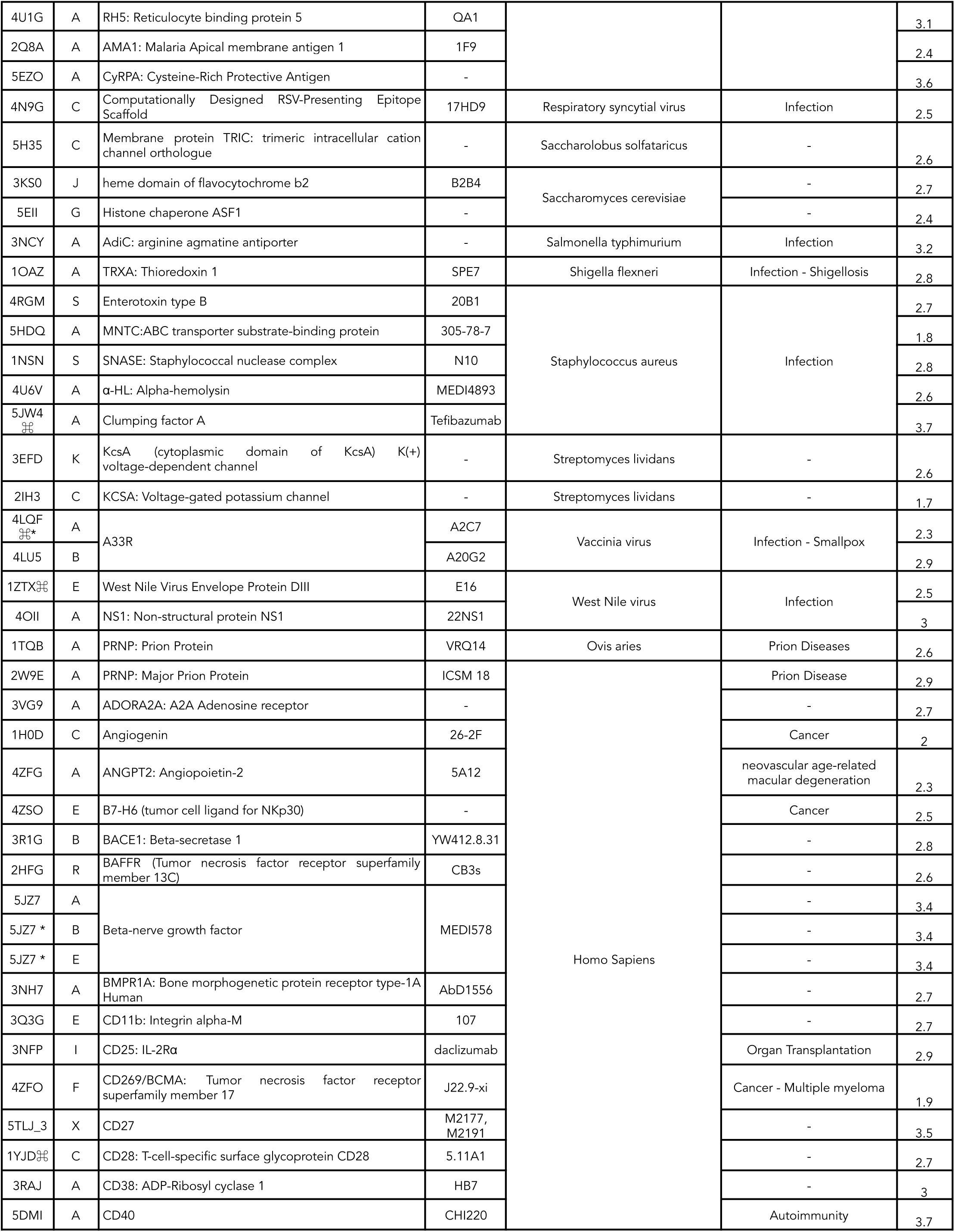
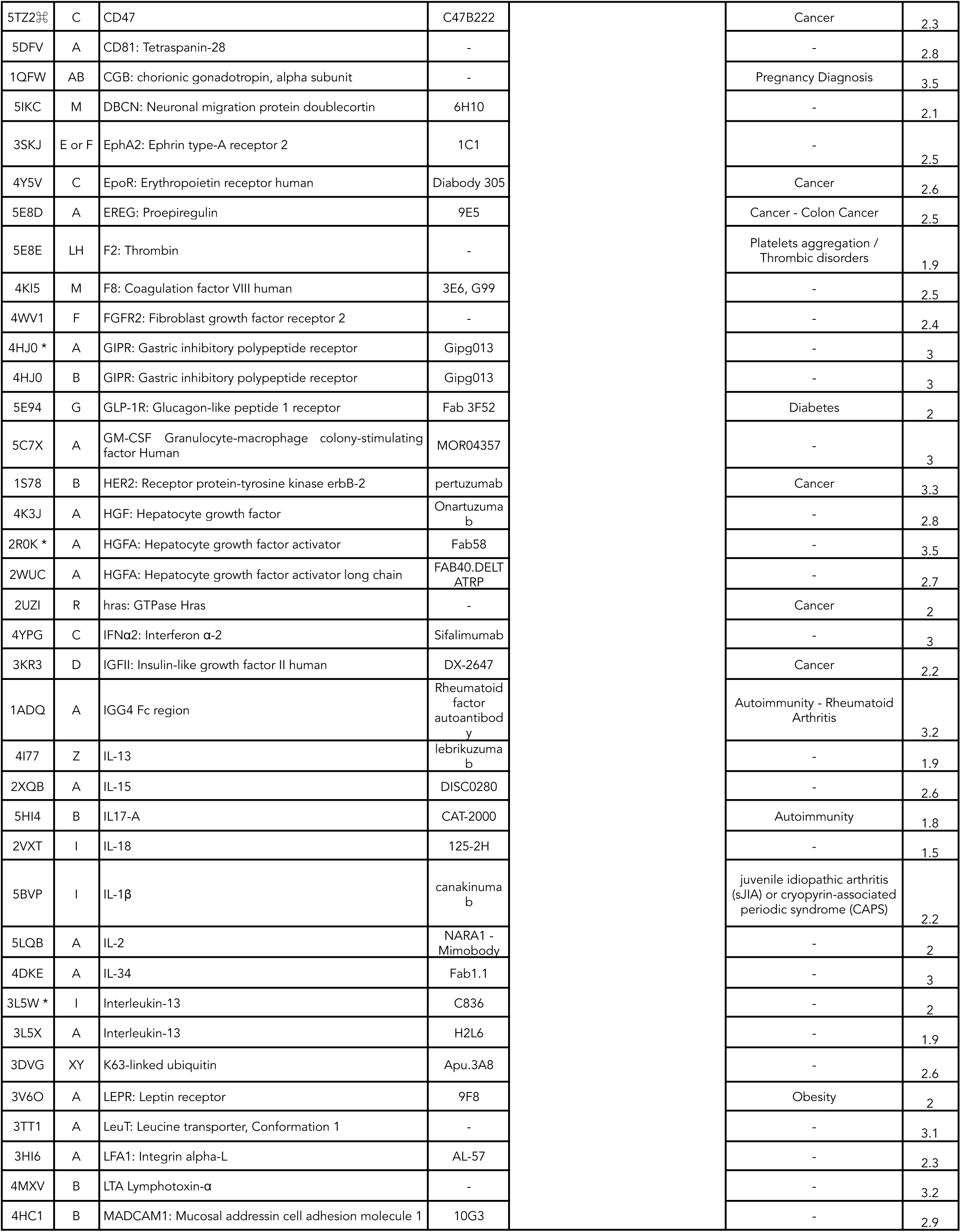
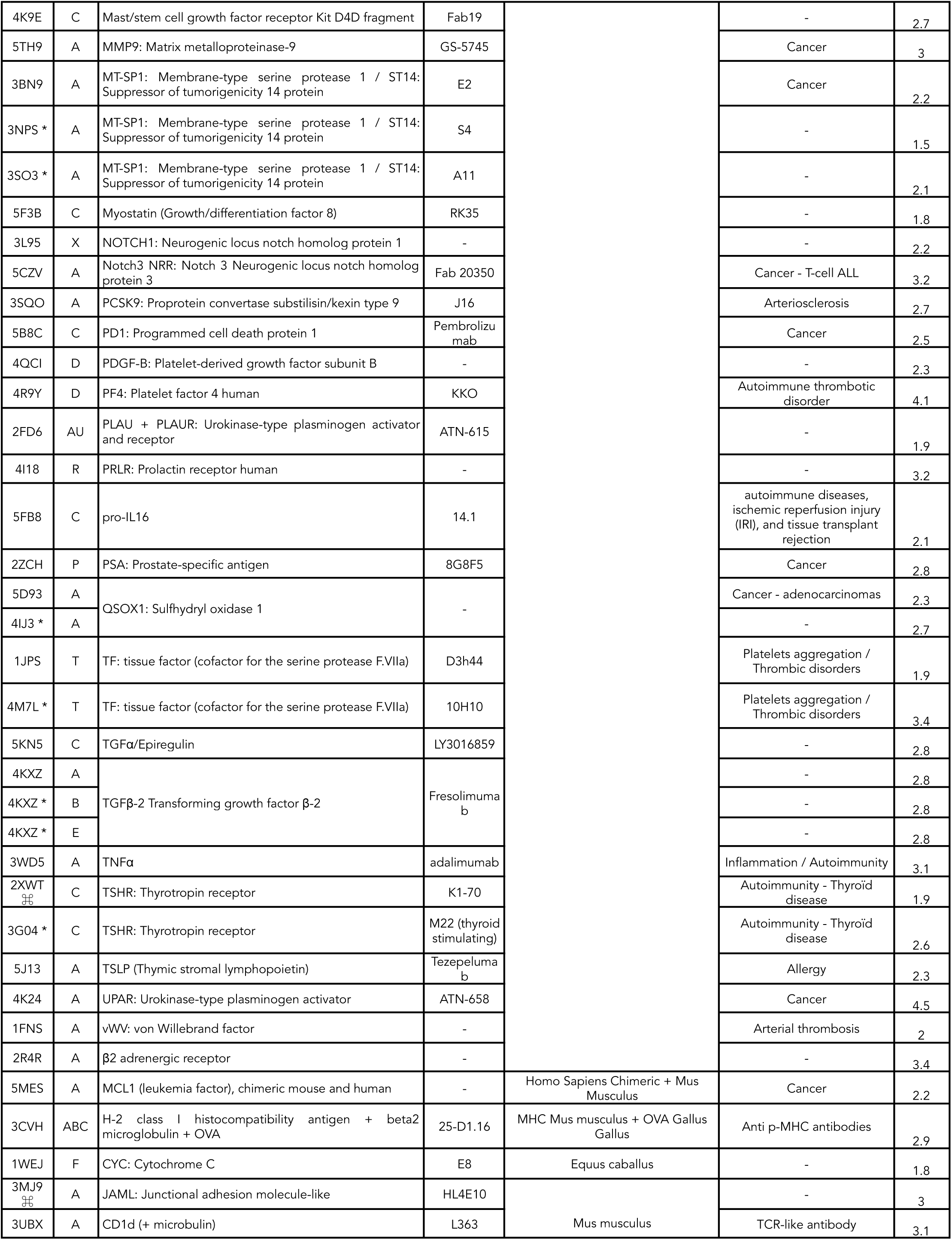
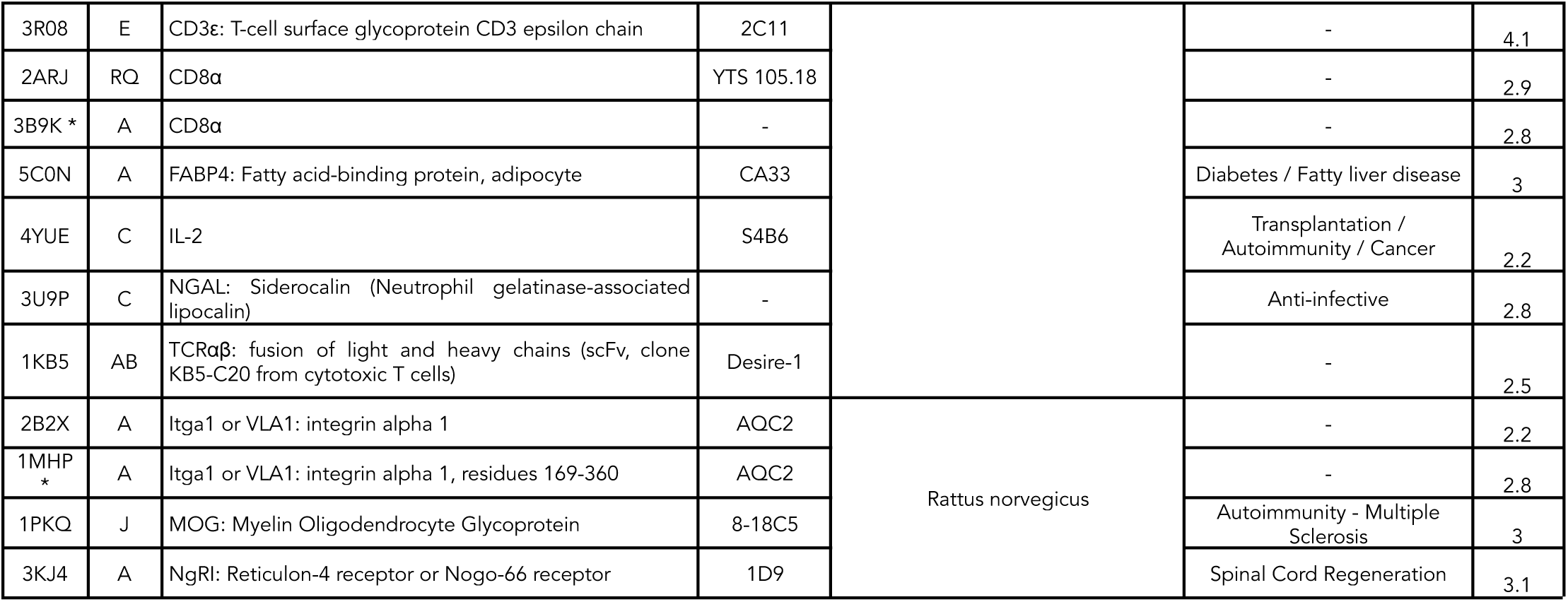
List of the 159 antigens provided in the Absolut! database. The PDB ID, the discretized chain, and information on the antibody-antigen crystal structure are provided. (*) these antigens with more than one discretization were not considered for ML tasks involving multiple antigens, to avoid considering the same antigen twice, leading to 142 remaining antigens for multi-class antigen ML tasks (see Figure 3, Supplementary Figure 9). (⌘) antigens containing glycans.

